# Developmental and behavioral phenotypes in a new mouse model of DDX3X syndrome

**DOI:** 10.1101/2021.01.22.427482

**Authors:** Andrea Boitnott, Dévina C Ung, Marta Garcia-Forn, Kristi Niblo, Danielle Mendonca, Michael Flores, Sylvia Maxwell, Jacob Ellegood, Lily R Qiu, Dorothy E Grice, Jason P Lerch, Mladen-Roko Rasin, Joseph D Buxbaum, Elodie Drapeau, Silvia De Rubeis

## Abstract

**Background:** Mutations in the X-linked gene *DDX3X* account for ~2% of intellectual disability in females, often co-morbid with behavioral problems, motor deficits, and brain malformations. *DDX3X* encodes an RNA helicase with emerging functions in corticogenesis and synaptogenesis.

**Methods:** We generated a *Ddx3x* haploinsufficient mouse (*Ddx3x*^+/−^) with construct validity for *DDX3X* loss-of-function mutations. We used standardized batteries to assess developmental milestones and adult behaviors, as well as magnetic resonance imaging and immunostaining of cortical projection neurons to capture early postnatal changes in brain development.

**Results:** *Ddx3x*^+/−^ mice show physical, sensory, and motor delays that evolve into behavioral anomalies in adulthood, including hyperactivity, anxiety-like behaviors, cognitive impairments, and motor deficits. Motor function further declines with age. These behavioral changes are associated with a reduction in brain volume, with some regions (e.g., cortex and amygdala) disproportionally affected. Cortical thinning is accompanied by defective cortical lamination, indicating that *Ddx3x* regulates the balance of glutamatergic neurons in the developing cortex.

**Conclusions:** These data shed new light on the developmental mechanisms driving DDX3X syndrome and support face validity of this novel pre-clinical mouse model.

## INTRODUCTION

DDX3X syndrome is a rare neurodevelopmental disorder (NDD) predominantly affecting females. The condition is estimated to account for ~2% of intellectual disability/developmental delay (ID/DD) in the female population (1, 2). In addition to ID/DD, the syndrome presents with a constellation of co-morbidities. Affected individuals can have low body weight, hypotonia and/or hypertonia with spasticity, microcephaly, seizures, movement disorders, gait anomalies, and behavioral problems. Brain malformations (e.g., agenesis of the corpus callosum and polymicrogyria) are also common (1, 3, 4).

DDX3X syndrome is due to mutations in the RNA helicase *DDX3X* (1, 2). Other genes in the same superfamily (DExD/H-box) are also emerging as associated to ID/DD (5–7). DDX3X facilitates the ATP-dependent unwinding of RNA secondary structures. Therefore, it is broadly implicated in RNA metabolism, especially mRNA translation (3, 8–10). *DDX3X* is located on the X chromosome but it escapes X chromosome inactivation (11, 12). As a result, females express two alleles, while males can only express one. The *DDX3X* paralog on the Y chromosome (*DDX3Y*) does not appear to compensate for this dosage imbalance as it has been shown to be translated only in spermatocytes (13). In line with this expression pattern, *DDX3Y* deletions cause subfertility or infertility, not adverse neurodevelopmental outcomes (14).

Females with DDX3X syndrome carry *de novo* mutations, either loss-of-function mutations plausibly leading to haploinsufficiency or missense/in-frame mutations (1–3). Genotype-phenotype analyses have shown that missense/in-frame mutations are associated with more severe phenotypes than loss-of-function mutations. For example, polymicrogyria has not been detected in individuals with loss-of-function mutations but only in a subset of females with missense/in-frame mutations, and this group is more delayed developmentally (3). Only a few affected males—all carrying missense mutations—have been identified (1, 3, 15, 16). These males often inherit the mutations from their mothers, who appear asymptomatic. This observation, alongside functional data in zebrafish (1, 16), suggests that the mutations found so far in males act as hypomorphic alleles. More disruptive mutations in *DDX3X* are postulated to cause prenatal lethality in males.

The developmental mechanisms underlying DDX3X syndrome are just beginning to emerge. Evidence from mouse studies indicate that *Ddx3x* is required for embryogenesis (17), hindbrain development (18), corticogenesis (3), and synaptogenesis (10). *Ddx3x* germline knockout in the mouse results in placentation defects, followed by early lethality at embryonic day E6.5 (17). Restricting the knockout to embryonic tissues nevertheless alters embryogenesis: *Ddx3x* null male embryos show multiple anomalies (e.g., cardiac malformations, defective neural tube closure, and underdeveloped brain) and die at ~E11.5 (17). *Ddx3x* heterozygous females were described as grossly normal, but were not characterized from developmental or behavioral perspectives (17). Mice with a *Ddx3x* conditional deletion in neural stem cells at E9.5 survive but show altered brain growth, especially at the expense of the cerebellum, accompanied by severe ataxia and seizures (18). In addition, *Ddx3x* regulates cortical neurogenesis: knockdown of ~25% *Ddx3x* in cortical neuronal progenitors at E14.5 impacts their differentiation into neurons (3). Finally, postnatal *Ddx3x* knockdown disrupts neurite outgrowth, and dendritic spine formation and maturation (10).

While *Ddx3x* has been found to regulate brain development (3, 18), its role in postnatal and adult behavior is unknown. Also, while studies have showed that *Ddx3x* is needed for corticogenesis (3) and synaptogenesis (10), the lack of analyses in an animal model has limited our ability to relate these cellular functions to behavior. Here, we present a novel *Ddx3x* haploinsufficient mouse (*Ddx3x*^+/−^) with construct validity for *DDX3X* loss-of-function mutations found in females. We show evidence for awry developmental trajectories that precede adult behavioral abnormalities, including anxiety-like behavior, hyperactivity, and memory impairments. Motor deficits are prominent in adulthood and worsen with ageing. These developmental and behavioral phenotypes are accompanied by reduced brain volume consequent to the overall growth delay and alterations in the cytoarchitecture of the developing cortex.

## METHODS AND MATERIALS

### Mice

All animal procedures were approved by the Institutional Animal Care and Use Committee of the Icahn School of Medicine at Mount Sinai. The *Ddx3x^flox^* mouse line was generated at Ozgene by introducing two loxP sites flanking exon 2 of mouse *Ddx3x* (OTTMUSG00000017078) in C57BL/6J embryonic stem cells. To generate the *Ddx3x* knockout (KO, -) allele, *Ddx3x^flox^*^/flox^ females were crossed with B6.Cg-Edil3^Tg(Sox2-cre)1Amc^/J males (Sox2-Cre) (19) (The Jackson Laboratory, stock number #008454). The colony was maintained in a room on a 12/12 h light/dark cycle, with lights on at 7 A.M. at a constant temperature of 21-22°C and 55% humidity. Standard rodent chow and potable water were available *ad libitum.* Animals were socially housed, with 3-5 mice per cage. Mice were weaned at postnatal day (P) 21. The colony was maintained on a C57BL/6J background. To generate the *Ddx3x*^+/−^, *Ddx3x*^+/+^ and *Ddx3x*^+/y^ mice in this study, homozygous *Ddx3x^flox^*^/flox^ were crossed with heterozygous Sox2-Cre/+ males. **Table S1-2** report the cohorts employed for all behavioral analyses. Genotyping is described in the **Supplemental Note**.

### Immunoblotting

P21 *Ddx3x*^+/y^, *Ddx3x*^+/+^, and *Ddx3x*^+/−^ mice were euthanized by decapitation. Total cortices or synaptosomes (see **Supplemental Note**) Following rapid dissection on ice, cortices were homogenized in ice-cold RIPA™ lysis buffer (ThermoFisher) containing protease, phosphatase, and RNase™ ribonuclease inhibitors (ThermoFisher), incubated on ice for 5 min, and centrifuged at 12,000g for 8 min at 4°C. The supernatants containing total cortex protein lysate were quantified using the Pierce™ BCA Protein Assay kit (ThermoFisher). Immunoblotting was performed using standard protocols with 10μg of protein loaded onto SDS-PAGE pre-cast gels (Bio-Rad). Antibodies used were anti-DDX3X (rabbit polyclonal, 1:1000, Millipore cat. #09-860) and anti-GAPDH (loading control, mouse monoclonal, 1:20,000, Millipore cat. #CB1001). Secondary antibodies were HRP-conjugated anti-rabbit and anti-mouse antibodies (1:5,000, Jackson ImmunoResearch Laboratories). Immunoblots were developed using ECL or West Femto (Thermo Scientific), visualized using Syngene G:Box imaging system, and quantified by densitometry in ImageJ.

### Testing of developmental milestones

Testing was performed as reported previously (20), using a battery of tests adapted from the Fox scale (21, 22). Four independent cohorts were used for the testing of developmental milestones, as indicated in **Table S1**. To control for litter size effects, litters were homogenized and limited to 6 pups per dam by adding excess pups to smaller litters on P1, whenever possible. Pups were identified by hind-paw tattoo using a nontoxic animal tattoo ink (Animal Identification & Marking Systems Inc) inserted subcutaneously through a 30-gauge hypodermic needle tip into the center of the paw. Testing occurred at the same time of day, every day from P1 to 21 and weight continued to be taken until P27. Individual pups were removed from the litter and placed on cotton pads under a heating lamp to maintain a temperature of 28°C throughout the testing. All tasks were conducted quickly and capped at 30 seconds to minimize maternal separation. The animals used for testing of developmental milestones were not employed for adult testing, to avoid confounding effects from maternal separation. All three genotypes were tested on the same day in randomized order by an experimenter blind to the animals’ genotype. All data were scored and analyzed blind to the genotype. Details for each test are described in the **Supplemental Note**.

### Gait analysis

Testing was performed as reported previously (20). Briefly, the paws of the mice were coated in non-toxic and water-washable dyes (back paws in blue and front paws in red. Mice were then placed over an absorbent paper on a runway and encouraged to walk in a straight line. Measures were taken as indicated in **Fig. S5**.

### Behavioral testing of adult and aged mice

Four independent cohorts of naïve mice were used for adult behavioral testing and two independent cohorts were used for behavioral testing in ageing (**Table S2**). For adult cohorts, testing started at 10 weeks +/− one week. For aged cohorts, testing started at 1 year old. Behavioral testing was conducted during the light phase in sound-attenuated rooms. A week prior testing, we recorded physical appearance and spontaneous activity, as detailed below. Testing was conducted in the order indicated in **Table S2.** Mice were habituated to the testing room for 30 min prior to testing. All surfaces and equipment were cleaned with 70% ethanol between trials and nitrile gloves were used. The two genotypes were tested on the same day in randomized order by an experimenter blind to genotype. All data were scored and analyzed blind to the genotype. Adult and ageing animals were handled daily for one week prior to behavioral testing to assess general health, psychical appearance, and spontaneous activity, as detailed in the **Supplemental Note**. Details for each test are described in the **Supplemental Note**.

### Magnetic resonance imaging (MRI)

P3 *Ddx3x*^+/+^ and *Ddx3x*^+/−^ mice were euthanized by decapitation, and tail biopsies isolated for genotyping. The whole heads underwent fixation suing 4% paraformaldehyde (PFA) and 2mM ProHance for 24 hrs, then transferred to 0.1M Phosphate-buffered saline (PBS) + 2mM ProHance + 0.02% sodium azide for at least 1 month prior to scanning. Images were acquired on a 7 Tesla MRI scanner (Agilent Inc.) (23, 24). The contrast required for registration and assessment of volume at early postnatal ages is not compatible with our typical T2-weighted imaging sequence. Therefore, diffusion weighted imaging was performed to enhance the contrast between white and gray matter to aid in the registration. The diffusion sequence used an in-house custom built 16-coil solenoid array to acquire images from 16 brains in parallel (25). The diffusion sequence used was a 3D diffusion-weighted FSE, with TR= 270 ms, echo train length = 6, first TE = 30 ms, TE = 10 ms for the remaining 5 echoes, one average, FOV = 25 mm × 14 mm × 14 mm, and a matrix size of 450 × 250 × 250, which yielded an image with 56 μm isotropic voxels. One b=0 s/mm^2^ image was acquired and 6 high b-value (b = 2147 s/mm^2^). The total imaging time was ~14 hours. To visualize and compare the mouse brains, the 6 high b-value images were averaged together to make a high-contrast image necessary for accurate registration (see **Fig. 7A**). Then these images were linearly (6 parameter followed by a 12 parameter) and nonlinearly registered together. All scans were then resampled with the appropriate transform and averaged to create a population atlas representing the average anatomy of the study sample. All registrations were performed using a combination of the mni_autoreg tools (26) and ANTS (27). The result of the registration was to have all scans deformed into exact alignment with each other in an unbiased fashion. For the volume measurements, this allowed for the analysis of the deformations needed to take each individual mouse’s anatomy into this final atlas space, the goal being to model how the deformation fields relate to genotype (24, 28). The Jacobian determinants of the deformation fields are then calculated as measures of volume at each voxel. These measurements were examined on a regional and a voxel-wise basis in order to localize the differences found within regions or across the brain. Regional volume differences were calculated by warping a pre-existing classified MRI atlas onto the population atlas, which allows for the volume of 62 segmented structures encompassing cortical lobes, large white matter structures (i.e. corpus callosum), cerebellum, and brainstem (29). Multiple comparisons were controlled for by using the False Discovery Rate (FDR) (30).

### Immunostaining

P3 *Ddx3x*^+/+^ and *Ddx3x*^+/−^ mice were euthanized by decapitation, and tail biopsies isolated for genotyping. Brains were fixed in 4% PFA in PBS overnight at 4°C, and then cryopreserved in 30% (w/v) sucrose, 0.05% sodium azide and 100mM glycine in PBS. Brains were embedded in Tissue-Tek® O.C.T. Compound (VWR) and sectioned using a Leica CM1860 cryostat. Six to eight coronal sections (40μm thickness) were selected from each mouse for immunostaining. The sections were washed in PBS + 0.05% Triton X-100 Thermo Scientific) and blocked in PBS + 0.5% Triton X-100 + 5% donkey serum (Sigma) for 1 hr at room temperature. Primary antibodies employed include rabbit monoclonal anti-SATB2, Abcam, #ab51502, 1:400; rat monoclonal anti-CTIP2/BCL11B, Millipore, #MABE1045, 1:500; and, goat polyclonal anti-BRN1/POU3F3, Novus Biosystems, #NBP1-49872, 1:200). Primary antibodies were incubated overnight at 4°C in the dark. After washing in PBS + 0.05% Triton X-100, sections were incubated for 2 hrs at room temperature with fluorescent secondary antibodies (anti-mouse Alexa Fluor 594, Invitrogen, #R37115; anti-mouse Alexa Fluor 488, Invitrogen, #A21202; anti-rat Alexa Fluor 568;all at 1:200 dilution). Sections were mounted using antifade mounting medium with DAPI (Vectashield, #H-1200).

### Fluorescence and confocal imaging and processing

Images of whole cortices were acquired using an EVOS M7000 microscope (Invitrogen) with 4X or 10X objectives. Representative images shown throughout the manuscript were acquired using a Leica TCS SP8 laser scanning confocal microscope (Leica Microsystems Heidelberg GmbH, Manheim, Germany) coupled to a Leica DMi8 inverted microscope at 10x magnification. Argon and 561nm lasers and HyD detectors were used. Images were acquired at 200 Hz and 2048×2048 pixels. Coronal sections of the left and right cortex were reconstructed from multiple individual images using GIMP. This software was also used for binning M1 and M2. Using a mouse brain atlas, M1 and M2 were located in the sections. To normalize all bins across all samples and sections, the beginning of the M1 bin started 400px from the peak of the cingulum bundle and spanned 600px wide. The bin for M2 was centered on the peak of the cingulum bundle with a width of 400px. Each region was binned evenly, 4 across and 10 down.

### Statistical analyses

All statistics and plots were generated using custom R scripts. Statistical tests, number of animals in each experiment, and significance are indicated in each figure legend. Data are shonw as mean±SEM. Outliers are defined as data points below Q1-1.5xIQR or above Q3+1.5xIQR, where Q1 is the first quartile, Q3 is the third quartile, and the IQR is the interquartile range. Outliers, shown in plots with the ⊗ symbol, were removed from the calculations of the mean and SEM and for the statistical tests. Repeated measure ANOVA was used for repeated observations. For comparisons between two groups, Shapiro–Wilk test was used to assess normality, followed by Student’s t-test for normally distributed data or Wilcoxon signed-rank test for not normally distributed data. In case of multiple comparisons, two-way ANOVA test was employed. Whenever possible, data are reported by individual (not only by group) as to most transparently report inter-individual variation.

## RESULTS

### A novel mouse with construct validity for DDX3X syndrome

We sought to generate a novel mouse line with construct validity for loss-of-function mutations clinically associated with DDX3X syndrome. Two *Ddx3x* knockout (KO) lines have been published (17, 18, 31), but the female heterozygous progeny have not been examined. Also, one of these lines (18, 31) is a brain-specific conditional and thus has limited construct validity. Therefore, we first generated a floxed *Ddx3x* mutant mouse (*Ddx3x*^flox^) by introducing loxP sites flanking exon 2. We selected this exon because it has 100% identity with the human sequence and there are at least 13 individuals with pathogenic loss-of-function mutations affecting this exon (**Fig. 1A**), making the design salient to the genetics of the human disorder.

**Figure 1.**
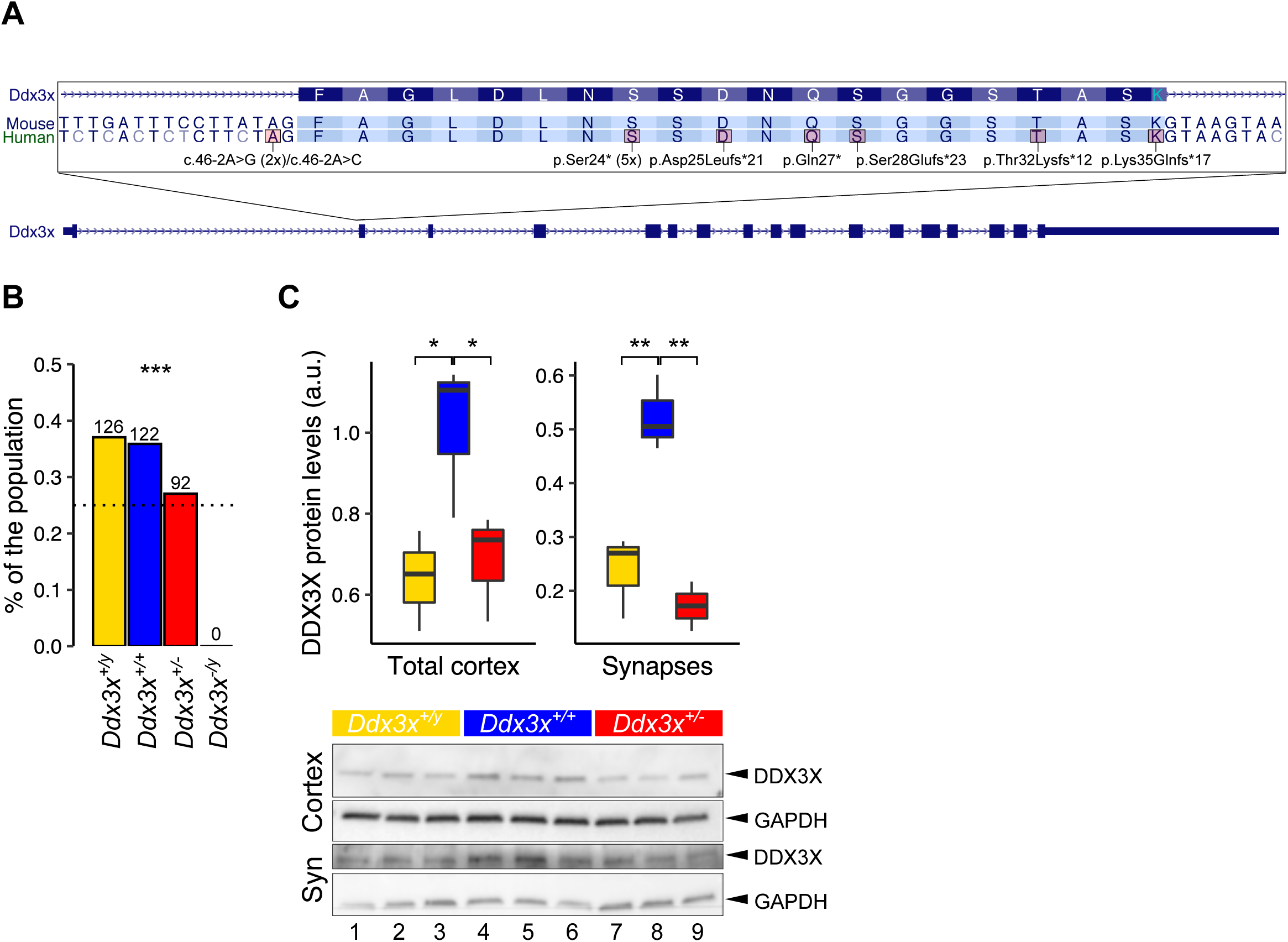
*Ddx3x*^+/−^ mice have construct validity for *DDX3X* loss-of-function mutations. **A) Disease relevance of the design.** The figure shows the Genome Browser view of *Ddx3x* exon 2, which was targeted to generate the *Ddx3x*^+/−^ line (see also **Fig. S1**). The sequences of murine and human exon 2 are identical. There are at least 13 pathogenic mutations clinically associated with DDX3X syndrome affecting exon 2: two mutations affecting the acceptor splice site of exon 2 [NM_001356.4:c.46-2A>G in two independent individuals (1, 3), c.46-2A>C (1)], the recurrent nonsense mutation p.Ser24* in 5 independent individuals [c.71C>A (2, 76), c71.C>G (76) or c71C>TA (3)], the nonsense p.Gln27* [c.79C>T (3)], the frameshift Asp25Leufs*21 (c.67_71dupCTCTTC, ClinVar #916064), the frameshift p.Ser28Glufs*23 [c.80dupA (3)], the frameshift p.Thr32Lysfs*12 (c.95delC, ClinVar # 872822), and the frameshift p.Lys35Glnfs*17 (c.95_98CAGC[3], ClinVar #803980). There is an additional mutation in the donor site of exon 1 [c.45+1G>T (2)] that likely affects splicing (not shown). **B) *Ddx3x*^+/−^ pups are born at lower rate, and *Ddx3x*^−/y^ die *in utero***. Percentage of the population across the four expected genotypes (*Ddx3x*^+/y^ in yellow; *Ddx3x*^+/+^ in blue; *Ddx3x*^+/−^ in red; and, *Ddx3x*^−/y^ in black) (Fisher’s exact test, ***p<0.001). **C) *Ddx3x*^+/−^ mice have reduced DDX3X expression at P21.** The plots show DDX3X protein expression, normalized to GAPDH, in total cortices (left panel) or purified synaptosomes (right panel) (n=3/genotype; mean ± SEM; Student’s t test; *p<0.05, **p<0.01). The figure shows the corresponding immunoblot for DDX3X and GAPDH in total cortices (upper panels) and purified synaptosomes (lower panels) of *Ddx3x*^+/y^ (yellow, lanes 1-3), *Ddx3x*^+/+^ (blue, lanes 4-6) and *Ddx3x*^+/−^ (red, lanes 7-9) mice (n=3/genotype).

As *Ddx3x* is indispensable for the formation of the placenta in mouse (17), we needed to avoid perturbing *Ddx3x* in extraembryonic tissues when generating the KO allele. We adopted the strategy used by Chen et al (2015) (17) and used a Sox2-Cre driver line to induce recombination only in the epiblast (not in extraembryonic tissues) at early gastrulation (19). The resulting progeny develop with a functional placenta (17).

We crossed the *Ddx3x*^flox/+^ females with Sox2-Cre/+ males to excise exon 2 and thus generate a *Ddx3x* targeted allele (**Fig. S1**). This breeding scheme is expected to generate: *Ddx3x*^flox/+^;+/+ females (hereafter referred to as *Ddx3x*^+/+^), *Ddx3x*^flox/y^;+/+ males (*Ddx3x*^+/y^), *Ddx3x*^flox/+^;Sox2-Cre/+ females (*Ddx3x*^+/−^), and, *Ddx3x*^flox/y^;Sox2-Cre/+ males (*Ddx3x*^−/y^). In line with earlier findings (17) and the dearth of male patients with pathogenic mutations in *DDX3X*, all *Ddx3x*^−/y^ male embryos died *in utero* (**Fig. 1B**). *Ddx3x*^+/−^ female pups were born at lower rate than expected (**Fig. 1B**), but did not show reduced viability from birth to the postnatal day 21 and survived to adulthood. To verify that *Ddx3x*^+/−^ mice faithfully recapitulate haploinsufficiency, we measured *Ddx3x* protein levels in cortices and purified cortical synapses. Female mice had higher *Ddx3x* protein expression in their cortices than males (**Fig. 1C**). This is in line with data showing that *DDX3X* is an escape gene across human tissues including the cortex (11, 12). *Ddx3x*^+/−^ mice have ~50% reduction in *Ddx3x* protein levels compared to *Ddx3x*^+/+^ littermates (**Fig. 1C**). Interestingly, both sex- and genotype-dependent differences in *Ddx3x* expression were observed in cortical synapses (**Fig. 1C**).

As individuals with DDX3X syndrome can have congenital anomalies, we subjected postnatal day 3 (P3) *Ddx3x*^+/+^ and *Ddx3x*^+/−^ mice to a full histopathological examination (n=5/genotype). We found no significant anatomical differences. We did observe a higher rate of acute suppurative alveolitis consistent with aspiration pneumonitis in *Ddx3x*^+/−^ pups (4/5 *Ddx3x*^+/−^ vs. 1/5 *Ddx3x*^+/+^), but that failed to reach statistical significance (Fisher exact test, *p*=0.21, data not shown).

### *Ddx3x*^+/−^ mice have widespread developmental delays

To capture the impact of *Ddx3x* haploinsufficiency on physical, sensory, and motor development, we monitored *Ddx3x*^+/−^ mice from birth to weaning using a standardized observation battery (32) adapted from the Fox scale (21, 22). All data were collected blind to genotype and across 4 independent cohorts (**Table S1**), each showing comparable results. *Ddx3x*^+/−^ pups had a growth delay (**Fig. 2A**) manifesting as lower body weight in adulthood (**Fig. 2B**). We conjectured that *Ddx3x*^+/−^ pups might be outcompeted by their siblings for access to the maternal milk supply. We therefore expected to see an inverse correlation between litter size and growth for the *Ddx3x*^+/−^ genotype. On the contrary, growth curves stratified based on the litter size, but irrespective to genotype: pups in smaller litters of all genotypes were at a disadvantage (**Fig. S2**). The higher rate of aspiration pneumonitis in *Ddx3x*^+/−^ pups, combined with the failure to thrive, suggests that *Ddx3x*^+/−^ pups had reduced competence in feeding. Notably, these observations align with the clinical phenotype, as females with DDX3X syndrome can have low body weight and feeding problems (1, 3, 4). In addition to failure to thrive, *Ddx3x*^+/−^ pups had a delay in eye opening (**Fig. 2C**) and pinna detachment (**Fig. S3A**). There were no delays noted in the development of the fur/skin or tooth eruption (**Fig. S3B-D**). These physical delays are again reminiscent of the human phenotype, as ophthalmological findings are common in DDX3X patients.

**Figure 2.**
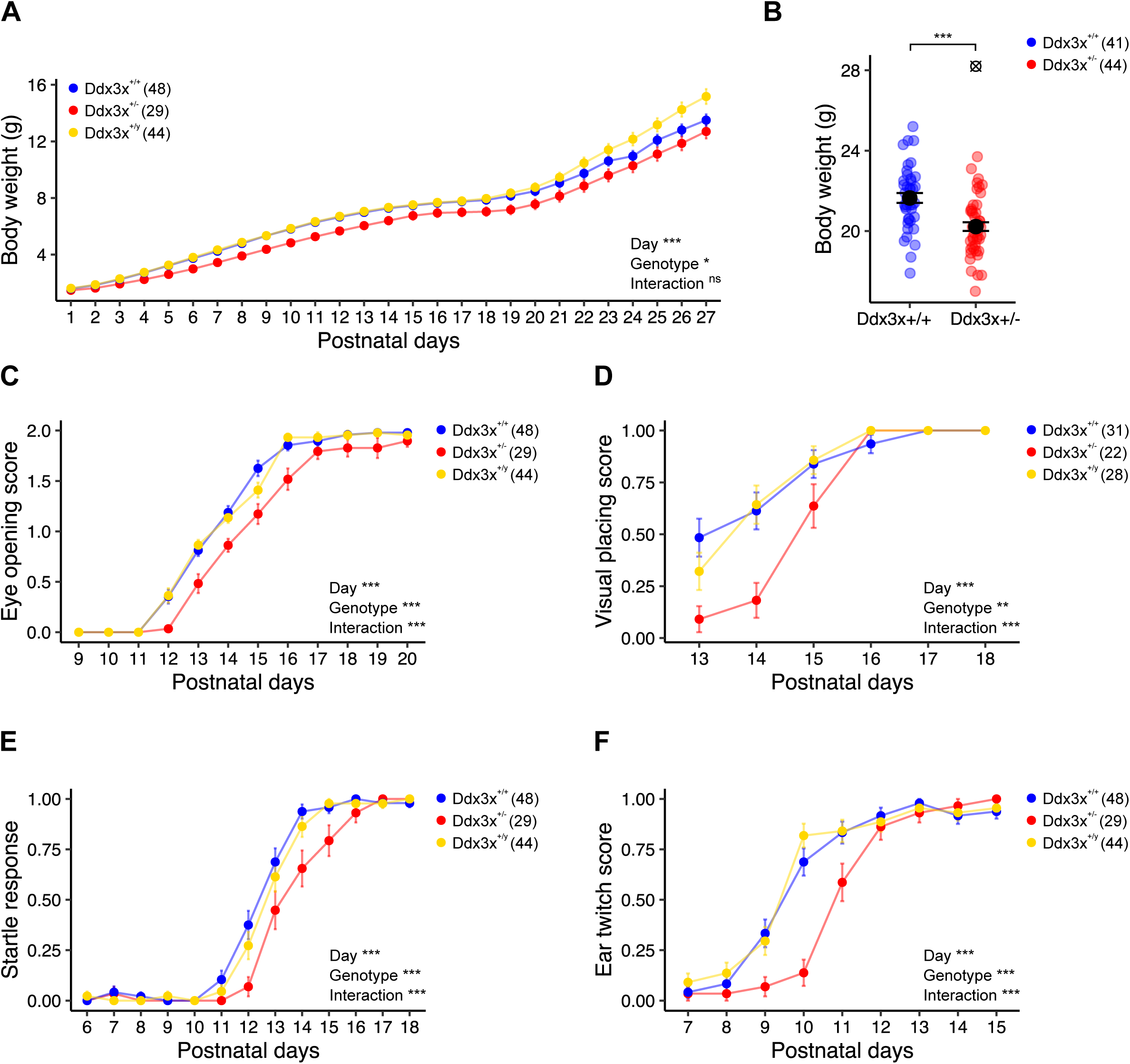
*Ddx3x*^+/−^ mice have postnatal physical and sensory delays. **A) *Ddx3x*^+/−^ pups show delayed growth.** The plot shows the body weight from postnatal day (P) 1 to 27 across the three genotypes (*n* shown in legend; mean ± SEM; repeated measure ANOVA between *Ddx3x*^+/+^ and *Ddx3x*^+/−^ genotypes). **B) *Ddx3x*^+/−^ adults have low body weight.** The plot shows the body weight of 10-week old mice (*n* shown in legend; mean ± SEM; Student’s t test after outliers removal and Shapiro-Wilk test for normality; ⊗ indicate outliers). **C) *Ddx3x*^+/−^ pups have a delay in eye opening.** The plot shows the eye opening scores (0, closed; 1, half-opened; 2, open) across the three genotypes (*n* shown in legend; mean ± SEM; repeated measure ANOVA between *Ddx3x*^+/+^ and *Ddx3x*^+/−^ genotypes). **D) *Ddx3x*^+/−^ pups show a delay in developing visual placing skills.** The plot shows the visual placing scores (0, response absent; 1, response present) across the three genotypes. Visual response was considered present (score 1) when pups lowered onto a flat surface reached out with the forepaws before the vibrissae touched the surface (*n* shown in legend; mean ± SEM; repeated measure ANOVA between *Ddx3x*^+/+^ and *Ddx3x*^+/−^ genotypes). **E) *Ddx3x*^+/−^ pups have a delay in developing a startle response to an auditory cue.** The plot shows the startle response score (0, response absent; 1, response present) across the three genotypes. Startle response was considered present (score 1) when pups turned their heads in response to an 80db click sound (*n* shown in legend; mean ± SEM; repeated measure ANOVA between *Ddx3x*^+/+^ and *Ddx3x*^+/−^ genotypes). **F) *Ddx3x*^+/−^ pups show a delay in developing an ear twitch response to a tactile stimulus.** The plot shows the reflex to ear twitch response (0, response absent; 1, response present) across the three genotypes (*n* shown in legend; mean ± SEM; repeated measure ANOVA between *Ddx3x*^+/+^ and *Ddx3x*^+/−^ genotypes). All data collected and scored blind to genotype. In all panels, *Ddx3x*^+/+^ (blue), *Ddx3x*^+/−^ (red), and *Ddx3x*^+/y^ (yellow). *p<0.05, **p<0.01, ***p<0.001; ns, non significant.

Physical delays were accompanied by somatosensory delays in processing visual, auditory, tactile, and vestibular cues. Delayed eye opening (**Fig. 2C**) translated into a delay in establishing visual placing competence (**Fig. 2D**). Additionally, *Ddx3x*^+/−^ pups were delayed in developing a startle reflex to an auditory stimulus (**Fig. 2E**) and a labyrinth-related reflex measured as aversion to a cliff ledge (**Fig. S3E**). Both alterations could be related to the delay in ear development (**Fig. S3A**). Responses to tactile cues varied: *Ddx3x*^+/−^ pups had delays in ear twitch reflex (**Fig. 2F**) and vibrissae-evoked forelimb placing (**Fig. S3F**), but not in forelimb grasping strength (**Fig. S3G**).

*Ddx3x*^+/−^ pups reached motor milestones later than their littermates. *Ddx3x*^+/−^ pups took longer to flip onto their paws from a supine position (righting reflex), suggesting hypotonia (**Fig. 3A**). Their motor coordination in response to vestibular cues of gravity was also affected, as measured by a delay in acquiring negative geotaxis skills (**Fig. 3B-D**). Neuromuscular development was abnormal, as *Ddx3x*^+/−^ pups showed alterations in both forelimbs and hindlimbs grip (**Fig. 3E-F, Fig. S4A**). No gross defects in locomotor activity in an open field (**Fig. S4B**) or air righting reflex (**Fig. S4C**) were noted. These motor delays closely resemble those observed in the DDX3X patient population.

**Figure 3.**
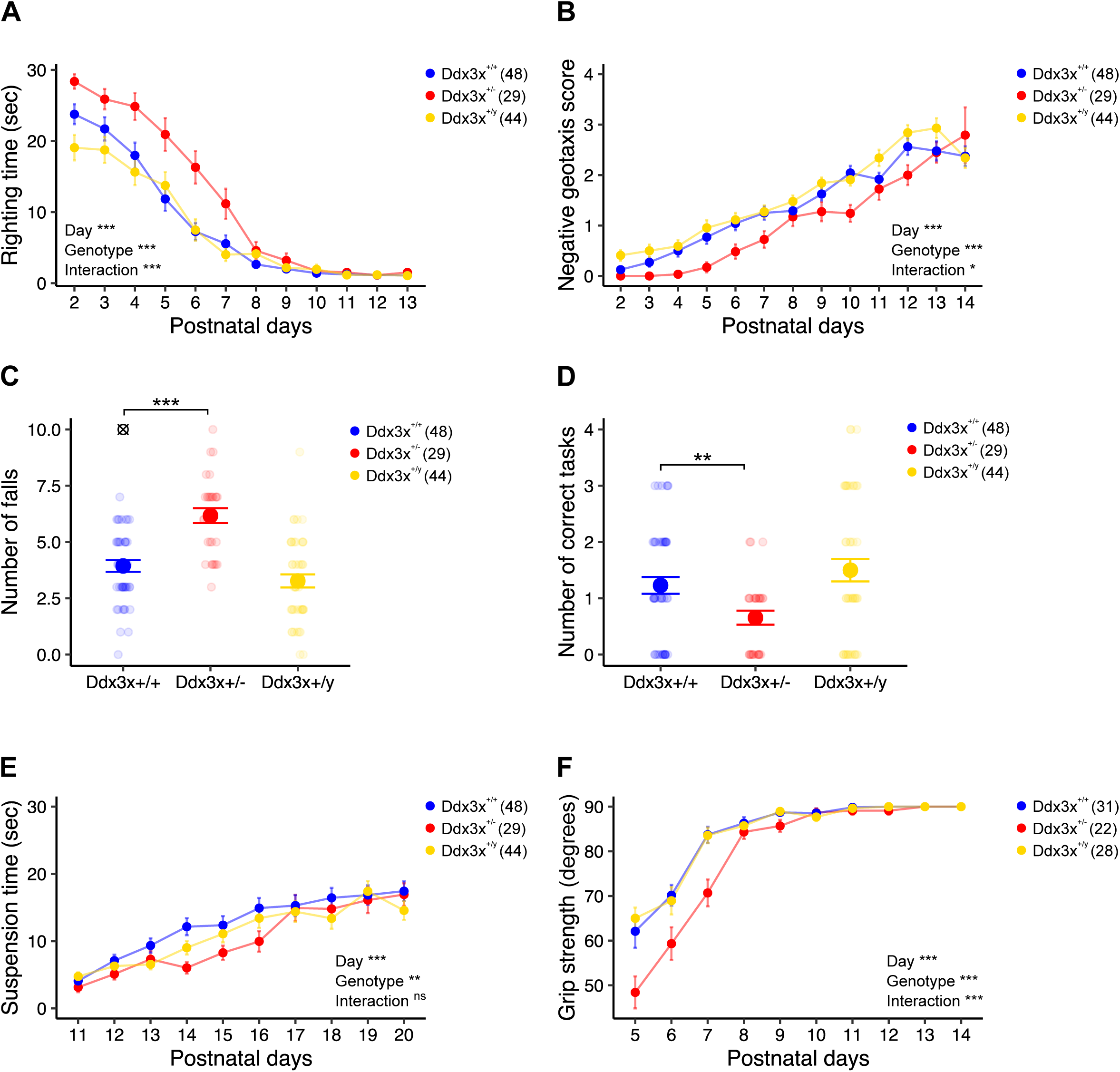
*Ddx3x*^+/−^ mice have postnatal motor delays. **A) *Ddx3x*^+/−^ pups have a delay in surface righting time.** The plot shows the time that it takes for a pup to turn on the four paws from a supine position across the three genotypes (*n* shown in legend; mean ± SEM; repeated measure ANOVA between *Ddx3x*^+/+^ and *Ddx3x*^+/−^ genotypes). **B) *Ddx3x*^+/−^ pups have a delay in acquiring negative geotaxis skills.** The plot shows the response of the pup when placed head down on a mesh covered platform at a 45° angle (0, fall; 1, stay/move down/walk down; 2, turn and stay; 3, turn, move up, and stay; 4, turn and move up to the top) across the three genotypes (*n* shown in legend; mean ± SEM; repeated measure ANOVA between *Ddx3x*^+/+^ and *Ddx3x*^+/−^ genotypes). **C) *Ddx3x*^+/−^ pups tend to fall more in a negative geotaxis task.** The plot shows the number of falls (score 0) per pup across the three genotypes (*n* shown in legend; mean ± SEM; Student’s t test after outliers removal and Shapiro-Wilk test for normality; ⊗ indicate outliers). **D) *Ddx3x*^+/−^ pups have a lower rate of success in a negative geotaxis task.** The plot shows the number of completed tasks (score 4) per pup across the three genotypes (*n* shown in legend; mean ± SEM; Student’s t test after outliers removal and Shapiro-Wilk test for normality; ⊗ indicate outliers). **E) *Ddx3x*^+/−^ pups have reduced motor endurance and performance in a rod suspension test.** The plot shows the time that it takes for a pup to fall off a rod across the three genotypes (*n* shown in legend; mean ± SEM; repeated measure ANOVA between *Ddx3x*^+/+^ and *Ddx3x*^+/−^ genotypes). **F) *Ddx3x*^+/−^ pups have reduced grip strength.** Grip strength was tested by placing pups on a mesh grid and rotating the grid from a horizontal to vertical position. The plot shows the angle of rotation to which pups fall off the grid across the three genotypes (*n* shown in legend; mean ± SEM; repeated measure ANOVA between *Ddx3x*^+/+^ and *Ddx3x*^+/−^ genotypes). All data collected and scored blind to genotype. In all panels, *Ddx3x*^+/+^ (blue), *Ddx3x*^+/−^ (red), and *Ddx3x*^+/y^ (yellow). *p<0.05, **p<0.01, ***p<0.001; ns, non significant.

Overall, we found that *Ddx3x*^+/−^ pups follow abnormal trajectories that impact physical, sensory and motor development.

### *Ddx3x*^+/−^ mice have hyperactivity- and anxiety-related behaviors, as well as associative memory deficits

With time, *Ddx3x*^+/−^ pups caught up with most developmental landmarks. However, the sensorimotor delays might disrupt exposure to stimuli during sensitive periods for brain developmental and plasticity, and thus translate into behavioral abnormalities later in life. We therefore sought to examine emotion states, cognition, and sociability of *Ddx3x*^+/−^ adult mice using a behavioral testing battery. All data were collected blind to genotype, and across four independent cohorts not employed for developmental testing (**Table S2**). During handling one week prior to behavioral testing, we noted no differences in physical appearance other than lower body weight (**Fig. 2B**). *Ddx3x*^+/−^ mice, however, showed changes in spontaneous general activity (**Fig. S5A**), reduced limb withdrawal indicative of reduced nociception (**Fig. S5B**), and reduced provoked biting reflex (**Fig. S5C**). We also noted a higher defecation index in *Ddx3x*^+/−^ mice compared to littermates (**Fig. S5D**), which is relevant given that gastrointestinal problems are frequent in DDX3X syndrome (1, 3, 4).

When assessed for free locomotor activity in an open field test, *Ddx3x*^+/−^ mice explored the arena more and moved faster than their littermates (**Fig. 4A-B, Fig. S6A**), in spite of the prominent early postnatal motor delays (**Fig. 3**). These observations indicate a hyperactivity-related phenotype. While *Ddx3x*^+/−^ mice entered the center zone as often as their littermates (**Fig. S6B**), it took them longer to enter it and they spent less time in it (**Fig. 4B-C**). This increase in latency and degree of thigmotaxis indicate an anxiety-like behavior. *Ddx3x*^+/−^ mice did not show, however, gross deficits when assessed in an elevated plus maze test (**Fig. S6C**).

**Figure 4.**
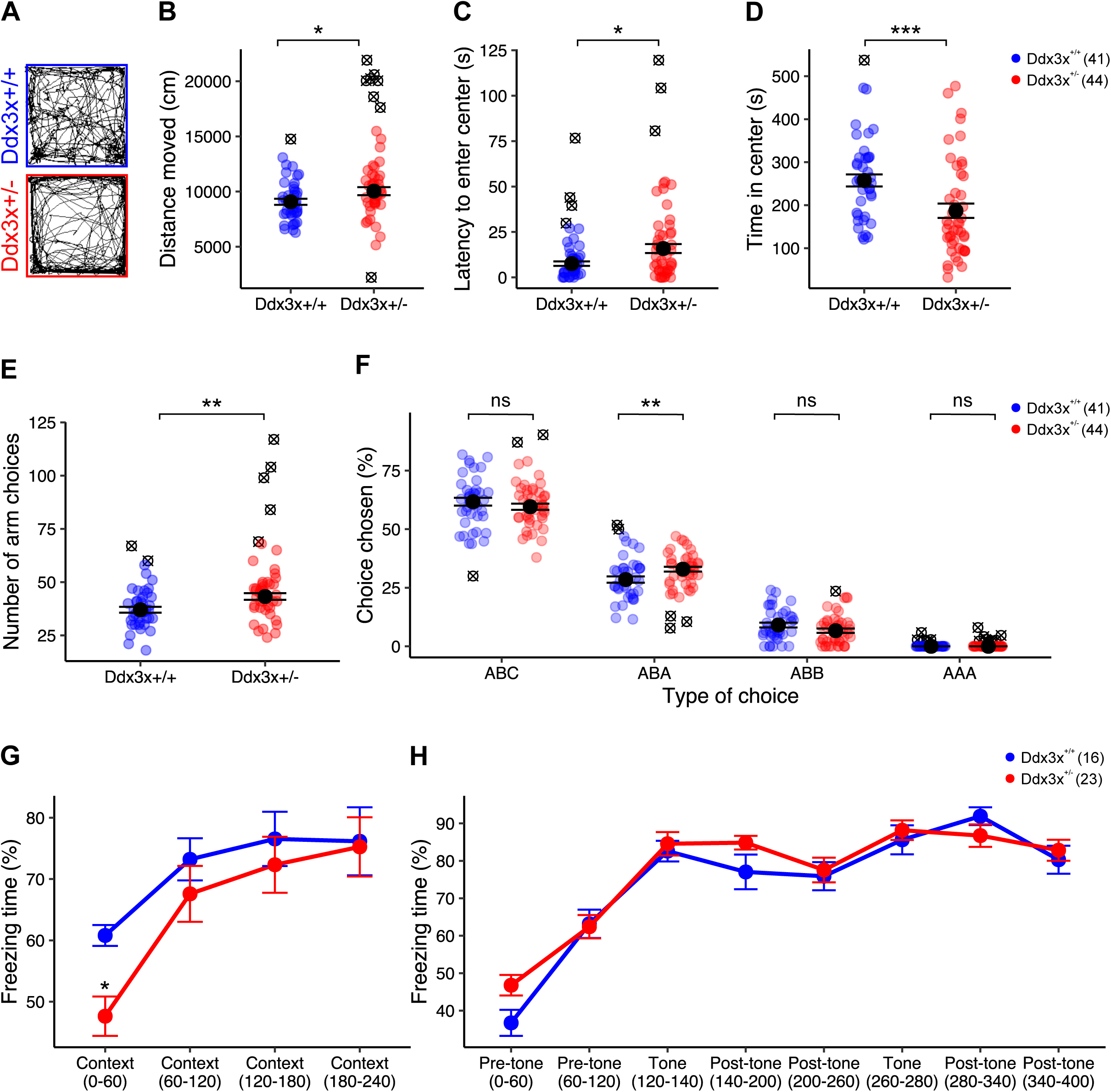
*Ddx3x*^+/−^ mice have hyperactivity- and anxiety-like behaviors. **A-B) *Ddx3x*^+/−^ mice show increased locomotor activity in open field test.** Panel A shows an example of activity of individual mice in a 10-min interval. The plot in B shows the distance covered in the 30-min open field test (*n* shown in legend; mean ± SEM; Student’s t test after outliers removal and Shapiro-Wilk test for normality; ⊗ indicate outliers). **C) *Ddx3x*^+/−^ mice display increased latency to enter the center zone.** The plot shows the latency to enter the center zone for the two genotypes (*n* shown in legend; mean ± SEM; Wilcoxon signed-rank test after outliers removal and Shapiro-Wilk test for normality; ⊗ indicate outliers). **D) *Ddx3x*^+/−^ mice have increased thigmotaxis.** The plot shows the time spent in the center zone for the two genotypes (*n* shown in legend; mean ± SEM; Wilcoxon signed-rank test after outliers removal and Shapiro-Wilk test for normality; ⊗ indicate outliers). **E) *Ddx3x*^+/−^ mice have increased activity in Y maze spontaneous alternation test.** The plot shows the number of arms explored in a 15 min test for the two genotypes (*n* shown in legend; mean ± SEM; Student’s t test after outliers removal and Shapiro-Wilk test for normality; ⊗ indicate outliers). **F) Short-term working memory is overall intact in *Ddx3x*^+/−^ mice in Y maze spontaneous alternation test.** The plot shows the percentage of choices, broken down by choice type (*n* shown in legend; mean ± SEM; Student’s t test after outliers removal and Shapiro-Wilk test for normality; ⊗ indicate outliers). ‘ABC’ indicates correct alternations, while ‘ABA’, ‘ABB’, ‘AAA’ errors. **G) *Ddx3x*^+/−^ mice display weaker initial recall of contextual fear memory.** After 24 hours from contextual and cued training (see **Fig. S6D**), mice were exposed to the same environment of training, but without the administration of tones or shocks, for 240 sec. Locomotor activity was recorded. The plot shows the % of time the mice spent freezing during this 240 sec testing trial (*n* shown in legend; mean ± SEM; repeated measure ANOVA over the first 120 sec). **H) *Ddx3x*^+/−^ mice show no changes in cued fear memory.** After 24 hrs from contextual testing (**G**), mice were exposed to a new environment, with administration of two tones without subsequent shock exposure. Locomotor activity was recorded. The plot shows the % of time the animals spent freezing during this 400 sec testing trial. All data collected and scored blind to genotype. In all panels, *Ddx3x*^+/+^ (blue) and *Ddx3x*^+/−^ (red). *p<0.05, **p<0.01, ***p<0.001; ns, non significant.

We also measured short-term working memory using a Y maze spontaneous alternation test. *Ddx3x*^+/−^ mice explored the arms more than their littermates (**Fig. 4E**), corroborating the hyperactivity-related phenotype observed in open field. There was no difference in the number of spontaneous alternations (‘ABC’, **Fig. 4F**) between the two genotypes, but *Ddx3x*^+/−^ mice re-entered the same arm after exploring a new one at higher frequency than their littermates (error ‘ABA’, **Fig. 4F**). Overall, these data suggest no overt deficits in short-term working memory.

We also tested associative learning and memory using the contextual and cued fear conditioning paradigm. After habituation and prior to training, both genotypes showed low baseline levels of freezing response (**Fig. S6D**). During training, mice were exposed to three tone-shock pairs (**Fig. S6D**). 24 hours after training, mice were placed in the same context without presentation of tone or shock. While both genotypes showed a freezing behavior elicited by the context, which increased over time, initial recall was weaker in *Ddx3x*^+/−^ mice compared to their littermates (**Fig. 4G**). The deficit was specific to contextual memory, as *Ddx3x*^+/−^ mice showed the same degree of freezing behavior than their littermates evoked by the tone in a new context (**Fig. 4H**). Since *Ddx3x*^+/−^ mice were indistinguishable from their littermates during training or cued testing (**Fig. S6D, Fig. 4H**), the reduced freezing during contextual testing is not a manifestation of their hyperactivity but a genuine memory deficit. *Ddx3x*^+/−^ mice did not show differences in sociability measured in a three-chamber social approach test (**Fig. S7A**) or recognition memory in a novel object recognition task (**Fig. S7B**).

Together, these data show that *Ddx3x*^+/−^ mice display hyperactivity- and anxiety-like behaviors, accompanied by contextual fear memory deficits.

### *Ddx3x*^+/−^ mice have motor deficits

Movement disorders and gait abnormalities are prevalent in the DDX3X patient population (1, 3, 4) and *Ddx3x*^+/−^ pups display pronounced motor delays (**Fig. 3**). To examine motor function later in life, we first assessed gait in juvenile and adult *Ddx3x*^+/−^ and *Ddx3x*^+/+^ mice. We found a transient alteration in gait (**Fig. S8**). Specifically, *Ddx3x*^+/−^ mice first showed a reduction in sway distance and distance between the back paws at P19, followed by a reduction in stride and diagonals lengths at P22 (**Fig. S8**). By P30, these changes were no longer apparent.

While adult *Ddx3x*^+/−^ mice had normal gait, we reasoned that defects in motor function and learning might be manifesting during more challenging tasks. We therefore used the accelerating rotarod test to measure motor coordination, endurance, and learning over two days, 24 hrs apart, each including 3 trials (**Table S2**). Adult (4-month old) mice of both genotypes improved their performance (latency to fall) over the trials, showing no deficits in short (1 hr) or long-term (24 hrs) motor learning (**Fig. 5A**). However, motor performance of *Ddx3x*^+/−^ mice was suboptimal, as mutant mice had a shorter latency to fall across all trials compared to their littermates (**Fig. 5A**).

**Figure 5.**
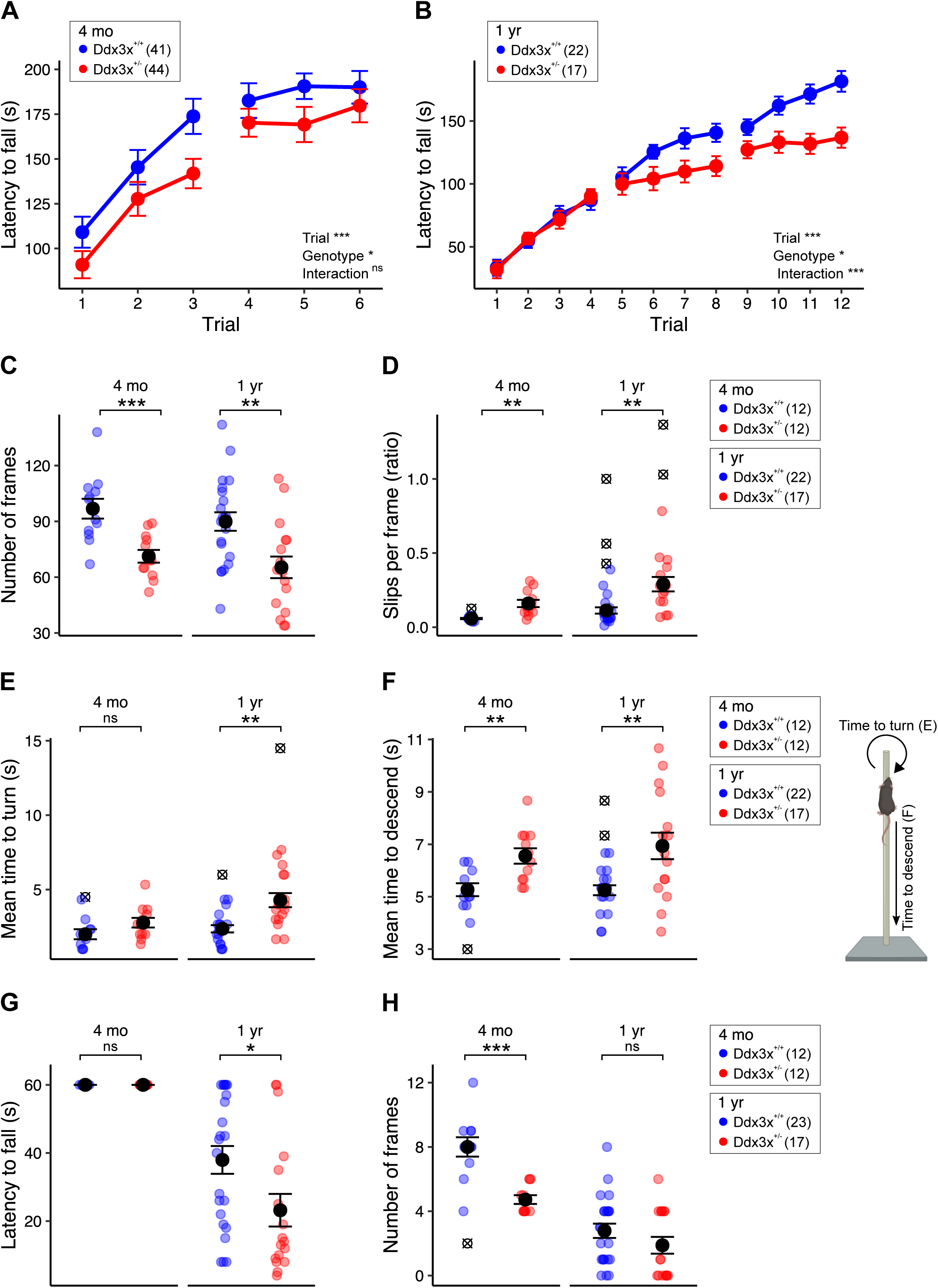
*Ddx3x*^+/−^ mice have other motor deficits and decline with age. **A) Adult *Ddx3x*^+/−^ mice have suboptimal motor performance on a rotarod test.** Trials were conducted one-hour apart with an acceleration over 5 minutes, and trials 1-3 were performed 24 hrs before trials 4-6, to assess both short-term (1 hour) and long-term (24 hrs) motor learning. The plot shows the latency to fall from the rod (*n* shown in legend; mean ± SEM; repeated measure ANOVA). **B) Ageing *Ddx3x*^+/−^ mice have altered motor learning on a rotarod test.** Trials were conducted one-hour apart with an acceleration over 5 min, with trials 1-4, 5-8 and 9-12 performed 24 hrs apart. The plot shows the latency to fall from the rod (*n* shown in legend; mean ± SEM; repeated measure ANOVA). **C-D) *Ddx3x*^+/−^ mice had impaired motor coordination in the balance beam test.** Panel C shows the number of frames covered (*n* shown in legend in panel D; mean ± SEM; Student’s t test after outliers removal and Shapiro-Wilk test for normality). Panel D shows the number of slips per frame (*n* shown in legend; mean ± SEM; Student’s t test (4 months old) and Wilcoxon signed-rank test (one-year old) after outliers removal and Shapiro-Wilk test for normality; ⊗ indicate outliers). **E) *Ddx3x*^+/−^ mice have age-dependent deficits in turning in the vertical pole test.** The plot shows the mean time to turn on the top of the vertical pole (*n* shown in legend in panel F; mean ± SEM; Student’s t test after outliers removal and Shapiro-Wilk test for normality; ⊗ indicate outliers). **F) *Ddx3x*^+/−^ mice have altered balance during the vertical pole test.** The plot shows the mean time to descend the vertical pole (*n* shown in legend; mean ± SEM; Student’s t test after outliers removal and Shapiro-Wilk test for normality; ⊗ indicate outliers). **G) *Ddx3x*^+/−^ mice have age-dependent changes in endurance in the wire hanging test.** The plot shows the latency to fall (*n* shown in legend in panel H; mean ± SEM; Wilcoxon signed-rank test (one-year old) after outliers removal and Shapiro-Wilk test for normality). **H) *Ddx3x*^+/−^ mice show changes in motor ability in the wire hanging test.** The plot shows the number of frames covered by the mice (*n* shown in legend; mean ± SEM; Wilcoxon signed-rank test after outliers removal and Shapiro-Wilk test for normality; ⊗ indicate outliers). All data collected and scored blind to genotype. In all panels, *Ddx3x*^+/+^ (blue) and *Ddx3x*^+/−^ (red). *p<0.05, **p<0.01, ***p<0.001; ns, non significant.

While the natural history of DDX3X syndrome has yet to be mapped, the oldest affected female reported so far (47 year old) suffered from a profound motor decline (4). We therefore probed motor function in 1 year-old mice (**Table S2**). To dissect in greater depth the effect on learning, we conducted testing over 3 days, 24 hrs apart, each including 4 trials, in 2 independent cohorts. In addition to poor motor performance, 1 year-old *Ddx3x*^+/−^ mice showed a decline in their motor function and learning (**Fig. 5B**).

To further assess fine motor coordination and balance, we used a balance beam test and a vertical pole test. When walking on the balance beam, adult (4-month old) *Ddx3x*^+/−^ mice covered a shorter distance on the beam (**Fig. 5C**) and slipped more frequently (**Fig. 5D**) than their littermates. Ageing (1-year old) *Ddx3x*^+/−^ mice had an even more pronounced loss of balance (**Fig. 5C-D**). When climbing on a vertical pole, 4-month old *Ddx3x*^+/−^ mice had no deficits in turning (**Fig. 5E**) but it took them longer to descend the pole (**Fig. 5F**) than their littermates. One-year old *Ddx3x*^+/−^ mice had difficulties both turning and descending (**Fig. 5E-F**).

To assess neuromuscular strength, we employed a wire hanging test. Although 4 month-old mice *Ddx3x*^+/−^ mice were able to hang on the wire as long as their littermates (**Fig. 5G**), they covered a reduced number of segments while moving on the wire (**Fig. 5H**), indicating intact motor endurance but altered coordination. Aging animals of both genotypes had, as expected, reduced strength, but *Ddx3x*^+/−^ mice performed significantly worse than their littermates (**Fig. 5G**).

These data demonstrate that *Ddx3x* haploinsufficiency results in significant motor deficits that are exacerbated with ageing.

### *Ddx3x*^+/−^ mice have reduced brain volume, with some regions disproportionally affected

Individuals diagnosed with DDX3X syndrome frequently have microcephaly and brain malformations, including agenesis of the corpus callosum, enlarged ventricles, and polymicrogyria (1, 3, 4). *Ddx3x* complete knockout leads to underdeveloped brain (17, 18), but the impact of haploinsufficiency, which is closer to the human condition, has not been studied. To bridge this gap, we performed magnetic resonance imaging (MRI) on *Ddx3x*^+/−^ and *Ddx3x*^+/+^ littermates at P3, a stage where developmental delays begin to emerge. We found that *Ddx3x*^+/−^ have a ~10% reduction in overall brain volume (n=10/genotype; Student’s t test, *p*=0.03) and this was also found in the majority of the regions assessed (**Fig. 6**, **Table S3**). As expected, given the low body weight in *Ddx3x*^+/−^ mice (**Fig. 2**), these brain volumes differences are no longer apparent when normalized to body weight (**Table S3**). Therefore, these changes in brain volume are part of an overall growth delay, and not a pure microcephaly phenotype.

**Figure 6.**
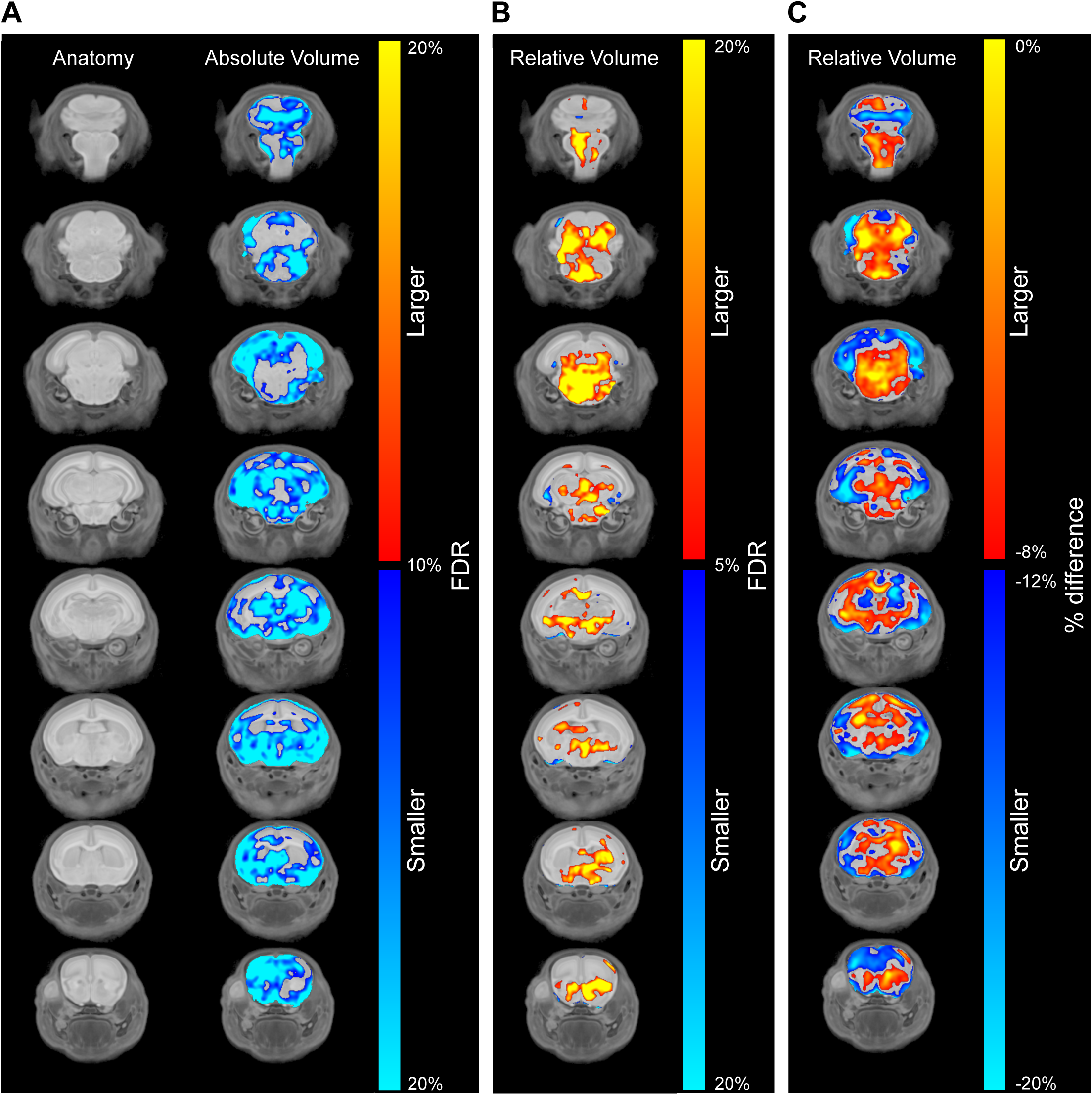
*Ddx3x*^+/−^ mice have voxelwise differences in brain volume. **A) *Ddx3x^+/−^* pups have reduced brain volume.** Coronal flythrough from anterior (top row) to posterior (bottom row), showing brain anatomy (left column) and absolute differences in *Ddx3x*^+/−^ vs *Ddx3x*^+/+^ mice (right column) at P3. The color coding indicates the false discovery rate (FDR) for regions that are smaller (blue gradient) or larger (red gradient) in *Ddx3x*^+/−^ compared to *Ddx3x*^+/+^ mice (n=10/genotype). **B) Changes in relative brain volume in *Ddx3x*^+/−^ pups.** Same as in A, showing voxelwise changes relative to the overall reduction in brain volume (n=10/genotype). **C) Specific brain regions are disproportionally reduced in *Ddx3x*^+/−^ pups.** Same as in A, with the color coding indicating the percentage of difference between *Ddx3x*^+/−^ and *Ddx3x*^+/+^ mice for regions that are disproportionally smaller (blue gradient) or larger (red gradient) that the ~10% reduction in overall brain volume (n=10/genotype). All data used to generate these plots, include individual-level data, are reported in **Table S3**. All data collected and analyzed blind to genotype.

**Figure 7.**
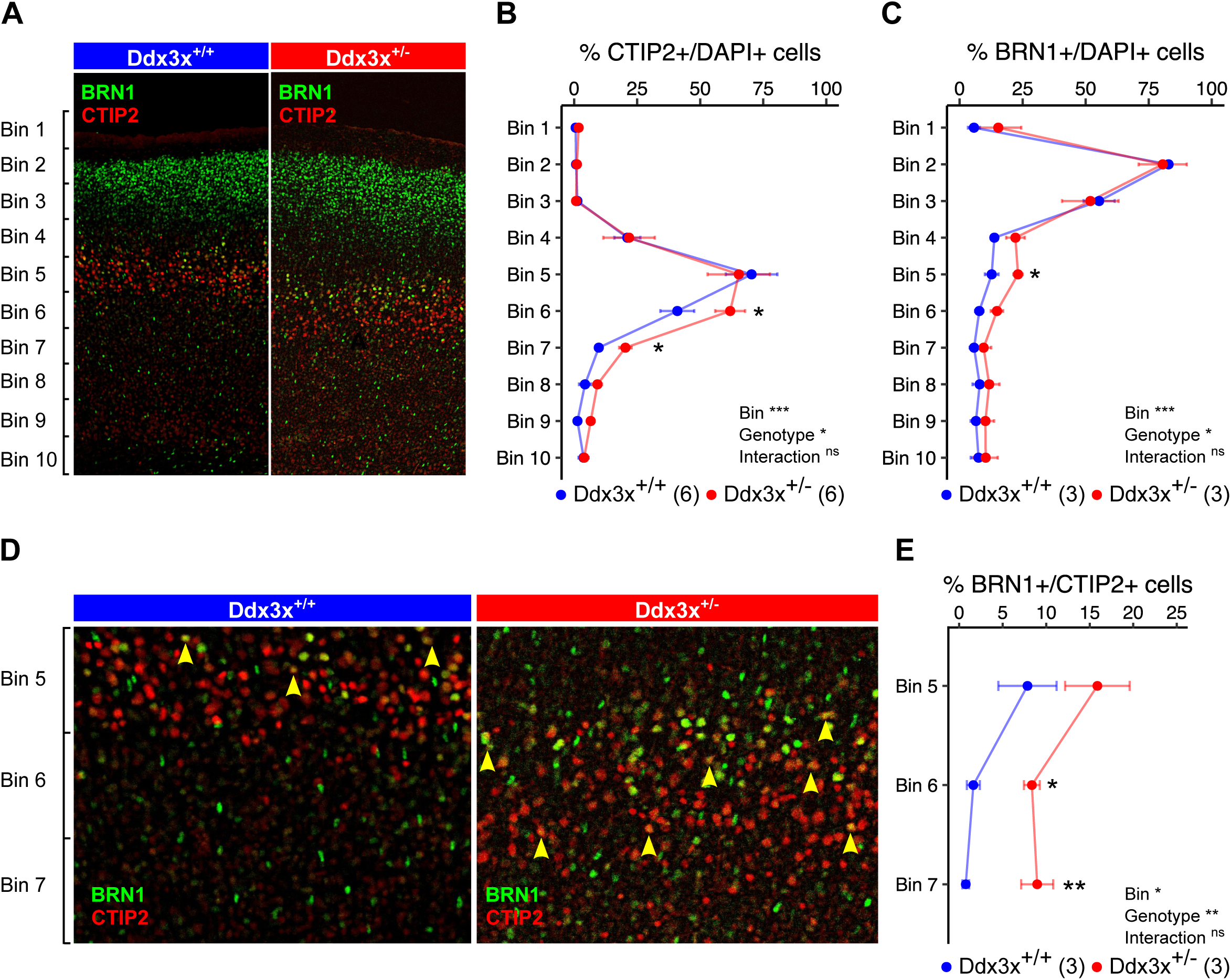
*Ddx3x*^+/−^ mice have altered cortical lamination. **A) *Ddx3x*^+/−^ mice show a misplacement of both subcerebral (ScPN) and intra-telencephalic (IT) projection neurons in the developing cortex.** Representative confocal images of coronal sections of primary motor cortex from *Ddx3x*^+/+^ and *Ddx3x*^+/−^ mice at P3, immunostained for CTIP2 (red), a marker of ScPN and BRN1 (green), a marker of IT in UL and DL. As expected, CTIP2+ ScPN are restricted to layer 5, while BRN1+ IT are predominantly in upper layers in control mice. **B) CTIP2+ ScPN extend deeper in *Ddx3x*^+/−^ mice.** Distribution of the percentage of cells (DAPI+) that are positive to the ScPN marker CTIP2, across ten equally sized bins from the pia (bin 1) to the ventricle (bin 10) in *Ddx3x*^+/+^ and *Ddx3x*^+/−^ mice (*n* shown in legend; 6-8 sections/mouse, with outliers across sections removed; mean ± SEM; Two-way ANOVA, followed by Student’s t test after outliers removal). **C) BRN1+ IT extend deeper in *Ddx3x*^+/−^ mice.** Distribution of the percentage of cells (DAPI+) that are positive to the IT marker BRN1, across ten equally sized bins from the pia (bin 1) to the ventricle (bin 10) in *Ddx3x*^+/+^ and *Ddx3x*^+/−^ mice (*n* shown in legend; 6-8 sections/mouse, with outliers across sections removed; mean ± SEM; Two-way ANOVA, followed by Student’s t test after outliers removal). **D) CTIP2+ and BRN1+ are mostly mutually exclusive, but not in *Ddx3x*^+/−^ mice.** The panels show a magnification of the bins 5-7 regions from panel A. As expected, in control mice, a minority of CTIP2+ neurons is positive to BRN1 (yellow arrows). **E) In *Ddx3x*^+/−^ mice, there are more CTIP2+BRN1+ neurons in deep layers.** Distribution of the percentage of CTIP2+ ScPN that are also positive to BRN1, across bins 5-7, where CTIP2+ ScPN are most abundant (*n* shown in legend; 6-8 sections/mouse, with outliers across sections removed; mean ± SEM; Two-way ANOVA, followed by Student’s t test after outliers removal). All data collected and scored blind to genotype. In all panels, *Ddx3x*^+/+^ (blue) and *Ddx3x*^+/−^ (red). *p<0.05, **p<0.01, ***p<0.001; ns, non significant.

To visually identify brain regions that had a larger deviation, we focused on those that were either larger (less than 8% reduction) or smaller (more than 12% reduction) than expected based on the ~10% overall reduction. We found that the volumetric reduction manifests disproportionally in the neocortex, part of the olfactory system (i.e., olfactory bulbs, the olfactory tubercle, and the lateral olfactory tract), and part of the limbic system (i.e., amygdala and the hippocampus, especially the subiculum) (**Fig. 6C**, **Table S3**). Interestingly, some of these areas are functionally interconnected and relevant for the behaviors disrupted in *Ddx3x*^+/−^ mice. For example, the hippocampal-amygdala circuit is indispensable for encoding of contextual fear memory (33, 34), which is altered in *Ddx3x*^+/−^ mice (**Fig. 4G**). Further, the amygdala, in concert with the cortex and the hippocampus, is critical to modulating anxiety-related behaviors (35), again abnormal in in *Ddx3x*^+/−^ mice (**Fig. 4C-D**).

Overall, these findings confirm that *Ddx3x* haploinsufficiency causes an overall growth delay that affects brain development as well. These changes are visible already during early postnatal life and functionally relate with the behavioral deficits later in adulthood.

### *Ddx3x*^+/−^ mice have abnormal neocortical lamination

Brain imaging findings indicate cortical thinning in *Ddx3x*^+/−^ mice. This is compatible with evidence showing that *Ddx3x* regulates cortical neurogenesis: radial glia cells knocked down for *Ddx3x* at E14.5 yield less neurons at E17.5 (3). It is unknown, however, whether this reduced neurogenesis is at the expense of a specific glutamatergic population and/or if it ultimately alters the laminar cytoarchitecture of the cortex. We therefore asked whether *Ddx3x* orchestrates the balance of intra-telencephalic and corticofugal glutamatergic neurons, which defines the lamination of the cortex. Intra-telencephalic neurons (IT) reside in both upper layers (UL) and deep layers (DL) of the cortex, and project to the ipsilateral cortex or the contralateral cortex via the corpus callosum (the latter, callosal projection neurons, CPN). Corticofugal neurons project to a variety of subcerebral targets (ScPN, including the spine and, the pons) or to the thalamus (CthPNs). These populations reside in DL; specifically, ScPN are mostly in layer V and CthPNs mostly in layer VI. Cell fates in the developing cortex are largely shaped by the expression of certain transcription factors, which can therefore be used to mark specific populations. For example, UL IT express BRN1/POU3F1 (36–38), while UL or DL CPN express SATB2 (39–43), with the caveat that SATB2 is detected also in a minority of ScPN (44, 45). ScPN, in contrast, can be discriminated by the expression of CTIP2/BCL11B (39–46).

Using BRN1, SATB2, and CTIP2 as population-specific markers, we immunostained brain sections from *Ddx3x*^+/−^ and *Ddx3x*^+/+^ mice at P3. We first examined the total number of cells (DAPI+), as well as CTIP2+, BRN1+ or SATB2+ populations, in coronal sections from the primary (M1) and secondary (M2) motor cortices. We observed a higher number of cells in M2 (but not in M1) of *Ddx3x*^+/−^ mice compared to their littermates (**Fig. S9**). This increase was largely attributable to an excess of CTIP2+ neurons, which showed a ~1.5 fold increase in *Ddx3x*^+/−^ cortices compared to controls (**Fig. S9**). There was also a modest excess of CTIP2+ neurons in M1, which did not, however, lead to a significant increase in the total number of cells (**Fig. S9**).

We then examined the relative distribution of CTIP2+, BRN1+, or SATB2+ populations across ten equally sized bins from the pia to the ventricle (**Fig. 7, Fig. S10**). The distributions of both CTIP2+ ScPN and BRN1+ IT neurons were altered in M1 of *Ddx3x*^+/−^ mice. Specifically, the majority of CTIP2+ ScPN neurons were residing, as expected, in layer V, but there was a subpopulation extending deeper in *Ddx3x*^+/−^ cortices compared to controls (**Fig. 7A, B**). Similarly, BRN1+ IT were mostly in UL, but significantly shifted in deeper layers in the M1 cortices of *Ddx3x*^+/−^ mice (**Fig. 7A, C**). These differences were not noted in M2 (**Fig. S10A-C**). SATB2+ CPN were detected in both UL and DL, and did not show differences between genotypes (**S10D-E**).

While CTIP2 and BRN1 are mostly mutually exclusive, there is a population of neurons expressing both BRN1 and CTIP2 that appears in the presumptive layer V at E15.5 and is still visible at P0 (36). We therefore asked whether the misplaced CTIP2+ neurons in *Ddx3x*^+/−^ mice are also BRN1+. We counted the double positive neurons in bins 5-7 of primary motor cortices of *Ddx3x*^+/−^ and *Ddx3x*^+/−^ littermates. Although the CTIP2+BRN1+ population represents a minority of the CTIP2+ neurons, it was detected in greater proportions in *Ddx3x*^+/−^ mice (**Fig. 7D-E**). This suggests that there might be defective cell fate specification of ScPN populations, also expressing IT markers.

These observations indicate that *Ddx3x* haploinsufficiency causes defects of intra-telencephalic and cortico-fugal populations in the motor cortex, which could result from alterations in neurogenesis or cell fate specification.

## DISCUSSION

To our knowledge, this is the first developmental, behavioral, and cellular characterization of a mouse model of DDX3X syndrome. *Ddx3x*^+/−^ mice have robust delays in physical, sensory, and motor milestones that closely resemble those seen in the patient population (1, 3, 4). Adult *Ddx3x*^+/−^ mice have hyperactivity, anxiety-like behaviors, memory deficits, and motor alterations, again aligning with the human phenotype (1, 3, 4). Motor function deteriorates with age, which is compatible with the late-onset neurological decline described in the only adult with DDX3X syndrome reported to date (4). The developmental and behavioral phenotypes are accompanied by reduced brain volume, with disproportionally thinner cortex, and smaller amygdala and hippocampus. Within the motor cortex, glutamatergic populations are misplaced, resulting in defective cytoarchitecture. These structural findings are in line with the brain malformations frequently detected in individuals with DDX3X syndrome (1, 3, 4).

Our results emphasize the importance of both sensory and motor function in the context of complex neurodevelopmental phenotypes. Altered somatosensation is emerging as a critical symptom in individuals with NDDs (47), and is captured in the phenotype of several mouse models (20, 48–53). Here we find that *Ddx3x*^+/−^ pups are delayed in responding to visual, auditory, and tactile cues. Further, *Ddx3x*^+/−^ pups have an underdeveloped olfactory system, with smaller olfactory bulbs, olfactory tubercle and lateral olfactory tract. Abnormal sensory experience during early postnatal life is known to robustly affect cortical plasticity and adult behavior, as shown by paradigms of sensory deprivation in mice (54–57). Hence, some of the adult manifestations in *Ddx3x*^+/−^ mice might be rooted in limited sensory stimuli exposure during critical periods of plasticity.

Motor dysfunction is not only a prominent trait in DDX3X syndrome (1, 3, 4), but is often one of earliest signs raising concerns about neurodevelopment. Motor development follows a complex trajectory in *Ddx3x*^+/−^ mice. Early anomalies, including postnatal hypotonia and abnormal gait in juvenile age, appear to rectify over time but evolve into changes in coordination in adulthood and further loss of balance and endurance with ageing. Cortical circuits controlling these functions include the corticopontine and corticospinal tracts that are formed by the ScPN. As ScPN are altered in number and position in the motor cortex of *Ddx3x*^+/−^ mice, defects in corticopontine and/or corticospinal circuits are likely to contribute to the motor phenotype. Interestingly, clinically small pons and small inferior vermis can be observed in individuals with DDX3X syndrome (3), suggesting that the cortico-ponto-cerebellar tract might be altered in this condition. We did not observe, however, a pronounced reduction in the volume of the pons or cerebellum, Also, divergence in motor circuits between species might explain the transient nature of certain motor deficits (e.g., gait) in mice but not in humans. For example, the rubrospinal tract, which can compensate for corticospinal lesions even in adulthood (58), is underdeveloped in humans compared to mice.

In addition to the sensory and motor phenotypes, *Ddx3x*^+/−^ mice show anxiety-like behaviors, hyperactivity, and specific deficits in the initial context-dependent recall of fear memory. Albeit complex and interlinked, the circuits subserving these behaviors require the cortex, the amygdala, and the hippocampus (34, 35, 59), all found disproportionally diminished in volume in *Ddx3x*^+/−^ mice. Additionally, mouse models of NDDs often show similar behavioral anomalies contingent to altered balance in cortical glutamatergic populations. For example, loss of CthPN specification caused by conditional ablation of the NDD gene *Tbr1* (60, 61) in layer 6 results in anxiety-related phenotypes (62). 16p11.2 deletion mice show more UL neurons, accompanied by hyperactivity and hippocampal-dependent memory deficits, and these anomalies can be reversed by *in utero* pharmacotherapy (63).

DDX3X is an RNA helicase critical for the post-transcriptional regulation of gene expression, especially mRNA translation (3, 8–10). Post-transcriptional control is surfacing as a significant source of risk for neurodevelopmental disorders. For example, risk genes for NDDs are enriched in targets of post-transcriptional regulators that are in turn encoded by risk genes, including RNA-binding proteins FMR1 (64), CELF4 (65) and RBFOX1 (64, 66, 67). *Ddx3x* regulates the translation of a subset of mRNAs in both neural progenitors (3) and post-mitotic neurons (3, 10) and amongst them is *Rac1*, whose human orthologous gene is associated with ID (68).

In parallel, post-transcriptional control is emerging as key for shaping cortical development (65, 69–73), and our findings and those by Lennox et al (2020) (3) add to this growing body of evidence. Lennox et al (2020) found that *in utero* knockdown of ~25% of *Ddx3x* in E14.5 cortices results in an increase in the proliferating pool and a decrease in post-mitotic neurons. In this experimental setting, *Ddx3x* is knocked down transiently (E14.5-E17.5) and in a cell-autonomous fashion (sparsely in radial glia cells and their progeny). In our model, where one *Ddx3x* allele is ablated in all the cells of the developing cortex since their birth, we see an increased number of ScPN neurons and a misplacement of both UL IT and ScPN neurons. The two sets of results are not incompatible, as the phenotype we observe is cell population specific and can be the result of defects in neurogenesis, migration and/or cell fate specification. Many NDD risk genes, in fact, have been found to regulate multiple steps of cortical development (e.g., *Chd8* important for both neurogenesis (74) and migration (75)).

The mouse model presented here is designed with validity for *DDX3X* loss-of-function mutations leading to haploinsufficiency. However, clinical and functional data (3) suggest that there is at least a second disease mechanism at play, with certain missense mutations leading to a gain of function effect. A subset of missense/in-frame deletion mutations are clinically associated with polymicrogyria and result in more severe clinical outcomes (3). Also, these mutations induce the formation of ectopic RNA granules (3). Our mouse model is less informative with respect to this group of mutations, and likely recapitulates less faithfully their phenotype. One hint to this limitation relates to the hypotonia phenotype. While individuals with missense/in-frame deletions and polymicrogyria often exhibit a mix of hypertonia and hypotonia (3), those with loss-of-function mutations are disproportionally affected by hypotonia, as do the *Ddx3x*^+/−^ mice. However, most missense mutations do not yield polymicrogyria and likely act as hypomorphic alleles. We would expect these mutations to yield a phenotype similar, but attenuated, to what we see in the *Ddx3x*^+/−^ mice. Future studies modeling missense mutations in mice will be key to discriminate the various mechanisms driving DDX3X syndrome.

The natural history of DDX3X syndrome is unknown. Our data, alongside a single clinical report of a 47 year-old affected individual, indicate that there is motor decline with ageing. The parsimonious explanation for these observations is that DDX3X pathology is engrained in prenatal development but also maintained in adulthood. The implications of these findings are twofold. First, they motivate longitudinally studies, especially focusing on motor function, in the patient population. Second, they might inform on the temporal windows where pathology (and thus phenotype) can be modified. The convergence of these two lines of work, alongside a better understanding of the disease mechanisms at play, are important next steps for designing therapies for DDX3X syndrome.

## ACKNOWLEDGMENTS AND DISCLOSURES

Research reported in this publication was supported by the Eunice Kennedy Shriver National Institute Of Child Health & Human Development of the National Institutes of Health under Award Number R21HD097561. The content is solely the responsibility of the authors and does not necessarily represent the official views of the National Institutes of Health. The work was also supported by the Beatrice and Samuel A. Seaver Foundation and the Friedman Brain Institute Research Award by the Fascitelli Family. DCU was supported by Fondation pour la Recherche Médicale, the Philippe Foundation, and the Beatrice and Samuel A. Seaver Foundation. MGF was supported by the Beatrice and Samuel A. Seaver Foundation. MF was supported by a Dean’s Undergraduate Research Fund Grant for Spring 2020. We thank Brett Collins for lab managing support and Zeynep Akpinar for technical support. LRQ, JE, and JPL would like to acknowledge support from the Ontario Brain Institute and the Canadian Institute for Health Research.

Conceptualization: SDR, AB, DCU, MGF, ED; Investigation: SDR, AB, DCU, MGF, KN, DM, MF, SM, JE, LRQ; Formal analyses: SDR, AB, DCU, MGF, JE; Writing-original draft: SDR, AB, DCU, MGF; Writing – review and editing: SDR, AB, DCU, MGF, KN, DM, MF, SM, DEG, MRR, JDB, ED, JE, LRQ, JPL; Funding acquisition: SDR, JDB; Resources: SDR, MRR, JDB; DEG provided expertise on relationship to clinical phenotype.

The authors report no biomedical financial interests or potential conflicts of interest.

## Notes

### Competing Interest Statement

The authors have declared no competing interest.

## REFERENCES

1. Snijders Blok L, Madsen E, Juusola J, Gilissen C, Baralle D, Reijnders MR, et al. (2015): Mutations in DDX3X Are a Common Cause of Unexplained Intellectual Disability with Gender-Specific Effects on Wnt Signaling. American journal of human genetics. 97:343–352.

2. DDD Study (2017): Prevalence and architecture of de novo mutations in developmental disorders. Nature. 542:433–438.

3. Lennox AL, Hoye ML, Jiang R, Johnson-Kerner BL, Suit LA, Venkataramanan S, et al. (2020): Pathogenic DDX3X Mutations Impair RNA Metabolism and Neurogenesis during Fetal Cortical Development. Neuron. 106:404–420 e408.

4. Wang X, Posey JE, Rosenfeld JA, Bacino CA, Scaglia F, Immken L, et al. (2018): Phenotypic expansion in DDX3X - a common cause of intellectual disability in females. Ann Clin Transl Neurol. 5:1277–1285.

5. Lessel D, Schob C, Kury S, Reijnders MRF, Harel T, Eldomery MK, et al. (2017): De Novo Missense Mutations in DHX30 Impair Global Translation and Cause a Neurodevelopmental Disorder. American journal of human genetics. 101:716–724.

6. Balak C, Benard M, Schaefer E, Iqbal S, Ramsey K, Ernoult-Lange M, et al. (2019): Rare De Novo Missense Variants in RNA Helicase DDX6 Cause Intellectual Disability and Dysmorphic Features and Lead to P-Body Defects and RNA Dysregulation. American journal of human genetics. 105:509–525.

7. Paine I, Posey JE, Grochowski CM, Jhangiani SN, Rosenheck S, Kleyner R, et al. (2019): Paralog Studies Augment Gene Discovery: DDX and DHX Genes. American journal of human genetics. 105:302–316.

8. Valentin-Vega YA, Wang YD, Parker M, Patmore DM, Kanagaraj A, Moore J, et al. (2016): Cancer-associated DDX3X mutations drive stress granule assembly and impair global translation. Sci Rep. 6:25996.

9. Soto-Rifo R, Rubilar PS, Limousin T, de Breyne S, Decimo D, Ohlmann T (2012): DEAD-box protein DDX3 associates with eIF4F to promote translation of selected mRNAs. EMBO J. 31:3745–3756.

10. Chen HH, Yu HI, Tarn WY (2016): DDX3 Modulates Neurite Development via Translationally Activating an RNA Regulon Involved in Rac1 Activation. J Neurosci. 36:9792–9804.

11. Garieri M, Stamoulis G, Blanc X, Falconnet E, Ribaux P, Borel C, et al. (2018): Extensive cellular heterogeneity of X inactivation revealed by single-cell allele-specific expression in human fibroblasts. Proc Natl Acad Sci U S A. 115:13015–13020.

12. Tukiainen T, Villani AC, Yen A, Rivas MA, Marshall JL, Satija R, et al. (2017): Landscape of X chromosome inactivation across human tissues. Nature. 550:244–248.

13. Ditton HJ, Zimmer J, Kamp C, Rajpert-De Meyts E, Vogt PH (2004): The AZFa gene DBY (DDX3Y) is widely transcribed but the protein is limited to the male germ cells by translation control. Hum Mol Genet. 13:2333–2341.

14. Foresta C, Ferlin A, Moro E (2000): Deletion and expression analysis of AZFa genes on the human Y chromosome revealed a major role for DBY in male infertility. Hum Mol Genet. 9:1161–1169.

15. Nicola P, Blackburn PR, Rasmussen KJ, Bertsch NL, Klee EW, Hasadsri L, et al. (2019): De novo DDX3X missense variants in males appear viable and contribute to syndromic intellectual disability. American journal of medical genetics Part A. 179:570–578.

16. Kellaris G, Khan K, Baig SM, Tsai IC, Zamora FM, Ruggieri P, et al. (2018): A hypomorphic inherited pathogenic variant in DDX3X causes male intellectual disability with additional neurodevelopmental and neurodegenerative features. Hum Genomics. 12:11.

17. Chen CY, Chan CH, Chen CM, Tsai YS, Tsai TY, Wu Lee YH, et al. (2016): Targeted inactivation of murine Ddx3x: essential roles of Ddx3x in placentation and embryogenesis. Hum Mol Genet. 25:2905–2922.

18. Patmore DM, Jassim A, Nathan E, Gilbertson RJ, Tahan D, Hoffmann N, et al. (2020): DDX3X Suppresses the Susceptibility of Hindbrain Lineages to Medulloblastoma. Dev Cell.

19. Hayashi S, Lewis P, Pevny L, McMahon AP (2002): Efficient gene modulation in mouse epiblast using a Sox2Cre transgenic mouse strain. Mech Dev. 119 Suppl 1:S97–S101.

20. Drapeau E, Riad M, Kajiwara Y, Buxbaum JD (2018): Behavioral Phenotyping of an Improved Mouse Model of Phelan-McDermid Syndrome with a Complete Deletion of the Shank3 Gene. eNeuro. 5.

21. Fox WM (1965): Reflex-ontogeny and behavioural development of the mouse. Anim Behav. 13:234–241.

22. Heyser CJ (2004): Assessment of developmental milestones in rodents. Curr Protoc Neurosci. Chapter 8:Unit 8 18.

23. Nieman BJ, Bock NA, Bishop J, Chen XJ, Sled JG, Rossant J, et al. (2005): Magnetic resonance imaging for detection and analysis of mouse phenotypes. NMR Biomed. 18:447–468.

24. Nieman BJ, Flenniken AM, Adamson SL, Henkelman RM, Sled JG (2006): Anatomical phenotyping in the brain and skull of a mutant mouse by magnetic resonance imaging and computed tomography. Physiol Genomics. 24:154–162.

25. Nieman BJ, Bishop J, Dazai J, Bock NA, Lerch JP, Feintuch A, et al. (2007): MR technology for biological studies in mice. NMR Biomed. 20:291–303.

26. Collins DL, Neelin P, Peters TM, Evans AC (1994): Automatic 3D intersubject registration of MR volumetric data in standardized Talairach space. J Comput Assist Tomogr. 18:192–205.

27. Avants BB, Yushkevich P, Pluta J, Minkoff D, Korczykowski M, Detre J, et al. (2010): The optimal template effect in hippocampus studies of diseased populations. NeuroImage. 49:2457–2466.

28. Lau JC, Lerch JP, Sled JG, Henkelman RM, Evans AC, Bedell BJ (2008): Longitudinal neuroanatomical changes determined by deformation-based morphometry in a mouse model of Alzheimer’s disease. NeuroImage. 42:19–27.

29. Dorr AE, Lerch JP, Spring S, Kabani N, Henkelman RM (2008): High resolution three-dimensional brain atlas using an average magnetic resonance image of 40 adult C57Bl/6J mice. NeuroImage. 42:60–69.

30. Genovese CR, Lazar NA, Nichols T (2002): Thresholding of statistical maps in functional neuroimaging using the false discovery rate. NeuroImage. 15:870–878.

31. Samir P, Kesavardhana S, Patmore DM, Gingras S, Malireddi RKS, Karki R, et al. (2019): DDX3X acts as a live-or-die checkpoint in stressed cells by regulating NLRP3 inflammasome. Nature. 573:590–594.

32. Drapeau E, Dorr NP, Elder GA, Buxbaum JD (2014): Absence of strong strain effects in behavioral analyses of Shank3-deficient mice. Disease models & mechanisms. 7:667–681.

33. Xu C, Krabbe S, Grundemann J, Botta P, Fadok JP, Osakada F, et al. (2016): Distinct Hippocampal Pathways Mediate Dissociable Roles of Context in Memory Retrieval. Cell. 167:961–972 e916.

34. Cai DJ, Aharoni D, Shuman T, Shobe J, Biane J, Song W, et al. (2016): A shared neural ensemble links distinct contextual memories encoded close in time. Nature. 534:115–118.

35. Calhoon GG, Tye KM (2015): Resolving the neural circuits of anxiety. Nat Neurosci. 18:1394–1404.

36. Dominguez MH, Ayoub AE, Rakic P (2013): POU-III transcription factors (Brn1, Brn2, and Oct6) influence neurogenesis, molecular identity, and migratory destination of upper-layer cells of the cerebral cortex. Cereb Cortex. 23:2632–2643.

37. Sugitani Y, Nakai S, Minowa O, Nishi M, Jishage K, Kawano H, et al. (2002): Brn-1 and Brn-2 share crucial roles in the production and positioning of mouse neocortical neurons. Genes Dev. 16:1760–1765.

38. Oishi K, Aramaki M, Nakajima K (2016): Mutually repressive interaction between Brn1/2 and Rorb contributes to the establishment of neocortical layer 2/3 and layer 4. Proc Natl Acad Sci U S A. 113:3371–3376.

39. Alcamo EA, Chirivella L, Dautzenberg M, Dobreva G, Farinas I, Grosschedl R, et al. (2008): Satb2 regulates callosal projection neuron identity in the developing cerebral cortex. Neuron. 57:364–377.

40. Arlotta P, Molyneaux BJ, Chen J, Inoue J, Kominami R, Macklis JD (2005): Neuronal subtype-specific genes that control corticospinal motor neuron development in vivo. Neuron. 45:207–221.

41. Baranek C, Dittrich M, Parthasarathy S, Bonnon CG, Britanova O, Lanshakov D, et al. (2012): Protooncogene Ski cooperates with the chromatin-remodeling factor Satb2 in specifying callosal neurons. Proc Natl Acad Sci U S A. 109:3546–3551.

42. Britanova O, de Juan Romero C, Cheung A, Kwan KY, Schwark M, Gyorgy A, et al. (2008): Satb2 is a postmitotic determinant for upper-layer neuron specification in the neocortex. Neuron. 57:378–392.

43. Srivatsa S, Parthasarathy S, Britanova O, Bormuth I, Donahoo AL, Ackerman SL, et al. (2014): Unc5C and DCC act downstream of Ctip2 and Satb2 and contribute to corpus callosum formation. Nat Commun. 5:3708.

44. Harb K, Magrinelli E, Nicolas CS, Lukianets N, Frangeul L, Pietri M, et al. (2016): Area-specific development of distinct projection neuron subclasses is regulated by postnatal epigenetic modifications. Elife. 5:e09531.

45. Leone DP, Heavner WE, Ferenczi EA, Dobreva G, Huguenard JR, Grosschedl R, et al. (2015): Satb2 Regulates the Differentiation of Both Callosal and Subcerebral Projection Neurons in the Developing Cerebral Cortex. Cereb Cortex. 25:3406–3419.

46. Canovas J, Berndt FA, Sepulveda H, Aguilar R, Veloso FA, Montecino M, et al. (2015): The Specification of Cortical Subcerebral Projection Neurons Depends on the Direct Repression of TBR1 by CTIP1/BCL11a. J Neurosci. 35:7552–7564.

47. Green SA, Hernandez L, Tottenham N, Krasileva K, Bookheimer SY, Dapretto M (2015): Neurobiology of Sensory Overresponsivity in Youth With Autism Spectrum Disorders. JAMA Psychiatry. 72:778–786.

48. He CX, Cantu DA, Mantri SS, Zeiger WA, Goel A, Portera-Cailliau C (2017): Tactile Defensiveness and Impaired Adaptation of Neuronal Activity in the Fmr1 Knock-Out Mouse Model of Autism. J Neurosci. 37:6475–6487.

49. Arroyo ED, Fiole D, Mantri SS, Huang C, Portera-Cailliau C (2019): Dendritic Spines in Early Postnatal Fragile X Mice Are Insensitive to Novel Sensory Experience. J Neurosci. 39:412–419.

50. Chen Q, Deister CA, Gao X, Guo B, Lynn-Jones T, Chen N, et al. (2020): Dysfunction of cortical GABAergic neurons leads to sensory hyper-reactivity in a Shank3 mouse model of ASD. Nat Neurosci. 23:520–532.

51. Orefice LL, Mosko JR, Morency DT, Wells MF, Tasnim A, Mozeika SM, et al. (2019): Targeting Peripheral Somatosensory Neurons to Improve Tactile-Related Phenotypes in ASD Models. Cell. 178:867–886 e824.

52. Orefice LL, Zimmerman AL, Chirila AM, Sleboda SJ, Head JP, Ginty DD (2016): Peripheral Mechanosensory Neuron Dysfunction Underlies Tactile and Behavioral Deficits in Mouse Models of ASDs. Cell. 166:299–313.

53. Hisaoka T, Komori T, Kitamura T, Morikawa Y (2018): Abnormal behaviours relevant to neurodevelopmental disorders in Kirrel3-knockout mice. Sci Rep. 8:1408.

54. Carvell GE, Simons DJ (1996): Abnormal tactile experience early in life disrupts active touch. J Neurosci. 16:2750–2757.

55. Zhang JB, Chen L, Lv ZM, Niu XY, Shao CC, Zhang C, et al. (2016): Oxytocin is implicated in social memory deficits induced by early sensory deprivation in mice. Mol Brain. 9:98.

56. Celikel T, Sakmann B (2007): Sensory integration across space and in time for decision making in the somatosensory system of rodents. Proc Natl Acad Sci U S A. 104:1395–1400.

57. Kuhlman SJ, Olivas ND, Tring E, Ikrar T, Xu X, Trachtenberg JT (2013): A disinhibitory microcircuit initiates critical-period plasticity in the visual cortex. Nature. 501:543–546.

58. Siegel CS, Fink KL, Strittmatter SM, Cafferty WB (2015): Plasticity of intact rubral projections mediates spontaneous recovery of function after corticospinal tract injury. J Neurosci. 35:1443–1457.

59. Kim WB, Cho JH (2020): Encoding of contextual fear memory in hippocampal-amygdala circuit. Nat Commun. 11:1382.

60. den Hoed J, Sollis E, Venselaar H, Estruch SB, Deriziotis P, Fisher SE (2018): Functional characterization of TBR1 variants in neurodevelopmental disorder. Sci Rep. 8:14279.

61. Satterstrom FK, Kosmicki JA, Wang J, Breen MS, De Rubeis S, An JY, et al. (2020): Large-Scale Exome Sequencing Study Implicates Both Developmental and Functional Changes in the Neurobiology of Autism. Cell. 180:568–584 e523.

62. Fazel Darbandi S, Robinson Schwartz SE, Qi Q, Catta-Preta R, Pai EL, Mandell JD, et al. (2018): Neonatal Tbr1 Dosage Controls Cortical Layer 6 Connectivity. Neuron. 100:831–845 e837.

63. Pucilowska J, Vithayathil J, Pagani M, Kelly C, Karlo JC, Robol C, et al. (2018): Pharmacological Inhibition of ERK Signaling Rescues Pathophysiology and Behavioral Phenotype Associated with 16p11.2 Chromosomal Deletion in Mice. J Neurosci. 38:6640–6652.

64. De Rubeis S, He X, Goldberg AP, Poultney CS, Samocha K, Cicek AE, et al. (2014): Synaptic, transcriptional and chromatin genes disrupted in autism. Nature. 515:209–215.

65. Popovitchenko T, Park Y, Page NF, Luo X, Krsnik Z, Liu Y, et al. (2020): Translational derepression of Elavl4 isoforms at their alternative 5′ UTRs determines neuronal development. Nature Communications. 11:1674.

66. Lee JA, Damianov A, Lin CH, Fontes M, Parikshak NN, Anderson ES, et al. (2016): Cytoplasmic Rbfox1 Regulates the Expression of Synaptic and Autism-Related Genes. Neuron. 89:113–128.

67. Gandal MJ, Zhang P, Hadjimichael E, Walker RL, Chen C, Liu S, et al. (2018): Transcriptome-wide isoform-level dysregulation in ASD, schizophrenia, and bipolar disorder. Science. 362.

68. Reijnders MRF, Ansor NM, Kousi M, Yue WW, Tan PL, Clarkson K, et al. (2017): RAC1 Missense Mutations in Developmental Disorders with Diverse Phenotypes. American journal of human genetics. 101:466–477.

69. Zahr SK, Yang G, Kazan H, Borrett MJ, Yuzwa SA, Voronova A, et al. (2018): A Translational Repression Complex in Developing Mammalian Neural Stem Cells that Regulates Neuronal Specification. Neuron. 97:520–537 e526.

70. DeBoer EM, Azevedo R, Vega TA, Brodkin J, Akamatsu W, Okano H, et al. (2014): Prenatal deletion of the RNA-binding protein HuD disrupts postnatal cortical circuit maturation and behavior. J Neurosci. 34:3674–3686.

71. Kraushar ML, Thompson K, Wijeratne HR, Viljetic B, Sakers K, Marson JW, et al. (2014): Temporally defined neocortical translation and polysome assembly are determined by the RNA-binding protein Hu antigen R. Proceedings of the National Academy of Sciences of the United States of America. 111:E3815–3824.

72. Kraushar ML, Viljetic B, Wijeratne HR, Thompson K, Jiao X, Pike JW, et al. (2015): Thalamic WNT3 Secretion Spatiotemporally Regulates the Neocortical Ribosome Signature and mRNA Translation to Specify Neocortical Cell Subtypes. J Neurosci. 35:10911–10926.

73. Popovitchenko T, Thompson K, Viljetic B, Jiao X, Kontonyiannis DL, Kiledjian M, et al. (2016): The RNA binding protein HuR determines the differential translation of autism-associated FoxP subfamily members in the developing neocortex. Sci Rep. 6:28998.

74. Durak O, Gao F, Kaeser-Woo YJ, Rueda R, Martorell AJ, Nott A, et al. (2016): Chd8 mediates cortical neurogenesis via transcriptional regulation of cell cycle and Wnt signaling. Nature neuroscience. 19:1477–1488.

75. Xu Q, Liu YY, Wang X, Tan GH, Li HP, Hulbert SW, et al. (2018): Autism-associated CHD8 deficiency impairs axon development and migration of cortical neurons. Mol Autism. 9:65.

76. Kaplanis J, Samocha KE, Wiel L, Zhang Z, Arvai KJ, Eberhardt RY, et al. (2020): Evidence for 28 genetic disorders discovered by combining healthcare and research data. Nature.

