## Supplemental Tables for "Developmental and behavioral phenotypes in a new mouse model of DDX3X syndrome"

### Slide 1
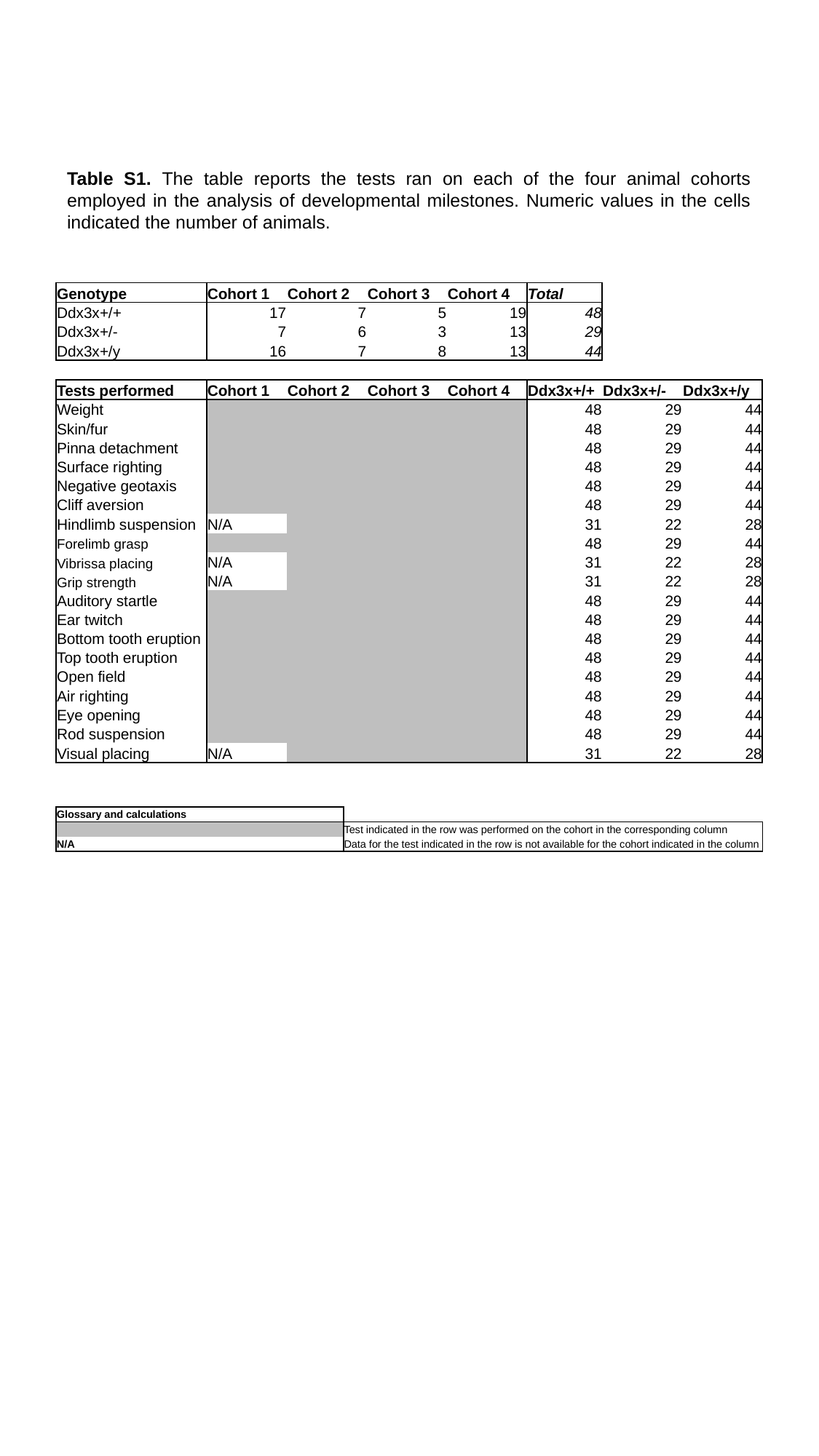

Table S1. The table reports the tests ran on each of the four animal cohorts employed in the analysis of developmental milestones. Numeric values in the cells indicated the number of animals.
| Genotype | Cohort 1 | Cohort 2 | Cohort 3 | Cohort 4 | Total | | |
| --- | --- | --- | --- | --- | --- | --- | --- |
| Ddx3x+/+ | 17 | 7 | 5 | 19 | 48 | | |
| Ddx3x+/- | 7 | 6 | 3 | 13 | 29 | | |
| Ddx3x+/y | 16 | 7 | 8 | 13 | 44 | | |
| Tests performed | Cohort 1 | Cohort 2 | Cohort 3 | Cohort 4 | Ddx3x+/+ | Ddx3x+/- | Ddx3x+/y |
| Weight | | | | | 48 | 29 | 44 |
| Skin/fur | | | | | 48 | 29 | 44 |
| Pinna detachment | | | | | 48 | 29 | 44 |
| Surface righting | | | | | 48 | 29 | 44 |
| Negative geotaxis | | | | | 48 | 29 | 44 |
| Cliff aversion | | | | | 48 | 29 | 44 |
| Hindlimb suspension | N/A | | | | 31 | 22 | 28 |
| Forelimb grasp | | | | | 48 | 29 | 44 |
| Vibrissa placing | N/A | | | | 31 | 22 | 28 |
| Grip strength | N/A | | | | 31 | 22 | 28 |
| Auditory startle | | | | | 48 | 29 | 44 |
| Ear twitch | | | | | 48 | 29 | 44 |
| Bottom tooth eruption | | | | | 48 | 29 | 44 |
| Top tooth eruption | | | | | 48 | 29 | 44 |
| Open field | | | | | 48 | 29 | 44 |
| Air righting | | | | | 48 | 29 | 44 |
| Eye opening | | | | | 48 | 29 | 44 |
| Rod suspension | | | | | 48 | 29 | 44 |
| Visual placing | N/A | | | | 31 | 22 | 28 |
| Glossary and calculations | |
| --- | --- |
| | Test indicated in the row was performed on the cohort in the corresponding column |
| N/A | Data for the test indicated in the row is not available for the cohort indicated in the column |

### Slide 2
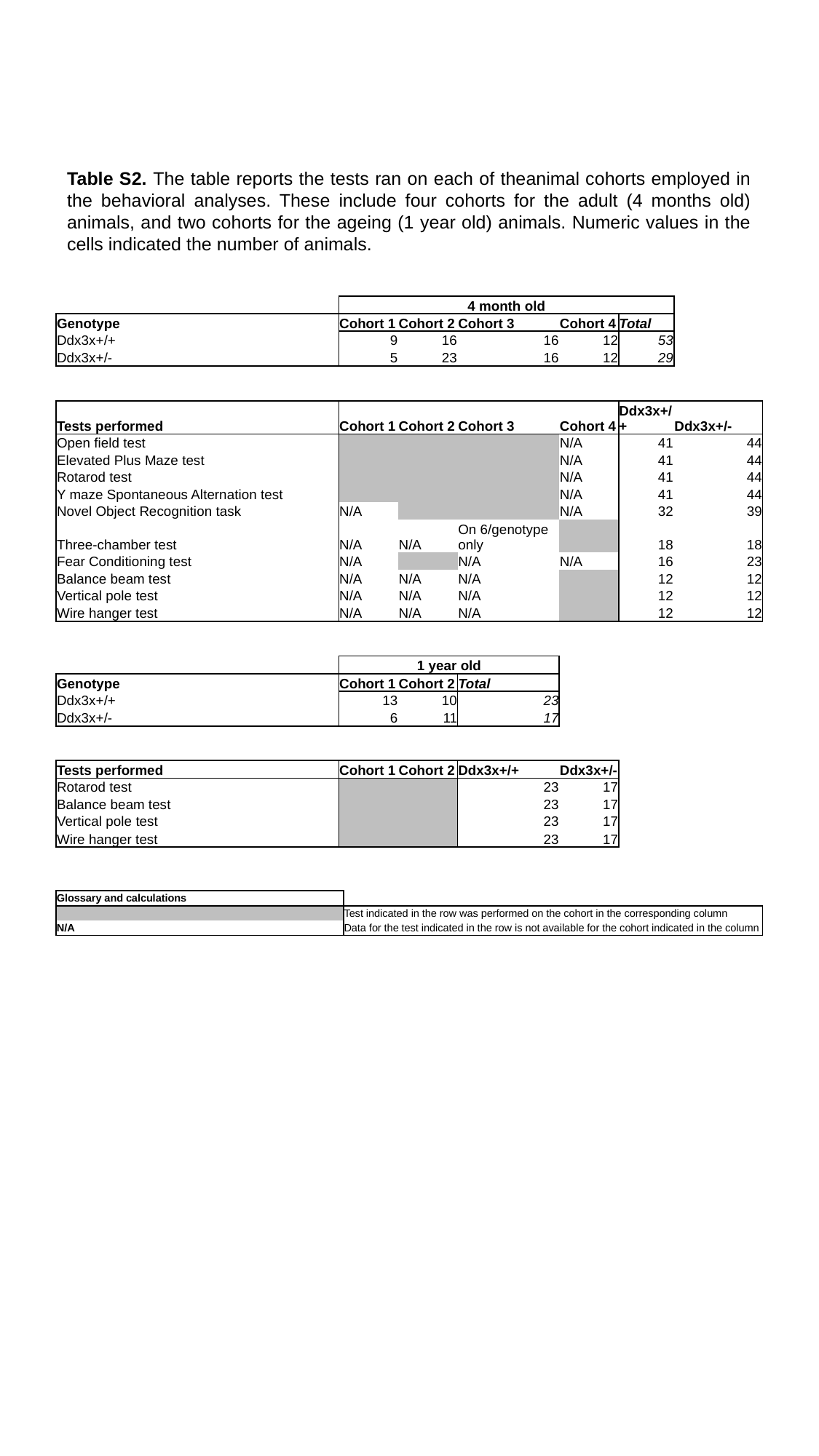

Table S2. The table reports the tests ran on each of theanimal cohorts employed in the behavioral analyses. These include four cohorts for the adult (4 months old) animals, and two cohorts for the ageing (1 year old) animals. Numeric values in the cells indicated the number of animals.
| | 4 month old | | | | | |
| --- | --- | --- | --- | --- | --- | --- |
| Genotype | Cohort 1 | Cohort 2 | Cohort 3 | Cohort 4 | Total | |
| Ddx3x+/+ | 9 | 16 | 16 | 12 | 53 | |
| Ddx3x+/- | 5 | 23 | 16 | 12 | 29 | |
| Tests performed | Cohort 1 | Cohort 2 | Cohort 3 | Cohort 4 | Ddx3x+/+ | Ddx3x+/- |
| Open field test | | | | N/A | 41 | 44 |
| Elevated Plus Maze test | | | | N/A | 41 | 44 |
| Rotarod test | | | | N/A | 41 | 44 |
| Y maze Spontaneous Alternation test | | | | N/A | 41 | 44 |
| Novel Object Recognition task | N/A | | | N/A | 32 | 39 |
| Three-chamber test | N/A | N/A | On 6/genotype only | | 18 | 18 |
| Fear Conditioning test | N/A | | N/A | N/A | 16 | 23 |
| Balance beam test | N/A | N/A | N/A | | 12 | 12 |
| Vertical pole test | N/A | N/A | N/A | | 12 | 12 |
| Wire hanger test | N/A | N/A | N/A | | 12 | 12 |
| | 1 year old | | | | | |
| Genotype | Cohort 1 | Cohort 2 | Total | | | |
| Ddx3x+/+ | 13 | 10 | 23 | | | |
| Ddx3x+/- | 6 | 11 | 17 | | | |
| Tests performed | Cohort 1 | Cohort 2 | Ddx3x+/+ | Ddx3x+/- | | |
| Rotarod test | | | 23 | 17 | | |
| Balance beam test | | | 23 | 17 | | |
| Vertical pole test | | | 23 | 17 | | |
| Wire hanger test | | | 23 | 17 | | |
| Glossary and calculations | |
| --- | --- |
| | Test indicated in the row was performed on the cohort in the corresponding column |
| N/A | Data for the test indicated in the row is not available for the cohort indicated in the column |

### Slide 3
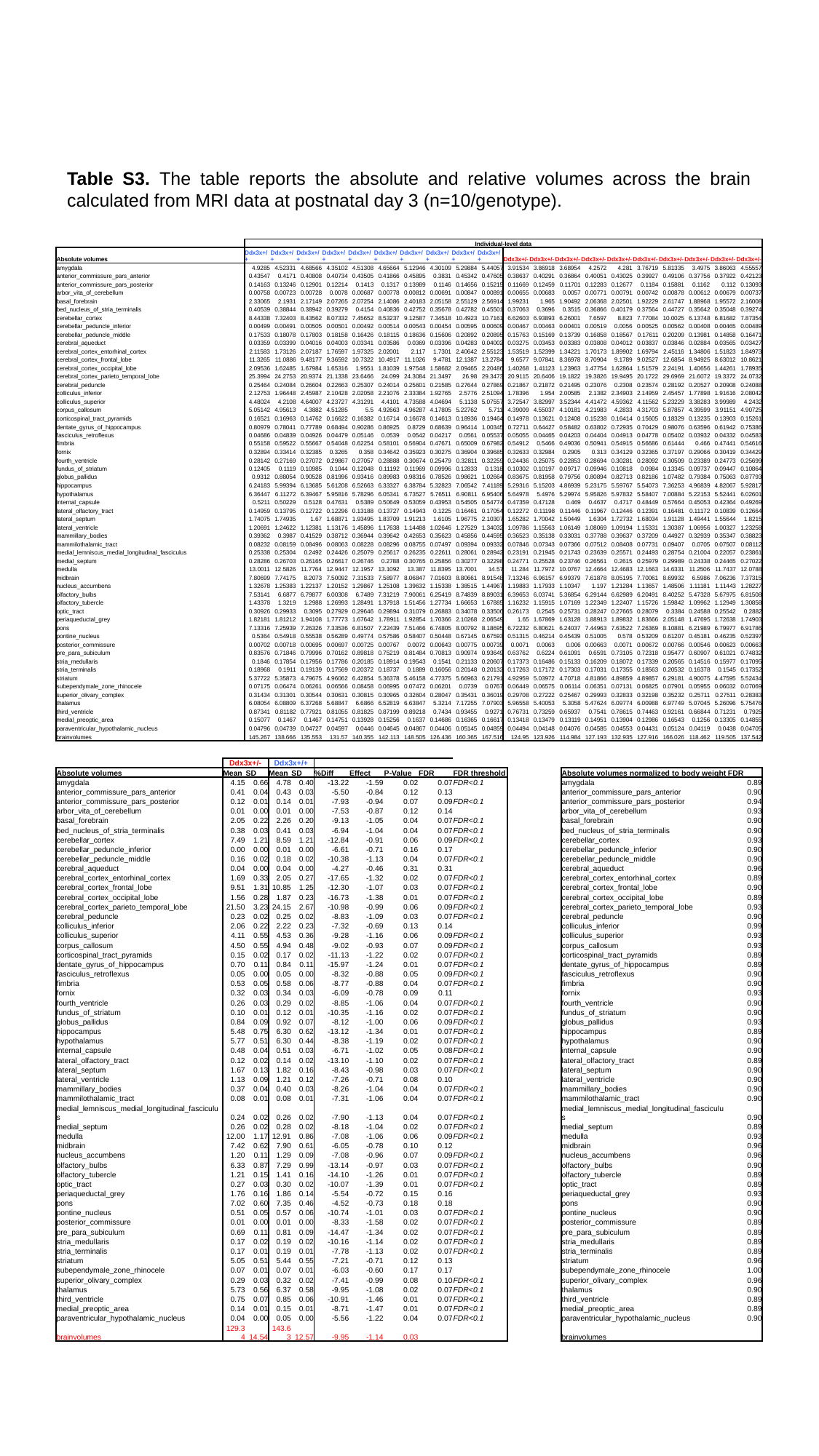

Table S3. The table reports the absolute and relative volumes across the brain calculated from MRI data at postnatal day 3 (n=10/genotype).
| | Individual-level data | | | | | | | | | | | | | | | | | | | |
| --- | --- | --- | --- | --- | --- | --- | --- | --- | --- | --- | --- | --- | --- | --- | --- | --- | --- | --- | --- | --- |
| Absolute volumes | Ddx3x+/+ | Ddx3x+/+ | Ddx3x+/+ | Ddx3x+/+ | Ddx3x+/+ | Ddx3x+/+ | Ddx3x+/+ | Ddx3x+/+ | Ddx3x+/+ | Ddx3x+/+ | Ddx3x+/- | Ddx3x+/- | Ddx3x+/- | Ddx3x+/- | Ddx3x+/- | Ddx3x+/- | Ddx3x+/- | Ddx3x+/- | Ddx3x+/- | Ddx3x+/- |
| amygdala | 4.9285 | 4.52331 | 4.68566 | 4.35102 | 4.51308 | 4.65664 | 5.12946 | 4.30109 | 5.29884 | 5.44057 | 3.91534 | 3.86918 | 3.68954 | 4.2572 | 4.281 | 3.76719 | 5.81335 | 3.4975 | 3.86063 | 4.55557 |
| anterior\_commissure\_pars\_anterior | 0.43547 | 0.4171 | 0.40808 | 0.40734 | 0.43505 | 0.41866 | 0.45895 | 0.3831 | 0.45342 | 0.47605 | 0.38637 | 0.40291 | 0.36864 | 0.40051 | 0.43025 | 0.39927 | 0.49106 | 0.37756 | 0.37922 | 0.42123 |
| anterior\_commissure\_pars\_posterior | 0.14163 | 0.13246 | 0.12901 | 0.12214 | 0.1413 | 0.1317 | 0.13989 | 0.1146 | 0.14656 | 0.15215 | 0.11669 | 0.12459 | 0.11701 | 0.12283 | 0.12677 | 0.1184 | 0.15881 | 0.1162 | 0.112 | 0.13093 |
| arbor\_vita\_of\_cerebellum | 0.00758 | 0.00723 | 0.00728 | 0.0078 | 0.00687 | 0.00778 | 0.00812 | 0.00691 | 0.00847 | 0.00891 | 0.00655 | 0.00683 | 0.0057 | 0.00771 | 0.00791 | 0.00742 | 0.00878 | 0.00612 | 0.00679 | 0.00737 |
| basal\_forebrain | 2.33065 | 2.1931 | 2.17149 | 2.07265 | 2.07254 | 2.14086 | 2.40183 | 2.05158 | 2.55129 | 2.56914 | 1.99231 | 1.965 | 1.90492 | 2.06368 | 2.02501 | 1.92229 | 2.61747 | 1.88968 | 1.95572 | 2.16008 |
| bed\_nucleus\_of\_stria\_terminalis | 0.40539 | 0.38844 | 0.38942 | 0.39279 | 0.4154 | 0.40836 | 0.42752 | 0.35678 | 0.42782 | 0.45501 | 0.37063 | 0.3696 | 0.3515 | 0.36866 | 0.40179 | 0.37564 | 0.44727 | 0.35642 | 0.35048 | 0.39274 |
| cerebellar\_cortex | 8.44338 | 7.32403 | 8.43562 | 8.07332 | 7.45652 | 8.53237 | 9.12587 | 7.34518 | 10.4923 | 10.7161 | 6.62603 | 6.93893 | 6.26001 | 7.6597 | 8.823 | 7.77084 | 10.0025 | 6.13748 | 6.81682 | 7.87354 |
| cerebellar\_peduncle\_inferior | 0.00499 | 0.00491 | 0.00505 | 0.00501 | 0.00492 | 0.00514 | 0.00543 | 0.00454 | 0.00595 | 0.00609 | 0.00467 | 0.00463 | 0.00401 | 0.00519 | 0.0056 | 0.00525 | 0.00562 | 0.00408 | 0.00465 | 0.00489 |
| cerebellar\_peduncle\_middle | 0.17533 | 0.18078 | 0.17803 | 0.18158 | 0.16426 | 0.18115 | 0.18636 | 0.15606 | 0.20892 | 0.20895 | 0.15763 | 0.15169 | 0.13739 | 0.16858 | 0.18567 | 0.17611 | 0.20209 | 0.13981 | 0.14858 | 0.16471 |
| cerebral\_aqueduct | 0.03359 | 0.03399 | 0.04016 | 0.04003 | 0.03341 | 0.03586 | 0.0369 | 0.03396 | 0.04283 | 0.04002 | 0.03275 | 0.03453 | 0.03383 | 0.03808 | 0.04012 | 0.03837 | 0.03846 | 0.02884 | 0.03565 | 0.03427 |
| cerebral\_cortex\_entorhinal\_cortex | 2.11583 | 1.73126 | 2.07187 | 1.76597 | 1.97325 | 2.02001 | 2.117 | 1.7301 | 2.40642 | 2.55123 | 1.53519 | 1.52399 | 1.34221 | 1.70173 | 1.89902 | 1.69794 | 2.45116 | 1.34806 | 1.51823 | 1.84973 |
| cerebral\_cortex\_frontal\_lobe | 11.3265 | 11.0886 | 9.48177 | 9.36592 | 10.7322 | 10.4917 | 11.1026 | 9.4781 | 12.1387 | 13.2784 | 9.6577 | 9.07841 | 8.36978 | 8.70904 | 9.1789 | 9.02527 | 12.6854 | 8.94925 | 8.63012 | 10.8621 |
| cerebral\_cortex\_occipital\_lobe | 2.09536 | 1.62485 | 1.67984 | 1.65316 | 1.9551 | 1.81039 | 1.97548 | 1.58682 | 2.09465 | 2.20486 | 1.40268 | 1.41123 | 1.23963 | 1.47754 | 1.62864 | 1.51579 | 2.24191 | 1.40656 | 1.44261 | 1.78935 |
| cerebral\_cortex\_parieto\_temporal\_lobe | 25.3994 | 24.2753 | 20.9374 | 21.1338 | 23.6466 | 24.099 | 24.3084 | 21.3497 | 26.98 | 29.3473 | 20.9115 | 20.6406 | 19.1822 | 19.3826 | 19.9495 | 20.1722 | 29.6969 | 21.6072 | 19.3372 | 24.0732 |
| cerebral\_peduncle | 0.25464 | 0.24084 | 0.26604 | 0.22663 | 0.25307 | 0.24014 | 0.25601 | 0.21585 | 0.27644 | 0.27869 | 0.21867 | 0.21872 | 0.21495 | 0.23076 | 0.2308 | 0.23574 | 0.28192 | 0.20527 | 0.20908 | 0.24088 |
| colliculus\_inferior | 2.12753 | 1.96448 | 2.45987 | 2.10428 | 2.02058 | 2.21076 | 2.33384 | 1.92765 | 2.5776 | 2.51094 | 1.78396 | 1.954 | 2.00585 | 2.1382 | 2.34903 | 2.14959 | 2.45457 | 1.77898 | 1.91616 | 2.08042 |
| colliculus\_superior | 4.48024 | 4.2108 | 4.64007 | 4.23727 | 4.31291 | 4.4101 | 4.73588 | 4.04694 | 5.1138 | 5.07557 | 3.72547 | 3.82997 | 3.52344 | 4.41472 | 4.59362 | 4.11562 | 5.23229 | 3.38283 | 3.99989 | 4.2432 |
| corpus\_callosum | 5.05142 | 4.95613 | 4.3882 | 4.51285 | 5.5 | 4.92663 | 4.96287 | 4.17805 | 5.22762 | 5.711 | 4.39009 | 4.55037 | 4.10181 | 4.21983 | 4.2833 | 4.31703 | 5.87857 | 4.39599 | 3.91151 | 4.90725 |
| corticospinal\_tract\_pyramids | 0.16521 | 0.16963 | 0.14762 | 0.16622 | 0.16382 | 0.16714 | 0.16678 | 0.14613 | 0.18936 | 0.19464 | 0.14978 | 0.13621 | 0.12408 | 0.15238 | 0.16414 | 0.15605 | 0.18329 | 0.13235 | 0.13903 | 0.15261 |
| dentate\_gyrus\_of\_hippocampus | 0.80979 | 0.78041 | 0.77789 | 0.68494 | 0.90286 | 0.86925 | 0.8729 | 0.68639 | 0.96414 | 1.00345 | 0.72711 | 0.64427 | 0.58482 | 0.63802 | 0.72935 | 0.70429 | 0.98076 | 0.63596 | 0.61942 | 0.75386 |
| fasciculus\_retroflexus | 0.04686 | 0.04839 | 0.04926 | 0.04479 | 0.05146 | 0.0539 | 0.0542 | 0.04217 | 0.0561 | 0.05537 | 0.05055 | 0.04465 | 0.04203 | 0.04404 | 0.04913 | 0.04778 | 0.05402 | 0.03932 | 0.04332 | 0.04583 |
| fimbria | 0.55158 | 0.59522 | 0.55667 | 0.54048 | 0.62254 | 0.58101 | 0.56904 | 0.47671 | 0.65009 | 0.67982 | 0.54912 | 0.5466 | 0.49036 | 0.50941 | 0.54915 | 0.56686 | 0.61444 | 0.466 | 0.47441 | 0.54616 |
| fornix | 0.32894 | 0.33414 | 0.32385 | 0.3265 | 0.358 | 0.34642 | 0.35923 | 0.30275 | 0.36904 | 0.39685 | 0.32633 | 0.32984 | 0.2905 | 0.313 | 0.34129 | 0.32365 | 0.37197 | 0.29066 | 0.30419 | 0.34429 |
| fourth\_ventricle | 0.28142 | 0.27169 | 0.27072 | 0.29867 | 0.27057 | 0.28888 | 0.30674 | 0.25479 | 0.32811 | 0.32259 | 0.24436 | 0.25075 | 0.22853 | 0.28694 | 0.30281 | 0.28092 | 0.30509 | 0.23389 | 0.24773 | 0.25699 |
| fundus\_of\_striatum | 0.12405 | 0.1119 | 0.10985 | 0.1044 | 0.12048 | 0.11192 | 0.11969 | 0.09996 | 0.12833 | 0.1318 | 0.10302 | 0.10197 | 0.09717 | 0.09946 | 0.10818 | 0.0984 | 0.13345 | 0.09737 | 0.09447 | 0.10864 |
| globus\_pallidus | 0.9312 | 0.88054 | 0.90528 | 0.81996 | 0.93416 | 0.89983 | 0.98316 | 0.78526 | 0.98621 | 1.02664 | 0.83675 | 0.81958 | 0.79756 | 0.80894 | 0.82713 | 0.82186 | 1.07482 | 0.79384 | 0.75063 | 0.87793 |
| hippocampus | 6.24183 | 5.99394 | 6.13685 | 5.61208 | 6.52663 | 6.33327 | 6.38784 | 5.32823 | 7.06542 | 7.41189 | 5.29316 | 5.15203 | 4.86939 | 5.23175 | 5.59767 | 5.54073 | 7.36253 | 4.96839 | 4.82067 | 5.92817 |
| hypothalamus | 6.36447 | 6.11272 | 6.39467 | 5.95816 | 5.78296 | 6.05341 | 6.73527 | 5.76511 | 6.90811 | 6.95406 | 5.64978 | 5.4976 | 5.29974 | 5.95826 | 5.97832 | 5.58407 | 7.00884 | 5.22153 | 5.52441 | 6.02601 |
| internal\_capsule | 0.5211 | 0.50229 | 0.5128 | 0.47631 | 0.5389 | 0.50649 | 0.53059 | 0.43953 | 0.54505 | 0.54774 | 0.47359 | 0.47128 | 0.469 | 0.4637 | 0.4717 | 0.48449 | 0.57664 | 0.45053 | 0.42364 | 0.49269 |
| lateral\_olfactory\_tract | 0.14959 | 0.13795 | 0.12722 | 0.12296 | 0.13188 | 0.13727 | 0.14943 | 0.1225 | 0.16461 | 0.17054 | 0.12272 | 0.11198 | 0.11446 | 0.11967 | 0.12446 | 0.12391 | 0.16481 | 0.11172 | 0.10839 | 0.12664 |
| lateral\_septum | 1.74075 | 1.74935 | 1.67 | 1.68871 | 1.93495 | 1.83709 | 1.91213 | 1.6105 | 1.96775 | 2.10307 | 1.65282 | 1.70042 | 1.50449 | 1.6304 | 1.72732 | 1.68034 | 1.91128 | 1.49441 | 1.55644 | 1.8215 |
| lateral\_ventricle | 1.20691 | 1.24622 | 1.12381 | 1.13176 | 1.45896 | 1.17638 | 1.14488 | 1.02646 | 1.27529 | 1.34032 | 1.09786 | 1.15563 | 1.06149 | 1.08069 | 1.09194 | 1.15331 | 1.30387 | 1.06956 | 1.00327 | 1.23258 |
| mammillary\_bodies | 0.39362 | 0.3987 | 0.41529 | 0.38712 | 0.36944 | 0.39642 | 0.42653 | 0.35623 | 0.45856 | 0.44595 | 0.36523 | 0.35138 | 0.33031 | 0.37788 | 0.39637 | 0.37209 | 0.44927 | 0.32939 | 0.35347 | 0.38823 |
| mammilothalamic\_tract | 0.08232 | 0.08159 | 0.08496 | 0.08063 | 0.08228 | 0.08296 | 0.08755 | 0.07497 | 0.09394 | 0.09332 | 0.07846 | 0.07343 | 0.07366 | 0.07512 | 0.08408 | 0.07731 | 0.09407 | 0.0705 | 0.07507 | 0.08112 |
| medial\_lemniscus\_medial\_longitudinal\_fasciculus | 0.25338 | 0.25304 | 0.2492 | 0.24426 | 0.25079 | 0.25617 | 0.26235 | 0.22611 | 0.28061 | 0.28942 | 0.23191 | 0.21945 | 0.21743 | 0.23639 | 0.25571 | 0.24493 | 0.28754 | 0.21004 | 0.22057 | 0.23861 |
| medial\_septum | 0.28286 | 0.26703 | 0.26165 | 0.26617 | 0.26746 | 0.2788 | 0.30765 | 0.25856 | 0.30277 | 0.32298 | 0.24771 | 0.25528 | 0.23746 | 0.26561 | 0.2615 | 0.25979 | 0.29989 | 0.24338 | 0.24465 | 0.27022 |
| medulla | 13.0011 | 12.5826 | 11.7764 | 12.9447 | 12.1957 | 13.1092 | 13.387 | 11.8395 | 13.7001 | 14.57 | 11.284 | 11.7972 | 10.0767 | 12.4664 | 12.4683 | 12.1663 | 14.6331 | 11.2506 | 11.7437 | 12.0788 |
| midbrain | 7.80699 | 7.74175 | 8.2073 | 7.50092 | 7.31533 | 7.58977 | 8.06847 | 7.01603 | 8.80661 | 8.91548 | 7.13246 | 6.96157 | 6.99379 | 7.61878 | 8.05195 | 7.70061 | 8.69932 | 6.5986 | 7.06236 | 7.37315 |
| nucleus\_accumbens | 1.32678 | 1.25383 | 1.22137 | 1.20152 | 1.29867 | 1.25108 | 1.39632 | 1.15338 | 1.38515 | 1.44967 | 1.19883 | 1.17933 | 1.10347 | 1.197 | 1.21284 | 1.13657 | 1.48506 | 1.11181 | 1.11443 | 1.28227 |
| olfactory\_bulbs | 7.53141 | 6.6877 | 6.79877 | 6.00308 | 6.7489 | 7.31219 | 7.90061 | 6.25419 | 8.74839 | 8.89031 | 6.39653 | 6.03741 | 5.36854 | 6.29144 | 6.62989 | 6.20491 | 8.40252 | 5.47328 | 5.67975 | 6.81508 |
| olfactory\_tubercle | 1.43378 | 1.3219 | 1.2988 | 1.26993 | 1.28491 | 1.37918 | 1.51456 | 1.27734 | 1.66653 | 1.67885 | 1.16232 | 1.15915 | 1.07169 | 1.22349 | 1.22407 | 1.15726 | 1.59842 | 1.09962 | 1.12949 | 1.30858 |
| optic\_tract | 0.30926 | 0.29933 | 0.3095 | 0.27929 | 0.29646 | 0.29894 | 0.31079 | 0.26883 | 0.34078 | 0.33506 | 0.26173 | 0.2545 | 0.25731 | 0.28247 | 0.27665 | 0.28079 | 0.3384 | 0.24588 | 0.25542 | 0.2882 |
| periaqueductal\_grey | 1.82181 | 1.81212 | 1.94108 | 1.77773 | 1.67642 | 1.78911 | 1.92854 | 1.70366 | 2.10268 | 2.06549 | 1.65 | 1.67869 | 1.63128 | 1.88913 | 1.89832 | 1.83666 | 2.05148 | 1.47695 | 1.72638 | 1.74903 |
| pons | 7.13316 | 7.25939 | 7.26326 | 7.33536 | 6.81507 | 7.22439 | 7.51466 | 6.74805 | 8.00792 | 8.18695 | 6.72232 | 6.80621 | 6.24037 | 7.44963 | 7.63522 | 7.26369 | 8.10881 | 6.21989 | 6.79977 | 6.91786 |
| pontine\_nucleus | 0.5364 | 0.54918 | 0.55538 | 0.56289 | 0.49774 | 0.57586 | 0.58407 | 0.50448 | 0.67145 | 0.67593 | 0.51315 | 0.46214 | 0.45439 | 0.51005 | 0.578 | 0.53209 | 0.61207 | 0.45181 | 0.46235 | 0.52397 |
| posterior\_commissure | 0.00702 | 0.00718 | 0.00695 | 0.00697 | 0.00725 | 0.00767 | 0.0072 | 0.00643 | 0.00775 | 0.00739 | 0.0071 | 0.0063 | 0.006 | 0.00663 | 0.0071 | 0.00672 | 0.00766 | 0.00546 | 0.00623 | 0.00663 |
| pre\_para\_subiculum | 0.83576 | 0.71846 | 0.79996 | 0.70162 | 0.89818 | 0.75219 | 0.81484 | 0.70813 | 0.90974 | 0.93649 | 0.63762 | 0.6224 | 0.61091 | 0.6591 | 0.73105 | 0.72318 | 0.95477 | 0.60907 | 0.61021 | 0.74832 |
| stria\_medullaris | 0.1846 | 0.17854 | 0.17956 | 0.17786 | 0.20185 | 0.18914 | 0.19543 | 0.1541 | 0.21133 | 0.20607 | 0.17373 | 0.16486 | 0.15133 | 0.16209 | 0.18072 | 0.17339 | 0.20565 | 0.14516 | 0.15977 | 0.17095 |
| stria\_terminalis | 0.18968 | 0.1911 | 0.19139 | 0.17569 | 0.20372 | 0.18737 | 0.1889 | 0.16056 | 0.20148 | 0.20132 | 0.17263 | 0.17172 | 0.17303 | 0.17031 | 0.17355 | 0.18563 | 0.20532 | 0.16378 | 0.1545 | 0.17352 |
| striatum | 5.37722 | 5.35873 | 4.79675 | 4.96062 | 6.42854 | 5.36378 | 5.46158 | 4.77375 | 5.66963 | 6.21791 | 4.92959 | 5.03972 | 4.70718 | 4.81866 | 4.89859 | 4.89857 | 6.29181 | 4.90075 | 4.47595 | 5.52434 |
| subependymale\_zone\_rhinocele | 0.07175 | 0.06474 | 0.06261 | 0.06566 | 0.08458 | 0.06995 | 0.07472 | 0.06201 | 0.0739 | 0.0767 | 0.06449 | 0.06575 | 0.06114 | 0.06351 | 0.07131 | 0.06825 | 0.07901 | 0.05955 | 0.06032 | 0.07069 |
| superior\_olivary\_complex | 0.31434 | 0.31301 | 0.30544 | 0.30631 | 0.30815 | 0.30965 | 0.32604 | 0.28047 | 0.35431 | 0.36019 | 0.29708 | 0.27222 | 0.25467 | 0.29993 | 0.32833 | 0.32198 | 0.35232 | 0.25711 | 0.27511 | 0.28383 |
| thalamus | 6.08054 | 6.08809 | 6.37268 | 5.68847 | 6.6866 | 6.52819 | 6.63847 | 5.3214 | 7.17255 | 7.07903 | 5.96558 | 5.40053 | 5.3058 | 5.47624 | 6.09774 | 6.00988 | 6.97749 | 5.07045 | 5.26096 | 5.75476 |
| third\_ventricle | 0.87341 | 0.81182 | 0.77921 | 0.81055 | 0.81825 | 0.87199 | 0.89218 | 0.7434 | 0.93455 | 0.9271 | 0.76731 | 0.73259 | 0.65937 | 0.7541 | 0.78615 | 0.74463 | 0.92161 | 0.66844 | 0.71231 | 0.7925 |
| medial\_preoptic\_area | 0.15077 | 0.1467 | 0.1467 | 0.14751 | 0.13928 | 0.15256 | 0.1637 | 0.14686 | 0.16365 | 0.16617 | 0.13418 | 0.13479 | 0.13119 | 0.14951 | 0.13904 | 0.12986 | 0.16543 | 0.1256 | 0.13305 | 0.14855 |
| paraventricular\_hypothalamic\_nucleus | 0.04796 | 0.04739 | 0.04727 | 0.04597 | 0.0446 | 0.04645 | 0.04867 | 0.04406 | 0.05145 | 0.04859 | 0.04494 | 0.04148 | 0.04076 | 0.04585 | 0.04553 | 0.04431 | 0.05124 | 0.04119 | 0.0438 | 0.04705 |
| brainvolumes | 145.267 | 138.666 | 135.553 | 131.57 | 140.355 | 142.113 | 148.505 | 126.436 | 160.365 | 167.516 | 124.95 | 123.926 | 114.984 | 127.193 | 132.935 | 127.916 | 166.026 | 118.462 | 119.505 | 137.542 |
| | Ddx3x+/- | | Ddx3x+/+ | | | | | | | | | |
| --- | --- | --- | --- | --- | --- | --- | --- | --- | --- | --- | --- | --- |
| Absolute volumes | Mean | SD | Mean | SD | %Diff | Effect | P-Value | FDR | FDR threshold | | Absolute volumes normalized to body weight | FDR |
| amygdala | 4.15 | 0.66 | 4.78 | 0.40 | -13.22 | -1.59 | 0.02 | 0.07 | FDR<0.1 | | amygdala | 0.89 |
| anterior\_commissure\_pars\_anterior | 0.41 | 0.04 | 0.43 | 0.03 | -5.50 | -0.84 | 0.12 | 0.13 | | | anterior\_commissure\_pars\_anterior | 0.90 |
| anterior\_commissure\_pars\_posterior | 0.12 | 0.01 | 0.14 | 0.01 | -7.93 | -0.94 | 0.07 | 0.09 | FDR<0.1 | | anterior\_commissure\_pars\_posterior | 0.94 |
| arbor\_vita\_of\_cerebellum | 0.01 | 0.00 | 0.01 | 0.00 | -7.53 | -0.87 | 0.12 | 0.14 | | | arbor\_vita\_of\_cerebellum | 0.93 |
| basal\_forebrain | 2.05 | 0.22 | 2.26 | 0.20 | -9.13 | -1.05 | 0.04 | 0.07 | FDR<0.1 | | basal\_forebrain | 0.90 |
| bed\_nucleus\_of\_stria\_terminalis | 0.38 | 0.03 | 0.41 | 0.03 | -6.94 | -1.04 | 0.04 | 0.07 | FDR<0.1 | | bed\_nucleus\_of\_stria\_terminalis | 0.90 |
| cerebellar\_cortex | 7.49 | 1.21 | 8.59 | 1.21 | -12.84 | -0.91 | 0.06 | 0.09 | FDR<0.1 | | cerebellar\_cortex | 0.93 |
| cerebellar\_peduncle\_inferior | 0.00 | 0.00 | 0.01 | 0.00 | -6.61 | -0.71 | 0.16 | 0.17 | | | cerebellar\_peduncle\_inferior | 0.90 |
| cerebellar\_peduncle\_middle | 0.16 | 0.02 | 0.18 | 0.02 | -10.38 | -1.13 | 0.04 | 0.07 | FDR<0.1 | | cerebellar\_peduncle\_middle | 0.90 |
| cerebral\_aqueduct | 0.04 | 0.00 | 0.04 | 0.00 | -4.27 | -0.46 | 0.31 | 0.31 | | | cerebral\_aqueduct | 0.96 |
| cerebral\_cortex\_entorhinal\_cortex | 1.69 | 0.33 | 2.05 | 0.27 | -17.65 | -1.32 | 0.02 | 0.07 | FDR<0.1 | | cerebral\_cortex\_entorhinal\_cortex | 0.89 |
| cerebral\_cortex\_frontal\_lobe | 9.51 | 1.31 | 10.85 | 1.25 | -12.30 | -1.07 | 0.03 | 0.07 | FDR<0.1 | | cerebral\_cortex\_frontal\_lobe | 0.90 |
| cerebral\_cortex\_occipital\_lobe | 1.56 | 0.28 | 1.87 | 0.23 | -16.73 | -1.38 | 0.01 | 0.07 | FDR<0.1 | | cerebral\_cortex\_occipital\_lobe | 0.89 |
| cerebral\_cortex\_parieto\_temporal\_lobe | 21.50 | 3.23 | 24.15 | 2.67 | -10.98 | -0.99 | 0.06 | 0.09 | FDR<0.1 | | cerebral\_cortex\_parieto\_temporal\_lobe | 0.93 |
| cerebral\_peduncle | 0.23 | 0.02 | 0.25 | 0.02 | -8.83 | -1.09 | 0.03 | 0.07 | FDR<0.1 | | cerebral\_peduncle | 0.90 |
| colliculus\_inferior | 2.06 | 0.22 | 2.22 | 0.23 | -7.32 | -0.69 | 0.13 | 0.14 | | | colliculus\_inferior | 0.99 |
| colliculus\_superior | 4.11 | 0.55 | 4.53 | 0.36 | -9.28 | -1.16 | 0.06 | 0.09 | FDR<0.1 | | colliculus\_superior | 0.93 |
| corpus\_callosum | 4.50 | 0.55 | 4.94 | 0.48 | -9.02 | -0.93 | 0.07 | 0.09 | FDR<0.1 | | corpus\_callosum | 0.93 |
| corticospinal\_tract\_pyramids | 0.15 | 0.02 | 0.17 | 0.02 | -11.13 | -1.22 | 0.02 | 0.07 | FDR<0.1 | | corticospinal\_tract\_pyramids | 0.89 |
| dentate\_gyrus\_of\_hippocampus | 0.70 | 0.11 | 0.84 | 0.11 | -15.97 | -1.24 | 0.01 | 0.07 | FDR<0.1 | | dentate\_gyrus\_of\_hippocampus | 0.89 |
| fasciculus\_retroflexus | 0.05 | 0.00 | 0.05 | 0.00 | -8.32 | -0.88 | 0.05 | 0.09 | FDR<0.1 | | fasciculus\_retroflexus | 0.90 |
| fimbria | 0.53 | 0.05 | 0.58 | 0.06 | -8.77 | -0.88 | 0.04 | 0.07 | FDR<0.1 | | fimbria | 0.90 |
| fornix | 0.32 | 0.03 | 0.34 | 0.03 | -6.09 | -0.78 | 0.09 | 0.11 | | | fornix | 0.93 |
| fourth\_ventricle | 0.26 | 0.03 | 0.29 | 0.02 | -8.85 | -1.06 | 0.04 | 0.07 | FDR<0.1 | | fourth\_ventricle | 0.90 |
| fundus\_of\_striatum | 0.10 | 0.01 | 0.12 | 0.01 | -10.35 | -1.16 | 0.02 | 0.07 | FDR<0.1 | | fundus\_of\_striatum | 0.90 |
| globus\_pallidus | 0.84 | 0.09 | 0.92 | 0.07 | -8.12 | -1.00 | 0.06 | 0.09 | FDR<0.1 | | globus\_pallidus | 0.93 |
| hippocampus | 5.48 | 0.75 | 6.30 | 0.62 | -13.12 | -1.34 | 0.01 | 0.07 | FDR<0.1 | | hippocampus | 0.89 |
| hypothalamus | 5.77 | 0.51 | 6.30 | 0.44 | -8.38 | -1.19 | 0.02 | 0.07 | FDR<0.1 | | hypothalamus | 0.90 |
| internal\_capsule | 0.48 | 0.04 | 0.51 | 0.03 | -6.71 | -1.02 | 0.05 | 0.08 | FDR<0.1 | | internal\_capsule | 0.90 |
| lateral\_olfactory\_tract | 0.12 | 0.02 | 0.14 | 0.02 | -13.10 | -1.10 | 0.02 | 0.07 | FDR<0.1 | | lateral\_olfactory\_tract | 0.89 |
| lateral\_septum | 1.67 | 0.13 | 1.82 | 0.16 | -8.43 | -0.98 | 0.03 | 0.07 | FDR<0.1 | | lateral\_septum | 0.90 |
| lateral\_ventricle | 1.13 | 0.09 | 1.21 | 0.12 | -7.26 | -0.71 | 0.08 | 0.10 | | | lateral\_ventricle | 0.90 |
| mammillary\_bodies | 0.37 | 0.04 | 0.40 | 0.03 | -8.26 | -1.04 | 0.04 | 0.07 | FDR<0.1 | | mammillary\_bodies | 0.90 |
| mammilothalamic\_tract | 0.08 | 0.01 | 0.08 | 0.01 | -7.31 | -1.06 | 0.04 | 0.07 | FDR<0.1 | | mammilothalamic\_tract | 0.90 |
| medial\_lemniscus\_medial\_longitudinal\_fasciculus | 0.24 | 0.02 | 0.26 | 0.02 | -7.90 | -1.13 | 0.04 | 0.07 | FDR<0.1 | | medial\_lemniscus\_medial\_longitudinal\_fasciculus | 0.90 |
| medial\_septum | 0.26 | 0.02 | 0.28 | 0.02 | -8.18 | -1.04 | 0.02 | 0.07 | FDR<0.1 | | medial\_septum | 0.89 |
| medulla | 12.00 | 1.17 | 12.91 | 0.86 | -7.08 | -1.06 | 0.06 | 0.09 | FDR<0.1 | | medulla | 0.93 |
| midbrain | 7.42 | 0.62 | 7.90 | 0.61 | -6.05 | -0.78 | 0.10 | 0.12 | | | midbrain | 0.96 |
| nucleus\_accumbens | 1.20 | 0.11 | 1.29 | 0.09 | -7.08 | -0.96 | 0.07 | 0.09 | FDR<0.1 | | nucleus\_accumbens | 0.96 |
| olfactory\_bulbs | 6.33 | 0.87 | 7.29 | 0.99 | -13.14 | -0.97 | 0.03 | 0.07 | FDR<0.1 | | olfactory\_bulbs | 0.90 |
| olfactory\_tubercle | 1.21 | 0.15 | 1.41 | 0.16 | -14.10 | -1.26 | 0.01 | 0.07 | FDR<0.1 | | olfactory\_tubercle | 0.89 |
| optic\_tract | 0.27 | 0.03 | 0.30 | 0.02 | -10.07 | -1.39 | 0.01 | 0.07 | FDR<0.1 | | optic\_tract | 0.89 |
| periaqueductal\_grey | 1.76 | 0.16 | 1.86 | 0.14 | -5.54 | -0.72 | 0.15 | 0.16 | | | periaqueductal\_grey | 0.93 |
| pons | 7.02 | 0.60 | 7.35 | 0.46 | -4.52 | -0.73 | 0.18 | 0.18 | | | pons | 0.90 |
| pontine\_nucleus | 0.51 | 0.05 | 0.57 | 0.06 | -10.74 | -1.01 | 0.03 | 0.07 | FDR<0.1 | | pontine\_nucleus | 0.90 |
| posterior\_commissure | 0.01 | 0.00 | 0.01 | 0.00 | -8.33 | -1.58 | 0.02 | 0.07 | FDR<0.1 | | posterior\_commissure | 0.89 |
| pre\_para\_subiculum | 0.69 | 0.11 | 0.81 | 0.09 | -14.47 | -1.34 | 0.02 | 0.07 | FDR<0.1 | | pre\_para\_subiculum | 0.89 |
| stria\_medullaris | 0.17 | 0.02 | 0.19 | 0.02 | -10.16 | -1.14 | 0.02 | 0.07 | FDR<0.1 | | stria\_medullaris | 0.89 |
| stria\_terminalis | 0.17 | 0.01 | 0.19 | 0.01 | -7.78 | -1.13 | 0.02 | 0.07 | FDR<0.1 | | stria\_terminalis | 0.89 |
| striatum | 5.05 | 0.51 | 5.44 | 0.55 | -7.21 | -0.71 | 0.12 | 0.13 | | | striatum | 0.96 |
| subependymale\_zone\_rhinocele | 0.07 | 0.01 | 0.07 | 0.01 | -6.03 | -0.60 | 0.17 | 0.17 | | | subependymale\_zone\_rhinocele | 1.00 |
| superior\_olivary\_complex | 0.29 | 0.03 | 0.32 | 0.02 | -7.41 | -0.99 | 0.08 | 0.10 | FDR<0.1 | | superior\_olivary\_complex | 0.96 |
| thalamus | 5.73 | 0.56 | 6.37 | 0.58 | -9.95 | -1.08 | 0.02 | 0.07 | FDR<0.1 | | thalamus | 0.90 |
| third\_ventricle | 0.75 | 0.07 | 0.85 | 0.06 | -10.91 | -1.46 | 0.01 | 0.07 | FDR<0.1 | | third\_ventricle | 0.89 |
| medial\_preoptic\_area | 0.14 | 0.01 | 0.15 | 0.01 | -8.71 | -1.47 | 0.01 | 0.07 | FDR<0.1 | | medial\_preoptic\_area | 0.89 |
| paraventricular\_hypothalamic\_nucleus | 0.04 | 0.00 | 0.05 | 0.00 | -5.56 | -1.22 | 0.04 | 0.07 | FDR<0.1 | | paraventricular\_hypothalamic\_nucleus | 0.90 |
| brainvolumes | 129.34 | 14.54 | 143.63 | 12.57 | -9.95 | -1.14 | 0.03 | | | | brainvolumes | |

### Slide 4
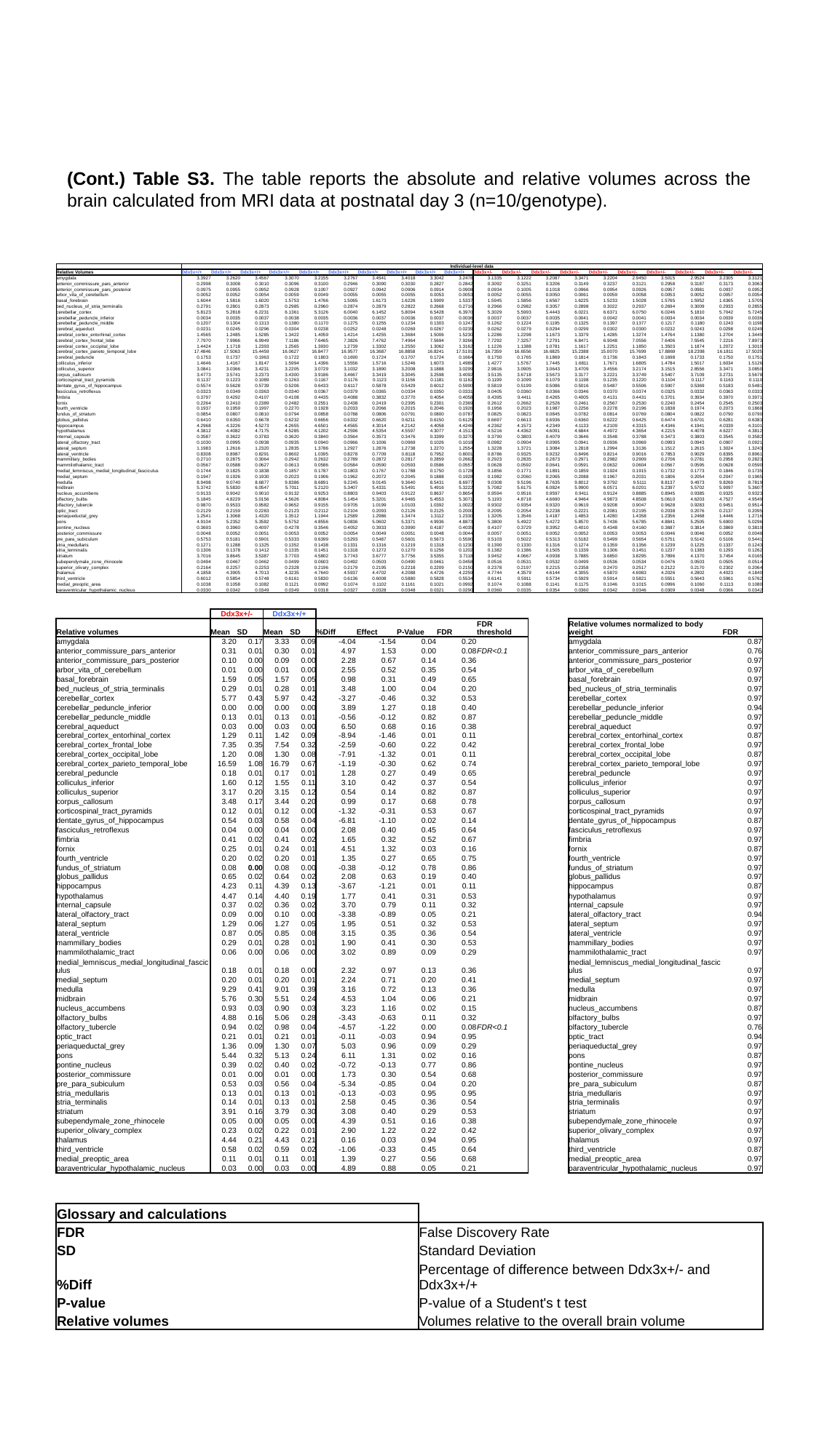

(Cont.) Table S3. The table reports the absolute and relative volumes across the brain calculated from MRI data at postnatal day 3 (n=10/genotype).
| | Individual-level data | | | | | | | | | | | | | | | | | | | |
| --- | --- | --- | --- | --- | --- | --- | --- | --- | --- | --- | --- | --- | --- | --- | --- | --- | --- | --- | --- | --- |
| Relative Volumes | Ddx3x+/+ | Ddx3x+/+ | Ddx3x+/+ | Ddx3x+/+ | Ddx3x+/+ | Ddx3x+/+ | Ddx3x+/+ | Ddx3x+/+ | Ddx3x+/+ | Ddx3x+/+ | Ddx3x+/- | Ddx3x+/- | Ddx3x+/- | Ddx3x+/- | Ddx3x+/- | Ddx3x+/- | Ddx3x+/- | Ddx3x+/- | Ddx3x+/- | Ddx3x+/- |
| amygdala | 3.3927 | 3.2620 | 3.4567 | 3.3070 | 3.2155 | 3.2767 | 3.4541 | 3.4018 | 3.3042 | 3.2478 | 3.1335 | 3.1222 | 3.2087 | 3.3471 | 3.2204 | 2.9450 | 3.5015 | 2.9524 | 3.2305 | 3.3121 |
| anterior\_commissure\_pars\_anterior | 0.2998 | 0.3008 | 0.3010 | 0.3096 | 0.3100 | 0.2946 | 0.3090 | 0.3030 | 0.2827 | 0.2842 | 0.3092 | 0.3251 | 0.3206 | 0.3149 | 0.3237 | 0.3121 | 0.2958 | 0.3187 | 0.3173 | 0.3063 |
| anterior\_commissure\_pars\_posterior | 0.0975 | 0.0955 | 0.0952 | 0.0928 | 0.1007 | 0.0927 | 0.0942 | 0.0906 | 0.0914 | 0.0908 | 0.0934 | 0.1005 | 0.1018 | 0.0966 | 0.0954 | 0.0926 | 0.0957 | 0.0981 | 0.0937 | 0.0952 |
| arbor\_vita\_of\_cerebellum | 0.0052 | 0.0052 | 0.0054 | 0.0059 | 0.0049 | 0.0055 | 0.0055 | 0.0055 | 0.0053 | 0.0053 | 0.0052 | 0.0055 | 0.0050 | 0.0061 | 0.0059 | 0.0058 | 0.0053 | 0.0052 | 0.0057 | 0.0054 |
| basal\_forebrain | 1.6044 | 1.5816 | 1.6020 | 1.5753 | 1.4766 | 1.5065 | 1.6173 | 1.6226 | 1.5909 | 1.5337 | 1.5945 | 1.5856 | 1.6567 | 1.6225 | 1.5233 | 1.5028 | 1.5765 | 1.5952 | 1.6365 | 1.5705 |
| bed\_nucleus\_of\_stria\_terminalis | 0.2791 | 0.2801 | 0.2873 | 0.2985 | 0.2960 | 0.2874 | 0.2879 | 0.2822 | 0.2668 | 0.2716 | 0.2966 | 0.2982 | 0.3057 | 0.2898 | 0.3022 | 0.2937 | 0.2694 | 0.3009 | 0.2933 | 0.2855 |
| cerebellar\_cortex | 5.8123 | 5.2818 | 6.2231 | 6.1361 | 5.3126 | 6.0040 | 6.1452 | 5.8094 | 6.5428 | 6.3970 | 5.3029 | 5.5993 | 5.4443 | 6.0221 | 6.6371 | 6.0750 | 6.0246 | 5.1810 | 5.7042 | 5.7245 |
| cerebellar\_peduncle\_inferior | 0.0034 | 0.0035 | 0.0037 | 0.0038 | 0.0035 | 0.0036 | 0.0037 | 0.0036 | 0.0037 | 0.0036 | 0.0037 | 0.0037 | 0.0035 | 0.0041 | 0.0042 | 0.0041 | 0.0034 | 0.0034 | 0.0039 | 0.0036 |
| cerebellar\_peduncle\_middle | 0.1207 | 0.1304 | 0.1313 | 0.1380 | 0.1170 | 0.1275 | 0.1255 | 0.1234 | 0.1303 | 0.1247 | 0.1262 | 0.1224 | 0.1195 | 0.1325 | 0.1397 | 0.1377 | 0.1217 | 0.1180 | 0.1243 | 0.1198 |
| cerebral\_aqueduct | 0.0231 | 0.0245 | 0.0296 | 0.0304 | 0.0238 | 0.0252 | 0.0248 | 0.0269 | 0.0267 | 0.0239 | 0.0262 | 0.0279 | 0.0294 | 0.0299 | 0.0302 | 0.0300 | 0.0232 | 0.0243 | 0.0298 | 0.0249 |
| cerebral\_cortex\_entorhinal\_cortex | 1.4565 | 1.2485 | 1.5285 | 1.3422 | 1.4059 | 1.4214 | 1.4255 | 1.3684 | 1.5006 | 1.5230 | 1.2286 | 1.2298 | 1.1673 | 1.3379 | 1.4285 | 1.3274 | 1.4764 | 1.1380 | 1.2704 | 1.3449 |
| cerebral\_cortex\_frontal\_lobe | 7.7970 | 7.9966 | 6.9949 | 7.1186 | 7.6465 | 7.3826 | 7.4762 | 7.4964 | 7.5694 | 7.9266 | 7.7292 | 7.3257 | 7.2791 | 6.8471 | 6.9048 | 7.0556 | 7.6406 | 7.5545 | 7.2216 | 7.8973 |
| cerebral\_cortex\_occipital\_lobe | 1.4424 | 1.1718 | 1.2393 | 1.2565 | 1.3930 | 1.2739 | 1.3302 | 1.2550 | 1.3062 | 1.3162 | 1.1226 | 1.1388 | 1.0781 | 1.1617 | 1.2251 | 1.1850 | 1.3503 | 1.1874 | 1.2072 | 1.3010 |
| cerebral\_cortex\_parieto\_temporal\_lobe | 17.4846 | 17.5063 | 15.4459 | 16.0627 | 16.8477 | 16.9577 | 16.3687 | 16.8858 | 16.8241 | 17.5191 | 16.7359 | 16.6556 | 16.6825 | 15.2388 | 15.0070 | 15.7699 | 17.8869 | 18.2398 | 16.1811 | 17.5025 |
| cerebral\_peduncle | 0.1753 | 0.1737 | 0.1963 | 0.1722 | 0.1803 | 0.1690 | 0.1724 | 0.1707 | 0.1724 | 0.1664 | 0.1750 | 0.1765 | 0.1869 | 0.1814 | 0.1736 | 0.1843 | 0.1698 | 0.1733 | 0.1750 | 0.1751 |
| colliculus\_inferior | 1.4646 | 1.4167 | 1.8147 | 1.5994 | 1.4396 | 1.5556 | 1.5716 | 1.5246 | 1.6073 | 1.4989 | 1.4277 | 1.5767 | 1.7445 | 1.6811 | 1.7671 | 1.6805 | 1.4784 | 1.5017 | 1.6034 | 1.5126 |
| colliculus\_superior | 3.0841 | 3.0366 | 3.4231 | 3.2205 | 3.0729 | 3.1032 | 3.1890 | 3.2008 | 3.1888 | 3.0299 | 2.9816 | 3.0905 | 3.0643 | 3.4709 | 3.4556 | 3.2174 | 3.1515 | 2.8556 | 3.3471 | 3.0850 |
| corpus\_callosum | 3.4773 | 3.5741 | 3.2373 | 3.4300 | 3.9186 | 3.4667 | 3.3419 | 3.3045 | 3.2598 | 3.4092 | 3.5135 | 3.6718 | 3.5673 | 3.3177 | 3.2221 | 3.3749 | 3.5407 | 3.7109 | 3.2731 | 3.5678 |
| corticospinal\_tract\_pyramids | 0.1137 | 0.1223 | 0.1089 | 0.1263 | 0.1167 | 0.1176 | 0.1123 | 0.1156 | 0.1181 | 0.1162 | 0.1199 | 0.1099 | 0.1079 | 0.1198 | 0.1235 | 0.1220 | 0.1104 | 0.1117 | 0.1163 | 0.1110 |
| dentate\_gyrus\_of\_hippocampus | 0.5574 | 0.5628 | 0.5739 | 0.5206 | 0.6433 | 0.6117 | 0.5878 | 0.5429 | 0.6012 | 0.5990 | 0.5819 | 0.5199 | 0.5086 | 0.5016 | 0.5487 | 0.5506 | 0.5907 | 0.5369 | 0.5183 | 0.5481 |
| fasciculus\_retroflexus | 0.0323 | 0.0349 | 0.0363 | 0.0340 | 0.0367 | 0.0379 | 0.0365 | 0.0334 | 0.0350 | 0.0331 | 0.0405 | 0.0360 | 0.0366 | 0.0346 | 0.0370 | 0.0374 | 0.0325 | 0.0332 | 0.0363 | 0.0333 |
| fimbria | 0.3797 | 0.4292 | 0.4107 | 0.4108 | 0.4435 | 0.4088 | 0.3832 | 0.3770 | 0.4054 | 0.4058 | 0.4395 | 0.4411 | 0.4265 | 0.4005 | 0.4131 | 0.4431 | 0.3701 | 0.3934 | 0.3970 | 0.3971 |
| fornix | 0.2264 | 0.2410 | 0.2389 | 0.2482 | 0.2551 | 0.2438 | 0.2419 | 0.2395 | 0.2301 | 0.2369 | 0.2612 | 0.2662 | 0.2526 | 0.2461 | 0.2567 | 0.2530 | 0.2240 | 0.2454 | 0.2545 | 0.2503 |
| fourth\_ventricle | 0.1937 | 0.1959 | 0.1997 | 0.2270 | 0.1928 | 0.2033 | 0.2066 | 0.2015 | 0.2046 | 0.1926 | 0.1956 | 0.2023 | 0.1987 | 0.2256 | 0.2278 | 0.2196 | 0.1838 | 0.1974 | 0.2073 | 0.1868 |
| fundus\_of\_striatum | 0.0854 | 0.0807 | 0.0810 | 0.0794 | 0.0858 | 0.0788 | 0.0806 | 0.0791 | 0.0800 | 0.0787 | 0.0825 | 0.0823 | 0.0845 | 0.0782 | 0.0814 | 0.0769 | 0.0804 | 0.0822 | 0.0790 | 0.0790 |
| globus\_pallidus | 0.6410 | 0.6350 | 0.6678 | 0.6232 | 0.6656 | 0.6332 | 0.6620 | 0.6211 | 0.6150 | 0.6129 | 0.6697 | 0.6613 | 0.6936 | 0.6360 | 0.6222 | 0.6425 | 0.6474 | 0.6701 | 0.6281 | 0.6383 |
| hippocampus | 4.2968 | 4.3226 | 4.5273 | 4.2655 | 4.6501 | 4.4565 | 4.3014 | 4.2142 | 4.4058 | 4.4246 | 4.2362 | 4.1573 | 4.2349 | 4.1133 | 4.2109 | 4.3315 | 4.4346 | 4.1941 | 4.0339 | 4.3101 |
| hypothalamus | 4.3812 | 4.4082 | 4.7175 | 4.5285 | 4.1202 | 4.2596 | 4.5354 | 4.5597 | 4.3077 | 4.1513 | 4.5216 | 4.4362 | 4.6091 | 4.6844 | 4.4972 | 4.3654 | 4.2215 | 4.4078 | 4.6227 | 4.3812 |
| internal\_capsule | 0.3587 | 0.3622 | 0.3783 | 0.3620 | 0.3840 | 0.3564 | 0.3573 | 0.3476 | 0.3399 | 0.3270 | 0.3790 | 0.3803 | 0.4079 | 0.3646 | 0.3548 | 0.3788 | 0.3473 | 0.3803 | 0.3545 | 0.3582 |
| lateral\_olfactory\_tract | 0.1030 | 0.0995 | 0.0938 | 0.0935 | 0.0940 | 0.0966 | 0.1006 | 0.0969 | 0.1026 | 0.1018 | 0.0982 | 0.0904 | 0.0995 | 0.0941 | 0.0936 | 0.0969 | 0.0993 | 0.0943 | 0.0907 | 0.0921 |
| lateral\_septum | 1.1983 | 1.2616 | 1.2320 | 1.2835 | 1.3786 | 1.2927 | 1.2876 | 1.2738 | 1.2270 | 1.2554 | 1.3228 | 1.3721 | 1.3084 | 1.2818 | 1.2994 | 1.3136 | 1.1512 | 1.2615 | 1.3024 | 1.3243 |
| lateral\_ventricle | 0.8308 | 0.8987 | 0.8291 | 0.8602 | 1.0395 | 0.8278 | 0.7709 | 0.8118 | 0.7952 | 0.8001 | 0.8786 | 0.9325 | 0.9232 | 0.8496 | 0.8214 | 0.9016 | 0.7853 | 0.9029 | 0.8395 | 0.8961 |
| mammillary\_bodies | 0.2710 | 0.2875 | 0.3064 | 0.2942 | 0.2632 | 0.2789 | 0.2872 | 0.2817 | 0.2859 | 0.2662 | 0.2923 | 0.2835 | 0.2873 | 0.2971 | 0.2982 | 0.2909 | 0.2706 | 0.2781 | 0.2958 | 0.2823 |
| mammilothalamic\_tract | 0.0567 | 0.0588 | 0.0627 | 0.0613 | 0.0586 | 0.0584 | 0.0590 | 0.0593 | 0.0586 | 0.0557 | 0.0628 | 0.0592 | 0.0641 | 0.0591 | 0.0632 | 0.0604 | 0.0567 | 0.0595 | 0.0628 | 0.0590 |
| medial\_lemniscus\_medial\_longitudinal\_fasciculus | 0.1744 | 0.1825 | 0.1838 | 0.1857 | 0.1787 | 0.1803 | 0.1767 | 0.1788 | 0.1750 | 0.1728 | 0.1856 | 0.1771 | 0.1891 | 0.1859 | 0.1924 | 0.1915 | 0.1732 | 0.1773 | 0.1846 | 0.1735 |
| medial\_septum | 0.1947 | 0.1926 | 0.1930 | 0.2023 | 0.1906 | 0.1962 | 0.2072 | 0.2045 | 0.1888 | 0.1928 | 0.1982 | 0.2060 | 0.2065 | 0.2088 | 0.1967 | 0.2031 | 0.1806 | 0.2054 | 0.2047 | 0.1965 |
| medulla | 8.9498 | 9.0740 | 8.6877 | 9.8386 | 8.6891 | 9.2245 | 9.0145 | 9.3640 | 8.5431 | 8.6977 | 9.0308 | 9.5196 | 8.7635 | 9.8012 | 9.3792 | 9.5111 | 8.8137 | 9.4973 | 9.8269 | 8.7819 |
| midbrain | 5.3742 | 5.5830 | 6.0547 | 5.7011 | 5.2120 | 5.3407 | 5.4331 | 5.5491 | 5.4916 | 5.3222 | 5.7082 | 5.6175 | 6.0824 | 5.9900 | 6.0571 | 6.0201 | 5.2397 | 5.5702 | 5.9097 | 5.3607 |
| nucleus\_accumbens | 0.9133 | 0.9042 | 0.9010 | 0.9132 | 0.9253 | 0.8803 | 0.9403 | 0.9122 | 0.8637 | 0.8654 | 0.9594 | 0.9516 | 0.9597 | 0.9411 | 0.9124 | 0.8885 | 0.8945 | 0.9385 | 0.9325 | 0.9323 |
| olfactory\_bulbs | 5.1845 | 4.8229 | 5.0156 | 4.5626 | 4.8084 | 5.1454 | 5.3201 | 4.9465 | 5.4553 | 5.3071 | 5.1193 | 4.8718 | 4.6690 | 4.9464 | 4.9873 | 4.8508 | 5.0610 | 4.6203 | 4.7527 | 4.9549 |
| olfactory\_tubercle | 0.9870 | 0.9533 | 0.9582 | 0.9652 | 0.9155 | 0.9705 | 1.0199 | 1.0103 | 1.0392 | 1.0022 | 0.9302 | 0.9354 | 0.9320 | 0.9619 | 0.9208 | 0.9047 | 0.9628 | 0.9283 | 0.9451 | 0.9514 |
| optic\_tract | 0.2129 | 0.2159 | 0.2283 | 0.2123 | 0.2112 | 0.2104 | 0.2093 | 0.2126 | 0.2125 | 0.2000 | 0.2095 | 0.2054 | 0.2238 | 0.2221 | 0.2081 | 0.2195 | 0.2038 | 0.2076 | 0.2137 | 0.2095 |
| periaqueductal\_grey | 1.2541 | 1.3068 | 1.4320 | 1.3512 | 1.1944 | 1.2589 | 1.2986 | 1.3474 | 1.3112 | 1.2330 | 1.3205 | 1.3546 | 1.4187 | 1.4853 | 1.4280 | 1.4358 | 1.2356 | 1.2468 | 1.4446 | 1.2716 |
| pons | 4.9104 | 5.2352 | 5.3582 | 5.5752 | 4.8556 | 5.0836 | 5.0602 | 5.3371 | 4.9936 | 4.8873 | 5.3800 | 5.4922 | 5.4272 | 5.8570 | 5.7436 | 5.6785 | 4.8841 | 5.2505 | 5.6900 | 5.0296 |
| pontine\_nucleus | 0.3693 | 0.3960 | 0.4097 | 0.4278 | 0.3546 | 0.4052 | 0.3933 | 0.3990 | 0.4187 | 0.4035 | 0.4107 | 0.3729 | 0.3952 | 0.4010 | 0.4348 | 0.4160 | 0.3687 | 0.3814 | 0.3869 | 0.3810 |
| posterior\_commissure | 0.0048 | 0.0052 | 0.0051 | 0.0053 | 0.0052 | 0.0054 | 0.0049 | 0.0051 | 0.0048 | 0.0044 | 0.0057 | 0.0051 | 0.0052 | 0.0052 | 0.0053 | 0.0053 | 0.0046 | 0.0046 | 0.0052 | 0.0048 |
| pre\_para\_subiculum | 0.5753 | 0.5181 | 0.5901 | 0.5333 | 0.6399 | 0.5293 | 0.5487 | 0.5601 | 0.5673 | 0.5590 | 0.5103 | 0.5022 | 0.5313 | 0.5182 | 0.5499 | 0.5654 | 0.5751 | 0.5142 | 0.5106 | 0.5441 |
| stria\_medullaris | 0.1271 | 0.1288 | 0.1325 | 0.1352 | 0.1438 | 0.1331 | 0.1316 | 0.1219 | 0.1318 | 0.1230 | 0.1390 | 0.1330 | 0.1316 | 0.1274 | 0.1359 | 0.1356 | 0.1239 | 0.1225 | 0.1337 | 0.1243 |
| stria\_terminalis | 0.1306 | 0.1378 | 0.1412 | 0.1335 | 0.1451 | 0.1318 | 0.1272 | 0.1270 | 0.1256 | 0.1202 | 0.1382 | 0.1386 | 0.1505 | 0.1339 | 0.1306 | 0.1451 | 0.1237 | 0.1383 | 0.1293 | 0.1262 |
| striatum | 3.7016 | 3.8645 | 3.5387 | 3.7703 | 4.5802 | 3.7743 | 3.6777 | 3.7756 | 3.5355 | 3.7118 | 3.9452 | 4.0667 | 4.0938 | 3.7885 | 3.6850 | 3.8295 | 3.7896 | 4.1370 | 3.7454 | 4.0165 |
| subependymale\_zone\_rhinocele | 0.0494 | 0.0467 | 0.0462 | 0.0499 | 0.0603 | 0.0492 | 0.0503 | 0.0490 | 0.0461 | 0.0458 | 0.0516 | 0.0531 | 0.0532 | 0.0499 | 0.0536 | 0.0534 | 0.0476 | 0.0503 | 0.0505 | 0.0514 |
| superior\_olivary\_complex | 0.2164 | 0.2257 | 0.2253 | 0.2328 | 0.2196 | 0.2179 | 0.2195 | 0.2218 | 0.2209 | 0.2150 | 0.2378 | 0.2197 | 0.2215 | 0.2358 | 0.2470 | 0.2517 | 0.2122 | 0.2170 | 0.2302 | 0.2064 |
| thalamus | 4.1858 | 4.3905 | 4.7013 | 4.3235 | 4.7640 | 4.5937 | 4.4702 | 4.2088 | 4.4726 | 4.2259 | 4.7744 | 4.3579 | 4.6144 | 4.3055 | 4.5870 | 4.6983 | 4.2026 | 4.2802 | 4.4023 | 4.1840 |
| third\_ventricle | 0.6012 | 0.5854 | 0.5748 | 0.6161 | 0.5830 | 0.6136 | 0.6008 | 0.5880 | 0.5828 | 0.5534 | 0.6141 | 0.5911 | 0.5734 | 0.5929 | 0.5914 | 0.5821 | 0.5551 | 0.5643 | 0.5961 | 0.5762 |
| medial\_preoptic\_area | 0.1038 | 0.1058 | 0.1082 | 0.1121 | 0.0992 | 0.1074 | 0.1102 | 0.1161 | 0.1021 | 0.0992 | 0.1074 | 0.1088 | 0.1141 | 0.1175 | 0.1046 | 0.1015 | 0.0996 | 0.1060 | 0.1113 | 0.1080 |
| paraventricular\_hypothalamic\_nucleus | 0.0330 | 0.0342 | 0.0349 | 0.0349 | 0.0318 | 0.0327 | 0.0328 | 0.0348 | 0.0321 | 0.0290 | 0.0360 | 0.0335 | 0.0354 | 0.0360 | 0.0342 | 0.0346 | 0.0309 | 0.0348 | 0.0366 | 0.0342 |
| | Ddx3x+/- | | Ddx3x+/+ | | | | | | | | | |
| --- | --- | --- | --- | --- | --- | --- | --- | --- | --- | --- | --- | --- |
| Relative volumes | Mean | SD | Mean | SD | %Diff | Effect | P-Value | FDR | FDR threshold | | Relative volumes normalized to body weight | FDR |
| amygdala | 3.20 | 0.17 | 3.33 | 0.09 | -4.04 | -1.54 | 0.04 | 0.20 | | | amygdala | 0.87 |
| anterior\_commissure\_pars\_anterior | 0.31 | 0.01 | 0.30 | 0.01 | 4.97 | 1.53 | 0.00 | 0.08 | FDR<0.1 | | anterior\_commissure\_pars\_anterior | 0.76 |
| anterior\_commissure\_pars\_posterior | 0.10 | 0.00 | 0.09 | 0.00 | 2.28 | 0.67 | 0.14 | 0.36 | | | anterior\_commissure\_pars\_posterior | 0.97 |
| arbor\_vita\_of\_cerebellum | 0.01 | 0.00 | 0.01 | 0.00 | 2.55 | 0.52 | 0.35 | 0.54 | | | arbor\_vita\_of\_cerebellum | 0.97 |
| basal\_forebrain | 1.59 | 0.05 | 1.57 | 0.05 | 0.98 | 0.31 | 0.49 | 0.65 | | | basal\_forebrain | 0.97 |
| bed\_nucleus\_of\_stria\_terminalis | 0.29 | 0.01 | 0.28 | 0.01 | 3.48 | 1.00 | 0.04 | 0.20 | | | bed\_nucleus\_of\_stria\_terminalis | 0.97 |
| cerebellar\_cortex | 5.77 | 0.43 | 5.97 | 0.42 | -3.27 | -0.46 | 0.32 | 0.53 | | | cerebellar\_cortex | 0.97 |
| cerebellar\_peduncle\_inferior | 0.00 | 0.00 | 0.00 | 0.00 | 3.89 | 1.27 | 0.18 | 0.40 | | | cerebellar\_peduncle\_inferior | 0.94 |
| cerebellar\_peduncle\_middle | 0.13 | 0.01 | 0.13 | 0.01 | -0.56 | -0.12 | 0.82 | 0.87 | | | cerebellar\_peduncle\_middle | 0.97 |
| cerebral\_aqueduct | 0.03 | 0.00 | 0.03 | 0.00 | 6.50 | 0.68 | 0.16 | 0.38 | | | cerebral\_aqueduct | 0.97 |
| cerebral\_cortex\_entorhinal\_cortex | 1.29 | 0.11 | 1.42 | 0.09 | -8.94 | -1.46 | 0.01 | 0.11 | | | cerebral\_cortex\_entorhinal\_cortex | 0.87 |
| cerebral\_cortex\_frontal\_lobe | 7.35 | 0.35 | 7.54 | 0.32 | -2.59 | -0.60 | 0.22 | 0.42 | | | cerebral\_cortex\_frontal\_lobe | 0.97 |
| cerebral\_cortex\_occipital\_lobe | 1.20 | 0.08 | 1.30 | 0.08 | -7.91 | -1.32 | 0.01 | 0.11 | | | cerebral\_cortex\_occipital\_lobe | 0.87 |
| cerebral\_cortex\_parieto\_temporal\_lobe | 16.59 | 1.08 | 16.79 | 0.67 | -1.19 | -0.30 | 0.62 | 0.74 | | | cerebral\_cortex\_parieto\_temporal\_lobe | 0.97 |
| cerebral\_peduncle | 0.18 | 0.01 | 0.17 | 0.01 | 1.28 | 0.27 | 0.49 | 0.65 | | | cerebral\_peduncle | 0.97 |
| colliculus\_inferior | 1.60 | 0.12 | 1.55 | 0.11 | 3.10 | 0.42 | 0.37 | 0.54 | | | colliculus\_inferior | 0.97 |
| colliculus\_superior | 3.17 | 0.20 | 3.15 | 0.12 | 0.54 | 0.14 | 0.82 | 0.87 | | | colliculus\_superior | 0.97 |
| corpus\_callosum | 3.48 | 0.17 | 3.44 | 0.20 | 0.99 | 0.17 | 0.68 | 0.78 | | | corpus\_callosum | 0.97 |
| corticospinal\_tract\_pyramids | 0.12 | 0.01 | 0.12 | 0.00 | -1.32 | -0.31 | 0.53 | 0.67 | | | corticospinal\_tract\_pyramids | 0.97 |
| dentate\_gyrus\_of\_hippocampus | 0.54 | 0.03 | 0.58 | 0.04 | -6.81 | -1.10 | 0.02 | 0.14 | | | dentate\_gyrus\_of\_hippocampus | 0.87 |
| fasciculus\_retroflexus | 0.04 | 0.00 | 0.04 | 0.00 | 2.08 | 0.40 | 0.45 | 0.64 | | | fasciculus\_retroflexus | 0.97 |
| fimbria | 0.41 | 0.02 | 0.41 | 0.02 | 1.65 | 0.32 | 0.52 | 0.67 | | | fimbria | 0.97 |
| fornix | 0.25 | 0.01 | 0.24 | 0.01 | 4.51 | 1.32 | 0.03 | 0.16 | | | fornix | 0.87 |
| fourth\_ventricle | 0.20 | 0.02 | 0.20 | 0.01 | 1.35 | 0.27 | 0.65 | 0.75 | | | fourth\_ventricle | 0.97 |
| fundus\_of\_striatum | 0.08 | 0.00 | 0.08 | 0.00 | -0.38 | -0.12 | 0.78 | 0.86 | | | fundus\_of\_striatum | 0.97 |
| globus\_pallidus | 0.65 | 0.02 | 0.64 | 0.02 | 2.08 | 0.63 | 0.19 | 0.40 | | | globus\_pallidus | 0.97 |
| hippocampus | 4.23 | 0.11 | 4.39 | 0.13 | -3.67 | -1.21 | 0.01 | 0.11 | | | hippocampus | 0.87 |
| hypothalamus | 4.47 | 0.14 | 4.40 | 0.19 | 1.77 | 0.41 | 0.31 | 0.53 | | | hypothalamus | 0.97 |
| internal\_capsule | 0.37 | 0.02 | 0.36 | 0.02 | 3.70 | 0.79 | 0.11 | 0.32 | | | internal\_capsule | 0.97 |
| lateral\_olfactory\_tract | 0.09 | 0.00 | 0.10 | 0.00 | -3.38 | -0.89 | 0.05 | 0.21 | | | lateral\_olfactory\_tract | 0.94 |
| lateral\_septum | 1.29 | 0.06 | 1.27 | 0.05 | 1.95 | 0.51 | 0.32 | 0.53 | | | lateral\_septum | 0.97 |
| lateral\_ventricle | 0.87 | 0.05 | 0.85 | 0.08 | 3.15 | 0.35 | 0.36 | 0.54 | | | lateral\_ventricle | 0.97 |
| mammillary\_bodies | 0.29 | 0.01 | 0.28 | 0.01 | 1.90 | 0.41 | 0.30 | 0.53 | | | mammillary\_bodies | 0.97 |
| mammilothalamic\_tract | 0.06 | 0.00 | 0.06 | 0.00 | 3.02 | 0.89 | 0.09 | 0.29 | | | mammilothalamic\_tract | 0.97 |
| medial\_lemniscus\_medial\_longitudinal\_fasciculus | 0.18 | 0.01 | 0.18 | 0.00 | 2.32 | 0.97 | 0.13 | 0.36 | | | medial\_lemniscus\_medial\_longitudinal\_fasciculus | 0.97 |
| medial\_septum | 0.20 | 0.01 | 0.20 | 0.01 | 2.24 | 0.71 | 0.20 | 0.41 | | | medial\_septum | 0.97 |
| medulla | 9.29 | 0.41 | 9.01 | 0.39 | 3.16 | 0.72 | 0.13 | 0.36 | | | medulla | 0.97 |
| midbrain | 5.76 | 0.30 | 5.51 | 0.24 | 4.53 | 1.04 | 0.06 | 0.21 | | | midbrain | 0.97 |
| nucleus\_accumbens | 0.93 | 0.03 | 0.90 | 0.03 | 3.23 | 1.16 | 0.02 | 0.15 | | | nucleus\_accumbens | 0.87 |
| olfactory\_bulbs | 4.88 | 0.16 | 5.06 | 0.28 | -3.43 | -0.63 | 0.11 | 0.32 | | | olfactory\_bulbs | 0.97 |
| olfactory\_tubercle | 0.94 | 0.02 | 0.98 | 0.04 | -4.57 | -1.22 | 0.00 | 0.08 | FDR<0.1 | | olfactory\_tubercle | 0.76 |
| optic\_tract | 0.21 | 0.01 | 0.21 | 0.01 | -0.11 | -0.03 | 0.94 | 0.95 | | | optic\_tract | 0.94 |
| periaqueductal\_grey | 1.36 | 0.09 | 1.30 | 0.07 | 5.03 | 0.96 | 0.09 | 0.29 | | | periaqueductal\_grey | 0.97 |
| pons | 5.44 | 0.32 | 5.13 | 0.24 | 6.11 | 1.31 | 0.02 | 0.16 | | | pons | 0.87 |
| pontine\_nucleus | 0.39 | 0.02 | 0.40 | 0.02 | -0.72 | -0.13 | 0.77 | 0.86 | | | pontine\_nucleus | 0.97 |
| posterior\_commissure | 0.01 | 0.00 | 0.01 | 0.00 | 1.73 | 0.30 | 0.54 | 0.68 | | | posterior\_commissure | 0.97 |
| pre\_para\_subiculum | 0.53 | 0.03 | 0.56 | 0.04 | -5.34 | -0.85 | 0.04 | 0.20 | | | pre\_para\_subiculum | 0.87 |
| stria\_medullaris | 0.13 | 0.01 | 0.13 | 0.01 | -0.13 | -0.03 | 0.95 | 0.95 | | | stria\_medullaris | 0.97 |
| stria\_terminalis | 0.14 | 0.01 | 0.13 | 0.01 | 2.58 | 0.45 | 0.36 | 0.54 | | | stria\_terminalis | 0.97 |
| striatum | 3.91 | 0.16 | 3.79 | 0.30 | 3.08 | 0.40 | 0.29 | 0.53 | | | striatum | 0.97 |
| subependymale\_zone\_rhinocele | 0.05 | 0.00 | 0.05 | 0.00 | 4.39 | 0.51 | 0.16 | 0.38 | | | subependymale\_zone\_rhinocele | 0.97 |
| superior\_olivary\_complex | 0.23 | 0.02 | 0.22 | 0.01 | 2.90 | 1.22 | 0.22 | 0.42 | | | superior\_olivary\_complex | 0.97 |
| thalamus | 4.44 | 0.21 | 4.43 | 0.21 | 0.16 | 0.03 | 0.94 | 0.95 | | | thalamus | 0.97 |
| third\_ventricle | 0.58 | 0.02 | 0.59 | 0.02 | -1.06 | -0.33 | 0.45 | 0.64 | | | third\_ventricle | 0.87 |
| medial\_preoptic\_area | 0.11 | 0.01 | 0.11 | 0.01 | 1.39 | 0.27 | 0.56 | 0.68 | | | medial\_preoptic\_area | 0.97 |
| paraventricular\_hypothalamic\_nucleus | 0.03 | 0.00 | 0.03 | 0.00 | 4.89 | 0.88 | 0.05 | 0.21 | | | paraventricular\_hypothalamic\_nucleus | 0.97 |
| Glossary and calculations | |
| --- | --- |
| FDR | False Discovery Rate |
| SD | Standard Deviation |
| %Diff | Percentage of difference between Ddx3x+/- and Ddx3x+/+ |
| P-value | P-value of a Student's t test |
| Relative volumes | Volumes relative to the overall brain volume |
