## Supplemental Figures for "Developmental and behavioral phenotypes in a new mouse model of DDX3X syndrome"

### Supplementary Figure 1

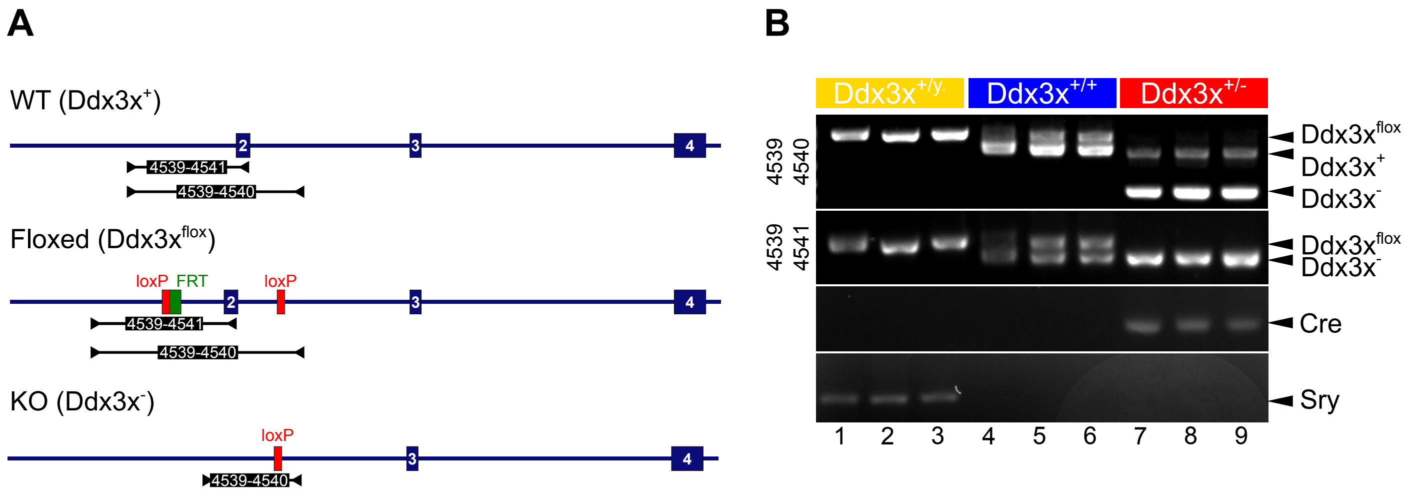

#### Supplementary Figure 2

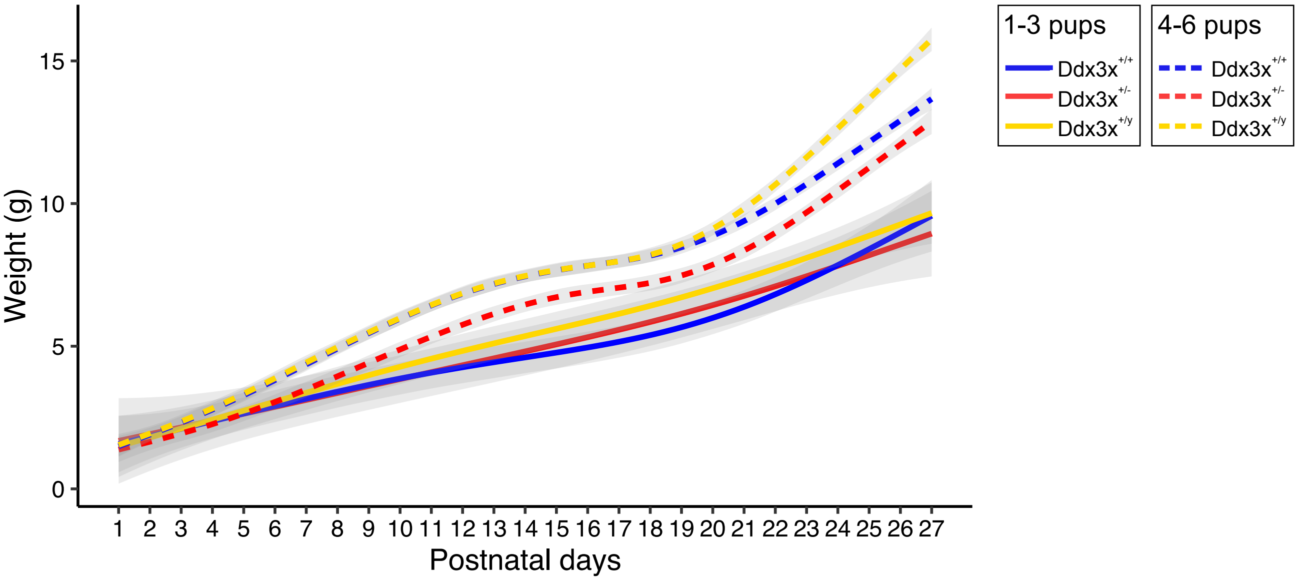

### Supplementary Figure 3

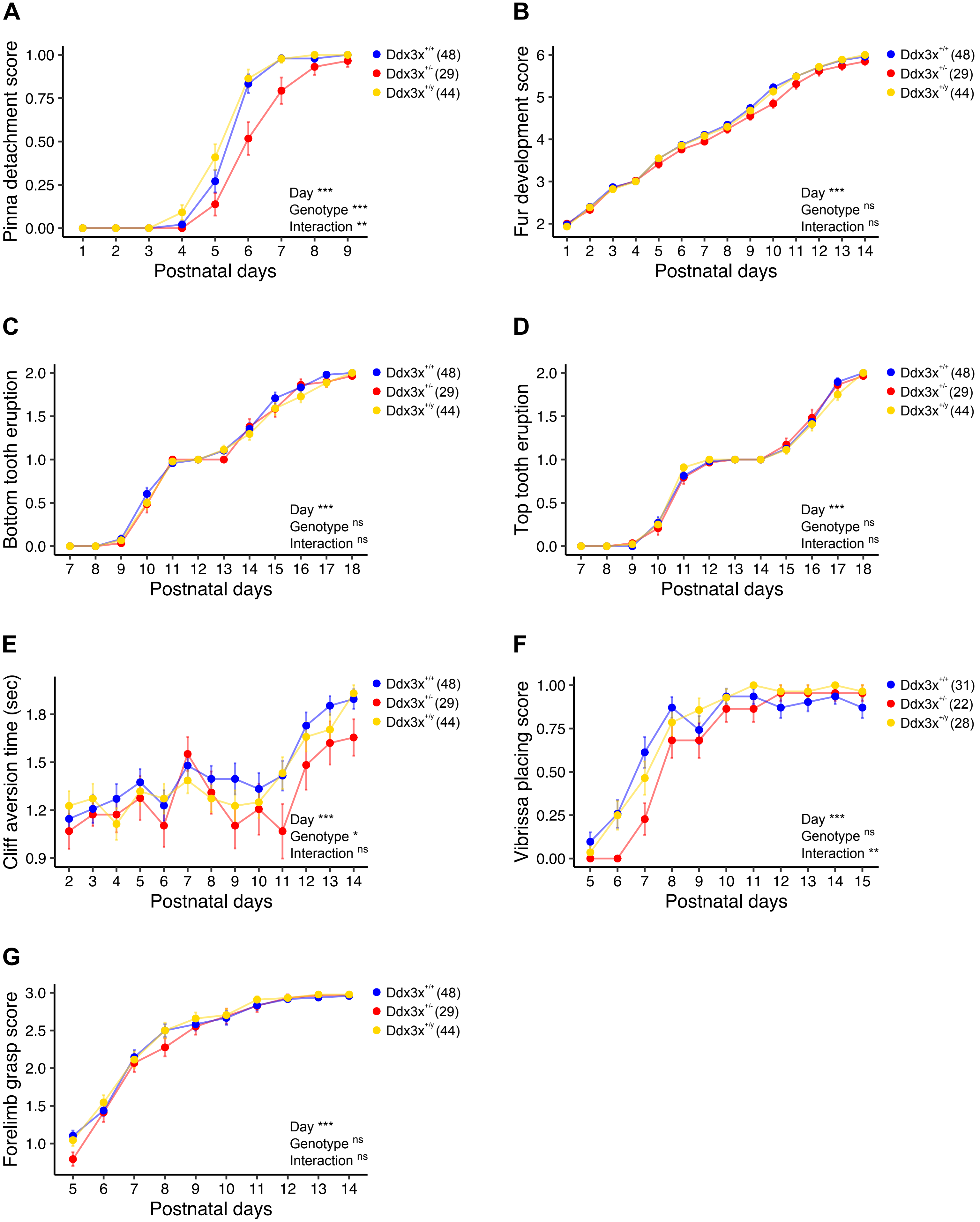

### Supplementary Figure 4

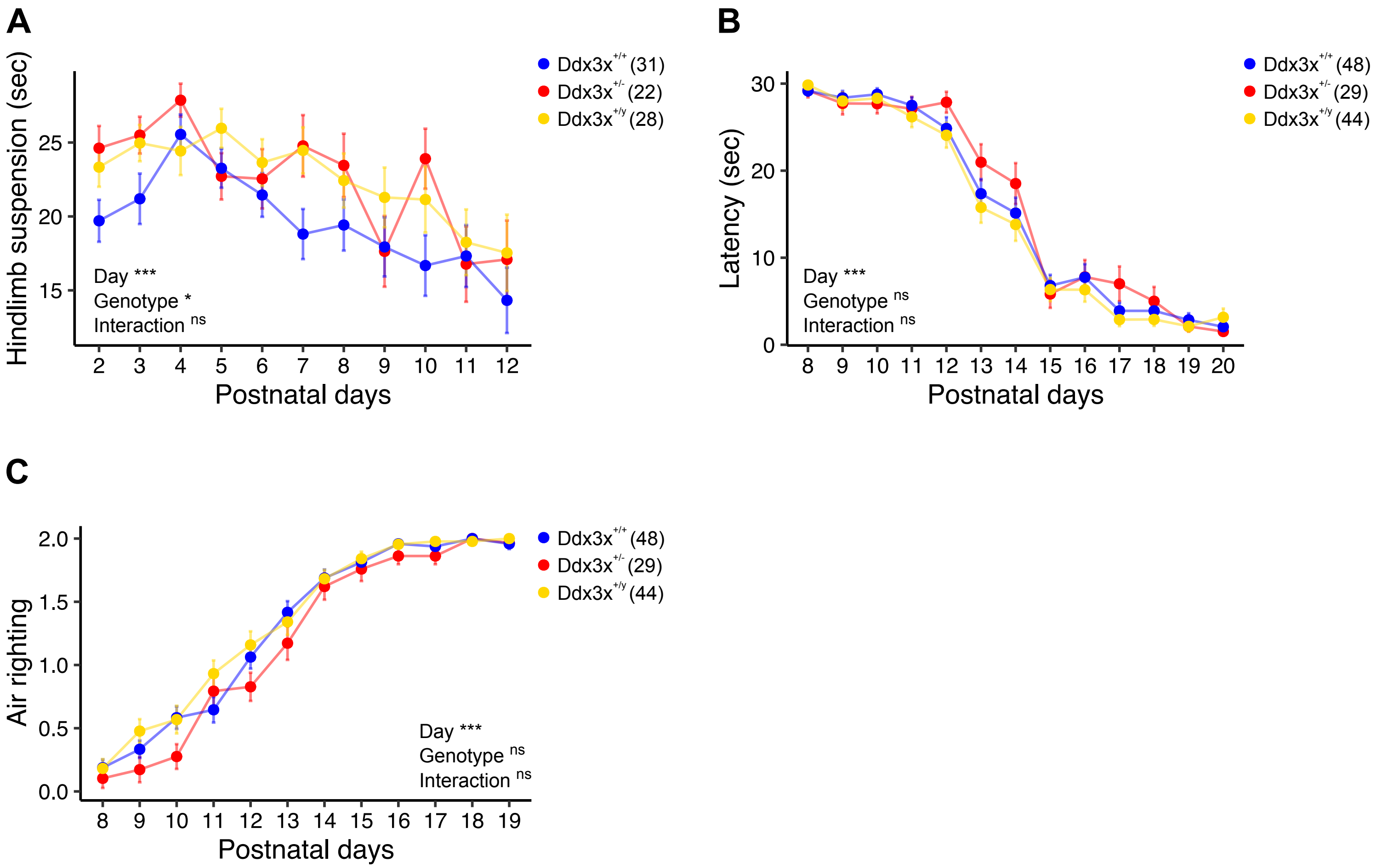

#### Supplementary Figure 5

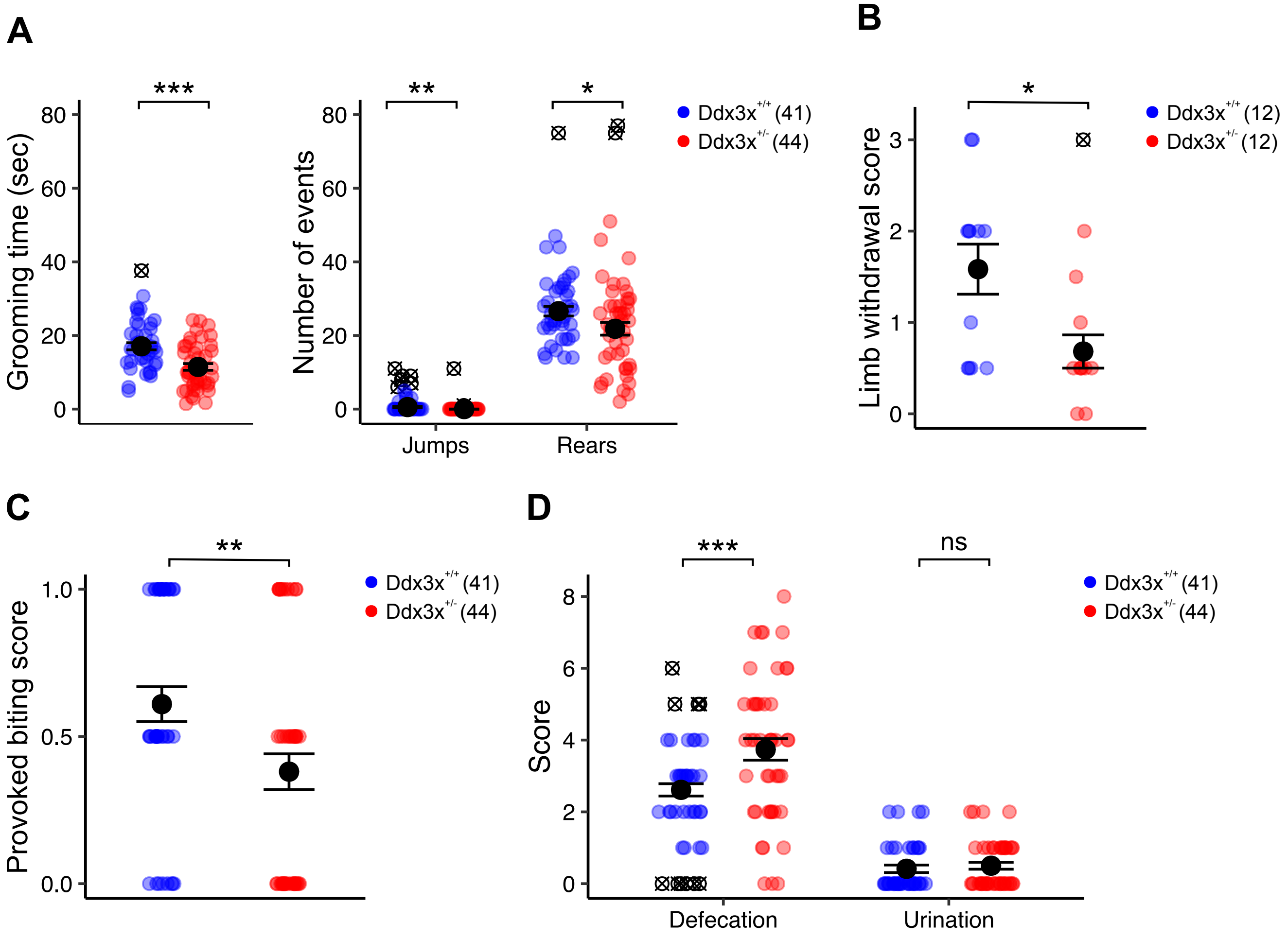

#### Supplementary Figure 6

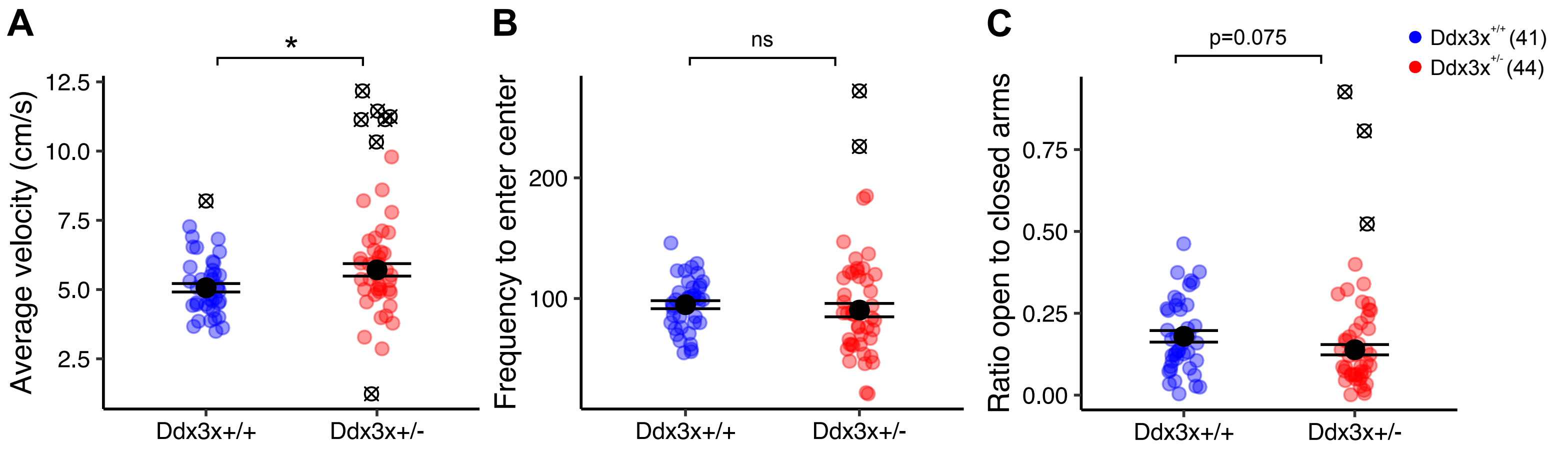


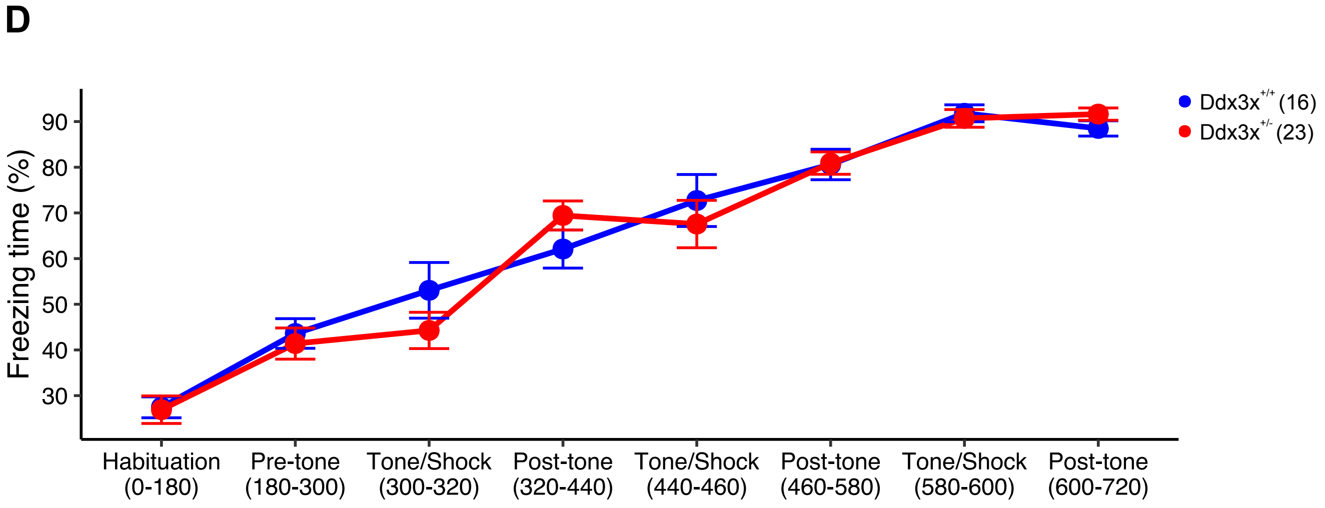

### Supplementary Figure 7

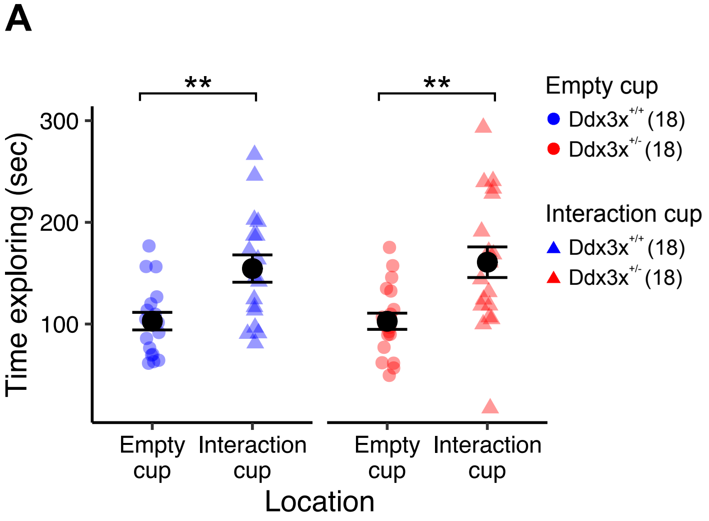


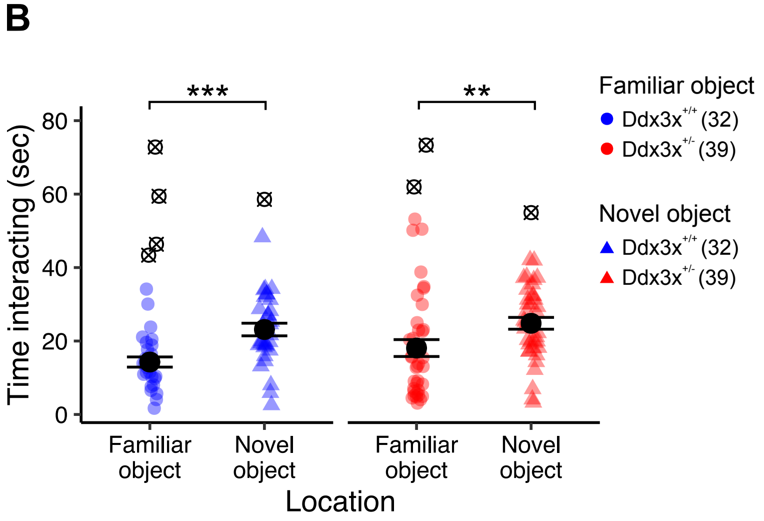

#### Supplementary Figure 8

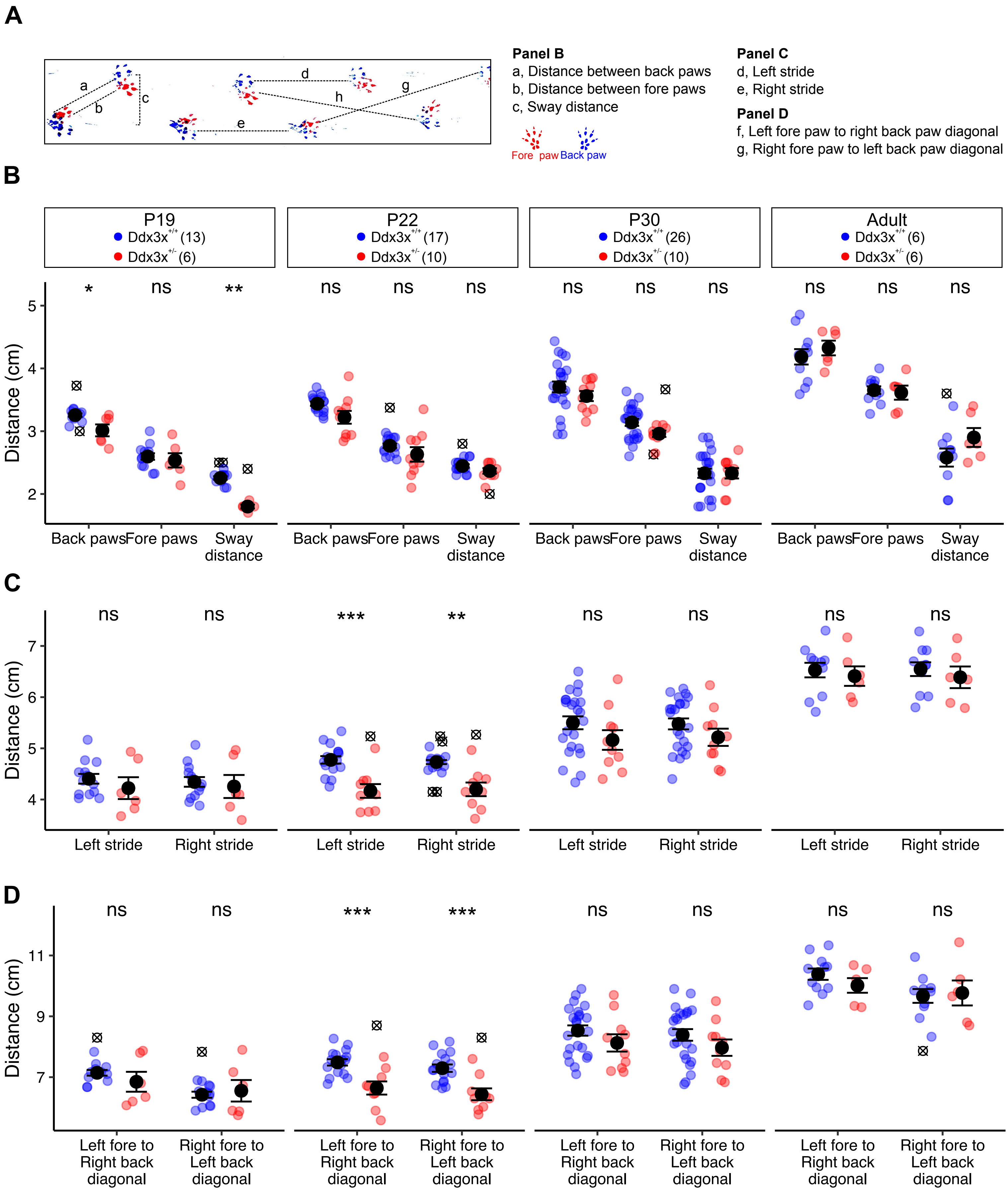

#### Supplementary Figure 9

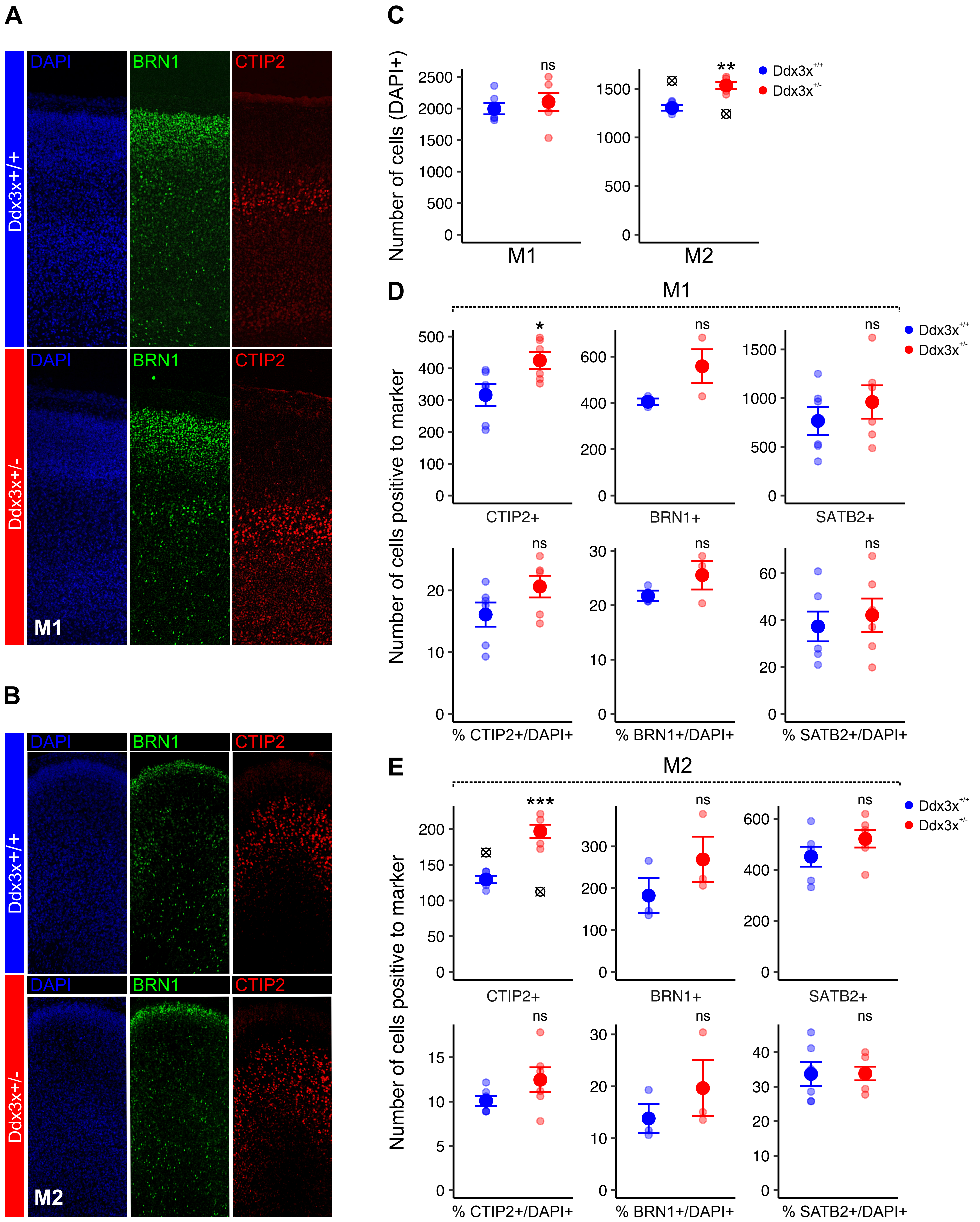

#### Supplementary Figure 10

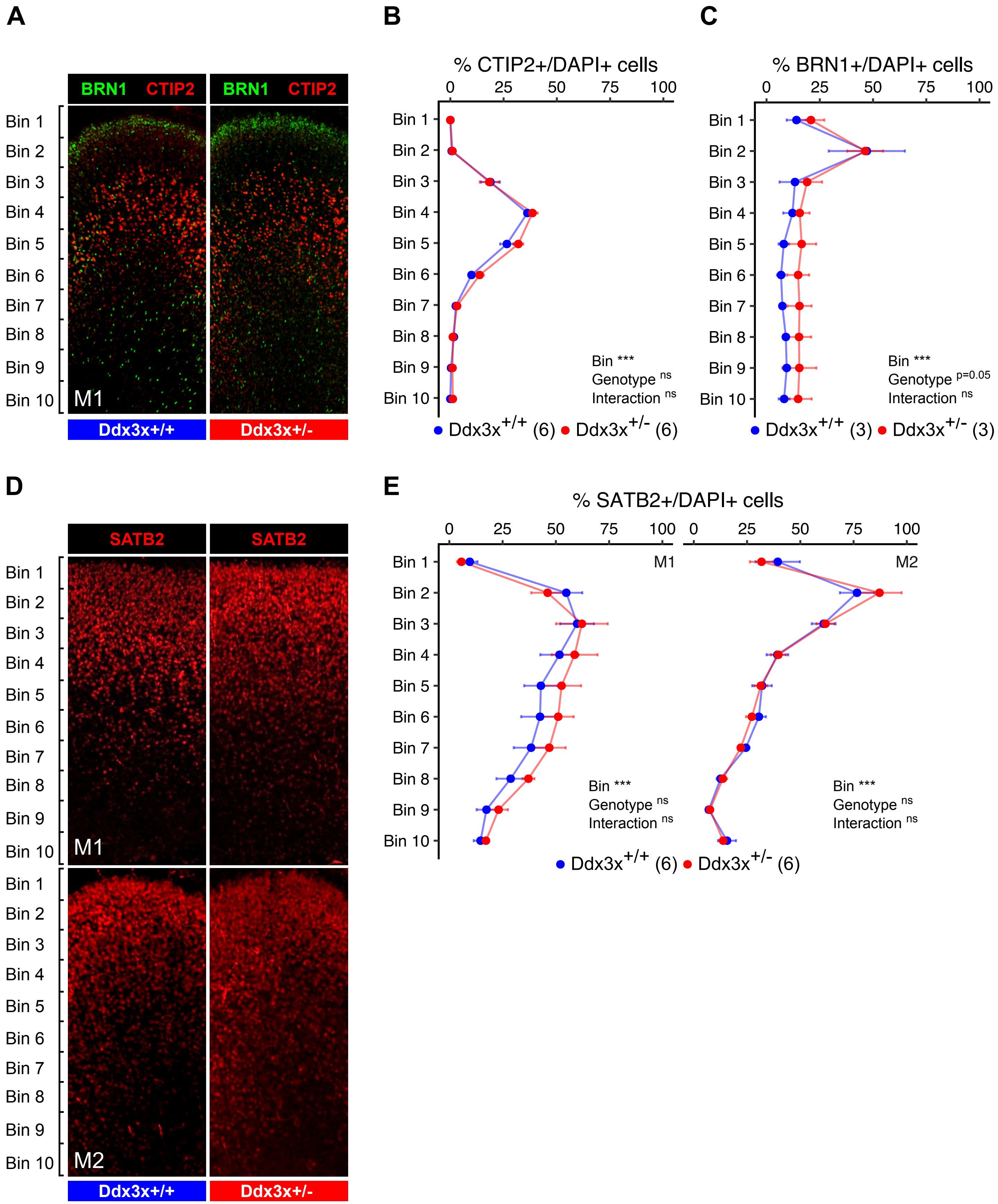
