## Supplemental Notes for "Developmental and behavioral phenotypes in a new mouse model of DDX3X syndrome"

### Supplemental Figure Legends

**Figure S1. Design and genotyping of the *Ddx3x*$^{flox}$ and *Ddx3x*$^{+/-}$ lines. A) Schematic representation of the murine *Ddx3x* wild-type (WT, *Ddx3x*$^{+}$), floxed (*Ddx3x*$^{flox}$) and knockout (KO, *Ddx3x*$^{-}$) alleles.** The scheme shows the arrangement of exons 2, 3, and 4 in the WT allele (upper panel), the location of the loxP sites (red) surrounding exon 2 of the flox allele (middle panel), and the excision of exon 2 in the KO allele (lower panel). The FLP recombinase target sequence (FRT, shown in green) is residual from an FLP-mediated removal of a neomycin cassette. The two primer pairs used for genotyping (see **Methods**) are also shown. **B) Genotyping of the *Ddx3x*$^{+}$, *Ddx3x*$^{flox}$ and *Ddx3x*$^{-}$ alleles.** The image shows an agarose gel electrophoresis results of a PCR on genomic DNA isolated from the offspring of *Ddx3x*$^{flox/+}$ females crossed with Sox2-Cre/+ males. The samples tested included 3 *Ddx3x*$^{flox/y}$;+/+ males (indicated as *Ddx3x*$^{+/y}$; yellow; lanes 1-3), 3 *Ddx3x*$^{flox/+}$;+/+ females (indicated as *Ddx3x*$^{+/+}$; blue; lanes 4-6) and 3 *Ddx3x*$^{flox/+}$;Sox2-Cre/+ females (indicated as *Ddx3x*$^{+/-}$; red; lanes 7-9). The PCR amplified the *Ddx3x*$^{+}$, *Ddx3x*$^{flox}$ and *Ddx3x*$^{-}$ alleles (using the primers pairs shown in panel A), the *Cre* transgene, and the male-specific locus *Sry*.

**Figure S2. Relationship between litter size and physical growth.** The plot shows the body weight from postnatal day (P) 1 to 27 across the three genotypes as in **Fig. 2A**, but broken down by litter size. Irrespective to genotype, pups in larger litter (4 to 6 pups, dashed lines) have better growth curves than pups in smaller litters (1-3 pups, solid lines) (smoothed conditional mean ± 0.95 confidence interval displayed in grey). All data collected and scored blind to genotype. In all panels, *Ddx3x*$^{+/+}$ (blue), *Ddx3x*$^{+/-}$ (red), and *Ddx3x*$^{+/y}$ (yellow).

**Figure S3. Additional physical and sensory milestones in *Ddx3x*$^{+/-}$ mice. A) *Ddx3x*$^{+/-}$ pups have a delay in ear development.** The plot shows the pinna detachment score over time across the three genotypes (*n* shown in legend; mean ± SEM; repeated measure ANOVA between *Ddx3x*$^{+/+}$ and *Ddx3x*$^{+/-}$ genotypes). **B) *Ddx3x*$^{+/-}$ pups have no changes in fur and skin development.** The plot shows the fur/skin development score over time across the three genotypes (*n* shown in legend; mean ± SEM; repeated measure ANOVA between *Ddx3x*$^{+/+}$ and *Ddx3x*$^{+/-}$ genotypes). **C) *Ddx3x*$^{+/-}$ pups have no changes in bottom tooth development.** The plot shows the score for the eruption of the bottom incisors over time across the three genotypes (*n* shown in legend; mean ± SEM; repeated measure ANOVA between *Ddx3x*$^{+/+}$ and *Ddx3x*$^{+/-}$ genotypes). **D) *Ddx3x*$^{+/-}$ pups have no changes in top tooth development.** The plot shows the score for the eruption of the top incisors over time across the three genotypes (*n* shown in legend; mean ± SEM; repeated measure ANOVA between *Ddx3x*$^{+/+}$ and *Ddx3x*$^{+/-}$ genotypes). **E) *Ddx3x*$^{+/-}$ pups display a delay in developing the cliff aversion reflex.** The plot shows the response to the test over time across the three genotypes (*n* shown in legend; mean ± SEM; repeated measure ANOVA between *Ddx3x*$^{+/+}$ and *Ddx3x*$^{+/-}$ genotypes). **F) *Ddx3x*$^{+/-}$ pups display a delay in the vibrissae placing reflex.** The plot shows the response to the test over time across the three genotypes (*n* shown in legend; mean ± SEM; repeated measure ANOVA between *Ddx3x*$^{+/+}$ and *Ddx3x*$^{+/-}$ genotypes). **G) *Ddx3x*$^{+/-}$ pups have no impairments in forelimb grasp.** The plot shows the forelimb grasp score over time across the three genotypes (*n* shown in legend; mean ± SEM; repeated measure ANOVA between *Ddx3x*$^{+/+}$ and *Ddx3x*$^{+/-}$ genotypes).

All data collected and scored blind to genotype. In all panels, *Ddx3x*$^{+/+}$ (blue), *Ddx3x*$^{+/-}$ (red), and *Ddx3x*$^{+/y}$ (yellow). *p<0.05, **p<0.01, ***p<0.001; ns, non significant.

**Figure S4. Additional motor milestones in *Ddx3x*$^{+/-}$ mice. A) *Ddx3x*$^{+/-}$ pups have a mild delay in developing hind limb suspension skills.** The plot shows the hind limb suspension score over time across the three genotypes (*n* shown in legend; mean ± SEM; repeated measure ANOVA between *Ddx3x*$^{+/+}$ and *Ddx3x*$^{+/-}$ genotypes). **B) *Ddx3x*$^{+/-}$ pups have no changes in overall locomotor activity in an open field crossing task.** The plot shows the latency to leave (sec) the center of an open field, and at this developmental stage is a measure of locomotor activity (*n* shown in legend; mean ± SEM; repeated measure ANOVA between *Ddx3x*$^{+/+}$ and *Ddx3x*$^{+/-}$ genotypes). **C) *Ddx3x*$^{+/-}$ pups have no impairments in air righting abilities.** The plot shows the time to right the body axis (sec) over time across the three genotypes (*n* shown in legend; mean ± SEM; repeated measure ANOVA between *Ddx3x*$^{+/+}$ and *Ddx3x*$^{+/-}$ genotypes).

All data collected and scored blind to genotype. In all panels, *Ddx3x*$^{+/+}$ (blue), *Ddx3x*$^{+/-}$ (red), and *Ddx3x*$^{+/y}$ (yellow). *p<0.05, ***p<0.001; ns, non significant.

**Figure S5. Changes in some measures of spontaneous activity in 4-month old animals. A) *Ddx3x*$^{+/-}$ mice have reduced grooming, number of jumps and rears.** The plot shows the grooming time (sec), number of jumps, or number of rears across the two genotypes (*n* shown in legend; mean ± SEM; Student's t test after outliers removal and Shapiro-Wilk test for normality for grooming time and number of rears; Wilcoxon signed-rank test after outliers removal and Shapiro-Wilk test for normality for number of jumps; ⊗ indicate outliers). **B) *Ddx3x*$^{+/-}$ mice have reduced nociception.** The plot shows the limb withdrawal across the two genotypes (*n* shown in legend; mean ± SEM; Student's t test after outliers removal and Shapiro-Wilk test for normality; ⊗ indicate outliers). **C) *Ddx3x*$^{+/-}$ mice have reduced provoked biting reflex.** The plot shows the score for the provoked biting reflex for the two genotypes (*n* shown in legend; mean ± SEM; Wilcoxon signed-rank test after outliers removal and Shapiro-Wilk test for normality). **D) *Ddx3x*$^{+/-}$ mice display increased defecation but typical urination.** The plot shows the score for the defecation and urination scores for the two genotypes (*n* shown in legend; mean ± SEM; Wilcoxon signed-rank test after outliers removal and Shapiro-Wilk test for normality; ⊗ indicate outliers).

All data collected and scored blind to genotype. In all panels, *Ddx3x*$^{+/+}$ (blue), and *Ddx3x*$^{+/-}$ (red). *p<0.05, **p<0.01, ***p<0.001; ns, non significant.

**Figure S6. Additional measures in open field test, elevated plus maze test, and fear conditioning training. A) *Ddx3x*$^{+/-}$ mice show increased velocity in open field test.** The plot shows the velocity of *Ddx3x*$^{+/+}$ (blue) and *Ddx3x*$^{+/-}$ (red) mice in the 30-minute open field test, in line with findings of increase locomotor activity (**Fig. 4A-B**) (*n* shown in legend; mean ± SEM; Student's t test after outliers removal and Shapiro-Wilk test for normality; ⊗ indicate outliers; *p<0.05). **B) *Ddx3x*$^{+/-}$ mice enter the center zone of the open field as often as their control littermates.** The plot shows the frequency to enter the center zone for the two genotypes (*n* shown in legend; mean ± SEM; ⊗ indicate outliers). **C) *Ddx3x*$^{+/-}$ mice have no deficits in elevated plus maze.** The plot shows the ratio of the time spent in the open arms vs the time spent in the closed arm in the elevated plus maze test (*n* shown in legend; Wilcoxon signed-rank test after outliers removal and Shapiro-Wilk test for normality; ⊗ indicate outliers). **D) *Ddx3x*$^{+/-}$ mice have no alterations in fear conditioning training.** The plot shows the percentage of time spent freezing during training (*n* shown in legend).

All data collected and scored blind to genotype. In all panels, *Ddx3x*$^{+/+}$ (blue), and *Ddx3x*$^{+/-}$ (red). *p<0.05; ns, non significant.

**Figure S7. *Ddx3x*$^{+/-}$ mice show no changes in sociability and novel object recognition. A) *Ddx3x*$^{+/-}$ mice spend more time with a mouse than an object, as their control littermates.** The plot shows the time *Ddx3x*$^{+/+}$ and *Ddx3x*$^{+/-}$ mice spent sniffing an empty cup (triangles) or a cup with another age-matched female mouse (circles) during a 10-min testing session (see **Methods**) (*n* shown in legend; mean ± SEM; Student's t test after outliers removal and Shapiro-Wilk test for normality; ⊗ indicate outliers). **B) *Ddx3x*$^{+/-}$ mice spend more time with a novel object than a familiar one, as their control littermates.** The plot shows the time *Ddx3x*$^{+/+}$ (blue) and *Ddx3x*$^{+/-}$ (red) mice spent interacting a novel object (triangles) or a familiar object (circles) during a 5 min testing session (see **Methods**) (*n* shown in legend; mean ± SEM; Student's t test after outliers removal and Shapiro-Wilk test for normality in *Ddx3x*$^{+/+}$ mice; Wilcoxon signed-rank test after outliers removal and Shapiro-Wilk test for normality in *Ddx3x*$^{+/-}$ mice; ⊗ indicate outliers).

All data collected and scored blind to genotype. In all panels, *Ddx3x*$^{+/+}$ (blue), and *Ddx3x*$^{+/-}$ (red). **p<0.01, ***p<0.001; ns, non significant.

**Figure S8. *Ddx3x*$^{+/-}$ mice have transient gait anomalies.** Gait was analyzed based on footprint patterns of mice walking in a straight line. The back paws were coated in blue paint and the front paws were coated in red paint and the mice were released onto a strip of paper placed in a runway. The different parameters measured to assess gait are shown. Plots show the length of each of the 7 parameters (panel A) measured in *Ddx3x*$^{+/+}$ and *Ddx3x*$^{+/-}$ mice at P19, P22, P30 and adulthood (*n* shown in legend; mean ± SEM; Student's t test or Wilcoxon signed-rank test after outliers removal and Shapiro-Wilk test for normality; ⊗ indicate outliers).

All data collected and scored blind to genotype. In all panels, *Ddx3x*$^{+/+}$ (blue) and *Ddx3x*$^{+/-}$ (red). *p<0.05, **p<0.01, ***p<0.001; ns, non significant.

**Figure S9. *Ddx3x*$^{+/-}$ mice have altered cortical lamination. A) Subcerebral (ScPN) and intra-telencephalic (IT) projection neurons in the developing primary motor cortex (M1).** Representative confocal images of coronal sections of M1 from *Ddx3x*$^{+/+}$ and *Ddx3x*$^{+/-}$ mice at P3, immunostained for DAPI (blue), CTIP2 (red), a marker of ScPN and BRN1 (green), a marker of IT. These images are the single-channel images of those shown in **Fig. 7A**. **B) ScPN and IT projection neurons in the developing secondary motor cortex (M2).** Representative confocal images of coronal sections of M2 from *Ddx3x*$^{+/+}$ and *Ddx3x*$^{+/-}$ mice at P3 immunostained for DAPI (blue), CTIP2 (red), and BRN1 (green). These images are the single-channel images of those shown in **Fig. S10A**. **C) *Ddx3x*$^{+/-}$ mice have more cells in M2.** The plots show the total number of cells (DAPI+) in M1 and M2 (*n*=6; 6-8 sections/mouse, with outliers across sections removed; mean ± SEM; Student's t test after outliers removal and Shapiro-Wilk test for normality; ⊗ indicate outliers). **D) *Ddx3x*$^{+/-}$ mice**

**have more CTIP2+ ScPN in M1. D)** The plots show the total number of cells positive for each marker in M1 (upper row) or the number of marker-positive cells as a percentage to the total number of cells (DAPI+) (*n*=6 for CTIP2 and SATB2; *n*=3 for BRN1; 6-8 sections/mouse, with outliers across sections removed; mean ± SEM; Student's t test after outliers removal and Shapiro-Wilk test for normality; ⊗ indicate outliers). **E)** ***Ddx3x***$^{+/-}$ **mice have more CTIP2+ ScPN in M2.** The plots show the total number of cells positive for each marker in M1 (upper row) or the number of marker-positive cells as a percentage to the total number of cells (DAPI+) (*n*=6 for CTIP2 and SATB2; *n*=3 for BRN1; 6-8 sections/mouse, with outliers across sections removed; mean ± SEM; Student's t test after outliers removal and Shapiro-Wilk test for normality; ⊗ indicate outliers).

All data collected and analyzed blind to genotype. In all panels, *Ddx3x*$^{+/+}$ (blue), and *Ddx3x*$^{+/-}$ (red). *p<0.05, **p<0.01, ***p<0.001; ns, non significant.

**Figure S10.** ***Ddx3x***$^{+/-}$ **mice have no other major cortical changes. A)** ***Ddx3x***$^{+/-}$ **mice have no changes in CTIP2+ ScPN and BRN1+ IT in M2.** Representative confocal images of coronal sections of M2 from *Ddx3x*$^{+/+}$ and *Ddx3x*$^{+/-}$ mice at P3, immunostained for CTIP2 (red) and BRN1 (green). **B) CTIP2+ ScPN are not misplaced in the M2 of** ***Ddx3x***$^{+/-}$ **mice.** Distribution of the percentage of cells (DAPI+) that are positive to CTIP2, across ten equally sized bins (*n* shown in legend; 6-8 sections/mouse, with outliers across sections removed; mean ± SEM; Two-way ANOVA). **C) BRN1+ IT are not misplaced in the M2 of** ***Ddx3x***$^{+/-}$ **mice.** Distribution of the percentage of cells (DAPI+) that are positive to BRN1, across ten equally sized bins (*n* shown in legend; 6-8 sections/mouse, with outliers across sections removed; mean ± SEM; Two-way ANOVA). **D)** ***Ddx3x***$^{+/-}$ **mice have no changes in SATB2+ CPN.** Representative confocal images of coronal sections of M1 and M2 motor cortices from *Ddx3x*$^{+/+}$ and *Ddx3x*$^{+/-}$ mice at P3 immunostained for SATB2 (red), a marker of UL and DL CPN. **E)** ***Ddx3x***$^{+/-}$ **mice have no changes in SATB2+ CPN.** Distribution of the percentage of cells (DAPI+) that are positive to SATB2, across ten equally sized bins (*n* shown in legend; 6-8 sections/mouse, with outliers across sections removed; mean ± SEM; Two-way ANOVA).

All data collected and analyzed blind to genotype. In all panels, *Ddx3x*$^{+/+}$ (blue), and *Ddx3x*$^{+/-}$ (red). ***p<0.001; ns, non significant.

**Supplemental Tables Legends**

**Table S1 (Related to Fig. 2-3). Cohorts used in the analysis of developmental milestones.** The table reports the tests ran on each of the four animal cohorts employed in the analysis of developmental milestones. Numeric values in the cells indicated the number of animals.

**Table S2 (Related to Fig. 4-5). Cohorts used in the analysis of adult behavior.** The table reports the tests ran on each of theanimal cohorts employed in the behavioral analyses. These include four cohorts for the adult (4-months old) animals, and two cohorts for the ageing (1-year old) animals. Numeric values in the cells indicated the number of animals.

**Table S3 (Related to Fig. 6). MRI data.** The table reports the absolute and relative volumes across the brain calculated from MRI data at postnatal day 3 (n=10/genotype).

**Supplemental Methods**

**Genotyping.** Tail biopsies were taken for identification purposes. Tail tissues were collected by dissecting ~0.2 cm of tail at P10. Genotyping was performed with Mouse Genotype (see **Fig. S1**). The *Ddx3x* alleles were identified using two sets of primers. Primers 4539 (5'- GATGCATACAACATCCTGAACC-3') and 4540 (5'-GCCCTGACTTCAAACCTCTTAG-3') yielded a 744 bp, a 899 bp, or a 412 bp amplicon for the wild type ($Ddx3x^{+}$), floxed ($Ddx3x^{flox}$), or Δexon2 ($Ddx3x^{-}$) alleles, respectively. Primers 4539 (5'-GATGCATACAACATCCTGAACC-3') and 4541 (5'-CTTGCTGTACTTCCTCCACTCTG-3') yielded a 515 bp or a 618 bp amplicon, for the $Ddx3x^{+}$ or $Ddx3x^{flox}$, respectively. The *Cre* transgene was amplified using primers 4572 (5'-AATGGTTTCCCGCAGAACCT-3') and 4573 (5'-GCATTGCTGTCACTTGGTCG-3'). To confirm male identity, the *Sry* locus was amplified using 4598 (5'-TGGCAGCCTGTTGATATCCC-3') and 4599 (5'-CTGAGGTGCTCCTGGTATGG-3').

**Synaptosomes.** Synaptosomes were isolated as described previously [1]. P21 $Ddx3x^{+/y}$, $Ddx3x^{+/+}$, and $Ddx3x^{+/-}$ mice were euthanized by cervical dislocation. The total cortices were rapidly dissected on ice and homogenized in ice-cold homogenizing buffer (0.32M sucrose, 1mM EDTA, 1mg/ml BSA, 5mM HEPES pH=7.4) in a glass Teflon® douncer, and centrifuged at 3,000g for 10 min at 4°C. The supernatants were recovered and centrifuged at 14,000g for 12 min at 4°C. The pellets containing synaptosomes were gently resuspended in Krebs-Ringer buffer (140mM NaCl, 5mM KCl, 5mM glucose, 1mM EDTA, 10mM HEPES pH=7.4), subjected to a density gradient upon addition of Percoll™ Plus (Sigma-Aldrich) (final 45% v/v/), and centrifuged at 14,000rpm for 2 min at 4°C to enrich the synaptosomes at the surface of the gradient. The synaptosomes were recovered, washed in Krebs Ringer buffer, and then centrifuged at 14,000rpm for 30 sec at 4°C. The pellets containing purified synaptosomes were gently resuspended in RIPA™ lysis buffer (ThermoFisher) containing protease, phosphatase, and RNase™ ribonuclease inhibitors (ThermoFisher using a polypropylene pellet pestle. The samples were incubated on ice for 5 min before centrifuging at 12,000g for 8 min at 4°C. The supernatants containing the synaptosomal lysate were recovered and used for immunoblots.

**Testing of developmental milestones.** Testing was performed as reported previously [2], using a battery of tests adapted from the Fox scale [3,4]. Details for each test are indicated below.

Physical Milestones. Body weight was taken between P1 and P28, and was measured in grams. Fur and skin development was monitored between P1 and P15 and scored as follows: 1 = bright red, 2 = nude/pink, 3 = nude/grey, 4 = grey/fuzzy on back and shoulders, 5 = black hair on back/grey fuzzy belly, 6 = black hair fully covering body. Ear development was monitored between P1 and P9, and was scored as follows: 0 = ear bud not detached from pinna, 1 = ear flap detached from pinna. Top and bottom tooth eruption was monitored between P7 and P18, and separately scored as follows: 0 = incisors not visible, 1 = incisors visible but not erupted, 2 = incisors fully erupted. Eye opening was monitored between P10 and P20, and scored as follows: 0 = eyes fully closed, 1 = eyes partially open, 2 = eyes fully open.

Sensory Milestones. Cliff aversion reflex was measured between P2 and P14. The reflex was assessed by placing the mouse on the edge of an opaque plexiglass platform 30 cm high and was scored as follows: 0 = fall off, 1 = stay, 2 = turn away. Additionally, the latency to turn away from the edge was recorded. Forelimb grasp reflex was measured between P4 and P13. The reflex was tested by stimulating the front paws with a small plastic zip tie. The responses were scored as follows: 0 = no response, 1 = paws folding in response to stimulation, 2 = paws grasping zip tie in response to stimulation, 3 = paws grasping strong enough to hold for at least 1 sec when hanging. Auditory startle response was measured between P6 and P18. The pups was exposed to an 80 dB click 30 cm above. The reflex was considered present (score 1) when the pup startled immediately after the click. Ear twitch reflex was assessed between P7 and P15 by stimulating each ear with the tip of a cotton swab pulled to form a filament. The reflex was considered present (score 1) when the ear moved in response to the stimulation. Vibrissa placing reflex (P5-15) and visual placing (P13-18) were tested by slowly lowering the pup by the tail onto a flat surface. The vibrissa response was considered present (score 1) when the pup reached out forward with its forepaws to prepare to place itself when its whiskers touched the surface. The visual response was considered present (score 1) when the pup reached out with its forepaws to prepare to place itself when it saw the surface close below, before the vibrissa touched the surface.

Motor Milestones. Surface righting reflex was measured between P2 and P13 by placing the pup supine on a flat surface and recording the latency to turn over to a prone position with all 4 paws on the ground. The negative geotaxis behavior (P2-16) was tested by placing the pup head down on a mesh covered platform at a 45 degree angle. Scoring was as follows: 0 = fall off, 1 = stay/move down/walk down, 2 = turn and stay, 3 =

turn, move up, and stay, 4 = turn and move up to the top. Air righting reflex (P8-19) was tested by dropping the pup from a supine position 30 cm above a padded surface and scored as follows: 0 = fall on back, 1 = fall onto side, 2 = fall onto all 4 paws, successfully righting themselves in the air. Locomotor activity in open field crossing was tested between P8 and P20 by recording the latency to exit a 13 cm diameter circle when placed in the center. Rod suspension skills were measured between P11 and P20 by recording the latency to stay suspended on a 3 mm rod, 30 cm above a padded surface. Hind limbs suspension skills were measured between P2 and P12 by recording the latency to stay suspended on the edge of a 50 mL falcon tube by the hind limbs. Grip strength was monitored between P5 and P14 by placing the mouse on a mesh grid and rotating the grid from a horizontal to vertical position. The angle of the grid when the mouse fell off was recorded.

**Handling and assessment of physical appearance and spontaneous activity prior to behavioral testing.** Adult and ageing animals were handled daily for one week prior to behavioral testing to assess general health, psychical appearance, and spontaneous activity, as detailed below.

On day 1, physical appearance was monitored in the housing room. Body weight (g) was taken and body length (cm) was measured from the base of the tail to the tip of the nose. Coat appearance was visually examined on a scale: 0 = ungroomed 0-25%, 1 = partial grooming 25-50%, 2 = semi-groomed 50-75%, 3 = groomed 75-100%. Skin color of footpads was visually measured on a scale: 0 = pink, 1 = purple, 2 = other. Whisker barbering was visually scored on a scale: 0 = normal, 1 = shortened/missing. Patches of missing fur on face and on body are visually measured separately on a scale: 0 = none, 1 = some, 2 = extensive. Wounding was visually measured on a scale: 0 = none, 1 = signs of previous wounding, 2 = slight wounds present, 3 = moderate wounds present, 4 = extensive wounds present.

On day 2, spontaneous general activity was monitored for 5 minutes by placing the animal in a 1000 mL jar in the behavioral suit. The following parameters were measured: grooming time (sec), number of rears, number of jumps, body position (scored on a scale: 0 = completely flat, 1 = lying on side, 2 = lying prone, 3 = sitting or standing, 4 = rearing on hind limbs, 5 = repeated vertical leaping), spontaneous activity (scored on a scale: 0 = none/resting, 1 = casual scratch/groom, slow movement, 2 = vigorous scratch/groom, moderate movement, 3 = vigorous rapid movement, 4 = extremely vigorous rapid movement), respiration rate (scored on a scale: 0 =

gasping, 1 = slow and shallow, 2 = normal, 3 = hyperventilation), tremor (scored on a scale: 0 = none, 1 = mild, 2 = marked), urination (scored on a scale: 0 = none, 1 = little, 2 = moderate, 3 = extensive), and defecation (number of fecal boli). After the 5 min observation, the mice were briskly transferred to an empty cage for observation by tipping the beaker over the cage. The following parameters were monitored: transfer arousal (scored on a scale: 0 = coma, 1 = prolonged freeze, slight movement, 2 = brief freeze, then active movement, 3 = no freeze, stretch attends, 4 = no freeze, immediate movement), eyelid closure (scored on a scale: 0 = eyes wide open, 1 = eyes half open, 2 = eyes closed), spontaneous piloerection (0 = none, 1 = coat standing on end), gait (0 = normal, 1 = fluid but abnormal, 2 = slow and halting, 3 = limited movement only, 4 = incapacitated), pelvic elevation (scored on a scale: 0 = markedly flattened, 1 = barely touches, 2 = normal, 3 = elevated), and tail elevation (scored on a scale: 0 = dragging, 1 = horizontally extended, 2 = less than 30° elevation, 3 = 30-60° elevation, 4 = 60-90° elevation).

On day 3, reflexes and reactions were observed in the housing room. Touch escape was measured by stroking a finger on the mouse's back, starting lightly and getting firmer, and scored on a scale: 0 = no response, 1 = escape response to firm stroke, 2 = escape response to light stroke, 3 = escape response to stroke approach. Trunk curl was measured by holding the mouse by the tail 30 cm above the surface and noting the presence or absence of a trunk curl. Bone tone was measured by compressing both sides of the mouse between the experimenter's thumb and index finger and scored on a scale: 0 = flaccid, 1 = slight resistance, 2 = extreme resistance. Startle reflex was measured by an 80 dB click 30 cm above the mouse and scored on a scale: 0 = none, 1 = ear/head twitch, 2 = jump. Limb withdrawal. Mice were restrained and sensibility was tested by pressing the rear paws using tweezers, and scored on a scale: 0 = no reaction; 0.5 = bending of toes, 1 = slight limb withdrawal; 2 = moderate limb withdrawal; 3 = fast limb withdrawal; 4 = vigorous and repetitive limb withdrawal.

On day 4, reflexes and reactions were observed in the housing room. The pinna reflex was tested by stimulating each ear with the tip of a cotton swap pulled to form a filament. The reflex was considered present when the ear moved in response to the stimulation. Salivation and lacrimation were observed separately and scored on a scale: 0 = none, 1 = mild, 2 = marked. Provoked biting was tested by inserting a 3 mm rod between the teeth at the side of the mouse's mouth and scored on a scale: 0 = none, 0.5 = nibble, 1 = biting. Vocalization was considered present if the mouse was audible squeaking during handling.

On day 5, motor reflexes were observed in the housing room. Forelimb placing was tested by slowly lowering the mouse by the tail onto a flat surface and scored on the following scale: 0 = none, 1 = reaching after nose contact, 2 = reaching after vibrissa contact, 3 = reaching after visually seeing surface, before vibrissa touch the surface. Hind limb placing was tested by having the mouse grip a vertical mesh grid with its forepaws while the experimenter held its tail so that its body is horizontal. When the tail was released, the response was scored on a scale: 0 = grabs but falls, 1 = grabs but hangs, 2 = grabs and pulls body onto the grid. Rod suspension skills were tested by recording the latency to stay suspended on a 3 mm rod, 30 cm above a padded surface (60 sec cutoff). Negative geotaxis was tested by placing the mouse head down on a mesh covered platform at a 45° angle and scored on a scale: 0 = fall off, 1 = stays, 2 = moves but does not turn 3 = turn and stays, 4 = turn and climbs up. The placing, hind limb placing, rod suspension, and negative geotaxis tasks were repeated three times in that order and the average of the three trials were used for analysis. The inverted screen task was tested by placing the mouse on a mesh platform, which was then inverted 180° 60 cm over a padded surface. The time that the mouse fell from an upside down position was recorded (60 sec cutoff).

**Open field test.** Overall activity and thigmotaxis were measured by placing the mice at the center of a 44x44 cm arena and allowed to freely explore for 30 min. The animals' activity was recorded by video tracking (Noldus EthoVision). The center zone of the arena was virtually defined as the area at least 7.6 cm away from any wall. The following measures were recorded: distance moved (cm), velocity (cm/s), time in center (s), frequency in center, and average latency to first enter center (s).

**Elevated plus maze test.** Mice were placed in the center of the arena facing one of the closed arms and allowed to freely explore for 10 min. The animals' activity was recorded by video tracking (Noldus EthoVision). The following measures were recorded: distance moved (cm), velocity (cm/s), frequency in closed arms, frequency in open arms, time in closed arms (s), time in open arms (s), and latency to first enter open arms (s).

**Accelerating rotarod.** Mice were tested on a motorized rod measuring 3 cm in diameter (Omnitech Electronics Inc) with gradual speed increase from 4 to 40 rpm over 5 min. Four month old mice were presented with three 5 min-trials per days, 1 hr apart, over two consecutive days. One year old mice were presented with four 5 min-trials per days, 1 hr apart, over three consecutive days. The latency to fall was recorded with Fusion software.

**Wire hanging test.** Mice, led from the base of the tail, were allowed to grasp a wooden rod placed 40 cm above the bench by both forepaws. The task consisted of a 5 sec-training session and two minutes after a 60 sec-testing session. The latency to fall and the distance traveled along the hanger were recorded.

**Balance beam test.** The balance beam test was performed as described before [5]. The beam consisted of a 50-cm long wooden square prism with 1.3 cm face placed horizontally 40 cm above the bench. Mice were allowed to freely walk on the beam for two sessions of 2 min each, separated by 4 hours. The number of slips and the distance covered were analyzed.

**Vertical pole test.** The vertical pole test was performed as described before [5]. A wooden pole wrapped in tape to facilitate walking was used. The test consisted of two consecutive training days and a testing day (3 trials each day). Mice were placed facing upwards just below the top of the pole and the time to complete the turn and the time to climb down the pole were measured. [5].

**Y maze spontaneous alternation test**. Mice were placed in the center of the Y-shaped maze and allowed to freely explore the maze for 15 min. The animals' activity was recorded by video tracking (Noldus EthoVision). The total number of choices, number of correct triad alternations (ABC), and the number of incorrect triad choices (ABB, ABA, or AAA) were calculated.

**Novel object recognition.** The paradigm was divided into four parts: handling and habituation, training, and testing. The apparatus used was an opaque white arena (44 x 44 cm). First, mice were habituated to the researcher (60 sec handling) and the arena (5 min of free exploration) for 5 consecutive days. 24 hrs after the last habituation day, training took place. Two identical objects were presented, which were placed in two opposite corners of the arena. Mice were placed in the center of the arena and allowed to freely explore for 10 min. After, 24 hrs later one of the two objects was replaced by a similar object, and mice were allowed to freely explore for 5 min. The animals' activity was recorded by video tracking (Noldus EthoVision). The following measures were recorded: distance moved (cm), velocity (cm/sec), cumulative duration interacting with the objects (sec), frequency to interact with the objects, and latency to first interact with the objects (sec). The time spent interacting with the objects were additionally recorded manually using stopwatches from recorded videos.

**Three-chamber social approach test.** Testing was performed using a three-chambered apparatus [6], as previously described [2]. On the first day, the target mice (4 month old C57BL/6J females) were trained by placing them under an inverted wire cup placed in the center of the front section of a side chamber, away from the door. Two trials (15 min, 2 hrs apart) were conducted. Target mice were observed for aggressive and hyperactive behaviors. Only docile target mice were used for testing. On the second day, subject mice were first habituated to the center chamber by placing them in the chamber with the side doors closed and allowing them to freely explore for 10 min. After a 90 min rest, subject mice were placed in the center chamber with the doors opened and allowed to freely explore the arena for 10 min. After another 90 min rest, mice were placed in the middle of the center chamber with the doors open. Each side chambers contained a weighted wire cup. On one side the cup was empty while on the other, the cup hosted the target mouse. Subject mice were allowed to freely explore the arena for 10 minutes. The side with the target mouse was counterbalanced between mice. Target mice were changed every trial and were employed only twice/day. In all sessions, the animals' activity was recorded by video tracking (Noldus EthoVision) and the following parameters were scored: distance moved (cm), velocity (cm/sec), cumulative time spent in each chamber (sec), and time sniffing the cup with the target mouse and the empty cup (sec). The time spent interacting with the cups was additionally recorded manually using stopwatches from recorded videos.

**Contextual and cued fear conditioning testing.** Testing was divided into three parts 24 hrs apart: training, contextual fear testing, and cued fear testing. A shock box with Med PC-IV software was used. On the first day (training), mice were placed in the shock box for a 12 min trial, and three tone-shock pairs were administered. The tones (20 sec, 80 dB, 2 kHz) were presented at 300, 440, and 580 sec marks. A mild foot shock (1 sec, 0.7 mA) was presented at the end of each 20 sec tone. Overhead white light was on, but no red light. White noise was on, and nitrile gloves and 70% EtOH were used. On the second day (contextual testing), mice were placed in the shock box for a 4 min trial where no tones or shocks were administered. The environment stayed exactly the same as training where overhead white light was on, but no red light, white noise was on, and nitrile gloves and 70% EtOH were used. On the third day (cued testing), mice were placed in a new shock box for a 7 min trial where two tones (20 s, 80 dB, 2 kHz) at the 120 and 260 sec marks were presented, but with no shock. The shock box was modified into a novel context by using a plastic floor insert and plastic curved wall insert to create a semicircle. Lemon extract was used to scent the room and Peroxiguard was used instead of 70% EtOH. Latex gloves were used instead of nitrile. Only red light was on and there was no white noise. In all sessions, the animals' activity was recorded by video tracking (Noldus EthoVision). The time spent freezing (1sec cutoff) was recorded.

**Supplemental references**

1 De Rubeis, S. *et al.* CYFIP1 coordinates mRNA translation and cytoskeleton remodeling to ensure proper dendritic spine formation. *Neuron* **79**, 1169-1182, doi:10.1016/j.neuron.2013.06.039 (2013).

2 Drapeau, E., Riad, M., Kajiwara, Y. & Buxbaum, J. D. Behavioral Phenotyping of an Improved Mouse Model of Phelan-McDermid Syndrome with a Complete Deletion of the Shank3 Gene. *eNeuro* **5**, doi:10.1523/ENEURO.0046-18.2018 (2018).

3 Fox, W. M. Reflex-ontogeny and behavioural development of the mouse. *Anim Behav* **13**, 234-241, doi:10.1016/0003-3472(65)90041-2 (1965).

4 Heyser, C. J. Assessment of developmental milestones in rodents. *Curr Protoc Neurosci* **Chapter 8**, Unit 8 18, doi:10.1002/0471142301.ns0818s25 (2004).

5 Creus-Muncunill, J. *et al.* Increased translation as a novel pathogenic mechanism in Huntington's disease. *Brain* **142**, 3158-3175, doi:10.1093/brain/awz230 (2019).

6 Nadler, J. J. *et al.* Automated apparatus for quantitation of social approach behaviors in mice. *Genes Brain Behav* **3**, 303-314, doi:10.1111/j.1601-183X.2004.00071.x (2004).
